# Human-specific lncRNAs contributed critically to human evolution by distinctly regulating gene expression

**DOI:** 10.1101/2023.05.31.543169

**Authors:** Jie Lin, Yujian Wen, Ji Tang, Xuecong Zhang, Huanlin Zhang, Hao Zhu

## Abstract

What genes and regulatory sequences critically differentiate modern humans from apes and archaic humans, which share highly similar genomes but show distinct phenotypes, has puzzled researchers for decades. Previous studies examined species-specific protein-coding genes and related regulatory sequences, revealing that birth, loss, and changes in these genes and sequences can drive speciation and evolution. However, investigations of species-specific lncRNA genes and related regulatory sequences, which regulate substantial genes, remain limited. We identified human-specific (HS) lncRNAs from GENCODE-annotated human lncRNAs, predicted their DNA binding domains (DBDs) and binding sites (DBSs), analyzed DBS sequences in modern humans (CEU, CHB, and YRI), archaic humans (Altai Neanderthals, Denisovans, and Vindija Neanderthals), and chimpanzees, and investigated how HS lncRNAs and their DBSs have influenced gene expression in archaic and modern humans. Our results suggest that these lncRNAs and DBSs have substantially reshaped gene expression. This reshaping has evolved continuously from archaic to modern humans, enabling humans to adapt to new environments and lifestyles, promoting brain evolution, and resulting in cross-population differences. The parallel analysis of gene expression in GTEx tissues by HS TFs and their DBSs indicates that HS lncRNAs have reshaped gene expression in the brain more significantly than HS TFs.

## 1. INTRODUCTION

The limited genomic but substantial phenotypic and behavioral differences between humans and other hominids make which sequence changes have critically driven human evolution an enduring puzzle (Anton et al., 2014; Pollen et al., 2023). Sequence changes include the birth and loss of genes, as well as the turnover and change of regulatory sequences (Albalat and Canestro, 2016; Kaessmann, 2010; Prud’homme et al., 2007). Studies have identified human-specific (HS) genes important for promoting human brain development (e.g., *NOTCH2NL* and *ASPM*) (Evans et al., 2005; Fiddes et al., 2018; Mekel-Bobrov et al., 2005; Pinson et al., 2022; Suzuki et al., 2018). However, gene-centric studies have limitations (Currat et al., 2006; Timpson et al., 2007; Yu et al., 2007), including that genes promoting brain enlargement may not critically determine other traits (e.g., bipedal walking); meanwhile, studies on species-specific lncRNA genes remain limited. LncRNA genes are a major class of new genes in mammals, and HS lncRNAs may have greatly influenced human evolution. On the other hand, multiple regulatory sequence changes important for human evolution have been identified, including new sequences (Liu et al., 2021), lost sequences (McLean et al., 2011), human accelerated regions (HARs) (Dong et al., 2016; Mangan et al., 2022; Prabhakar et al., 2006; Whalen and Pollard, 2022), and turnover of transcription factor (TF) DNA-binding domains (TF DBD, commonly called DNA-binding motifs) and DNA-binding sites (DBS) (Krieger et al., 2022; Otto et al., 2009; Zhang et al., 2023). However, no systematic investigation has been reported on lncRNA DBD and DBS.

Studies have generated multiple important findings about lncRNAs and epigenetic regulation. (a) About one-third of human lncRNAs are primate-specific (Derrien et al., 2012). (b) Species-specific lncRNAs exhibit distinct expression in tissues and organs of different species (Sarropoulos et al., 2019). (c) Many lncRNAs can bind to DNA sequences by forming RNA:DNA triplexes (Abu Almakarem et al., 2012), recruit histone and DNA modification enzymes to DBSs, and regulate transcription (Lee, 2009; Roadmap Epigenomics et al., 2015). (d) Approximately 40% of differentially expressed human genes result from interspecies epigenetic differences (Hernando-Herraez et al., 2015). Thus, besides HS TFs and their DBSs, HS lncRNAs and their DBSs also critically regulate gene expression human-specifically.

This study focuses on HS lncRNAs, HS lncRNA DBSs, and their impacts on human evolution by exploring multiple methods and resources. The first is RNA sequence search based on structure and sequence alignment to identify orthologues of lncRNA genes in genomes (Nawrocki, 2014; Nawrocki and Eddy, 2013). The second is the specific base-pairing rules that RNA and DNA sequences follow to form RNA:DNA triplexes (Abu Almakarem et al., 2012); these rules allow computational prediction of lncRNA DBDs and DBSs (Lin et al., 2019; Wen et al., 2022). The third is gene expression in organs and tissues, especially the Genotype-Tissue Expression (GTEx) project (GTEx Consortium, 2017), which provides data for examining and comparing gene expression regulation by HS lncRNAs and HS TFs. Finally, the genomes of modern humans, archaic humans, and multiple apes (especially chimpanzees) allow cross-species genomic and transcriptomic analysis (1000 Genomes Project Consortium et al., 2012; Chimpanzee Sequencing and Analysis Consortium, 2005; Meyer et al., 2012; Prufer et al., 2017; Pruüfer et al., 2013).

This study identified HS lncRNAs upon the orthologues of the GENCODE-annotated human lncRNA genes in 16 mammalian genomes, predicted their DBDs and DBSs in modern humans (CEU, CHB, and YRI, the three representative populations of Europeans, Asians, and Africans), archaic humans (Altai Neanderthals, Denisovans, and Vindija Neanderthals), and identified the counterparts of HS lncRNA DBSs in chimpanzees (Figure 1A). Our DBS prediction method combines a local alignment algorithm with RNA:DNA base-pairing rules to identify RNA:DNA triplexes, thus predicting DBDs and DBSs simultaneously. This outperforms DBS prediction using RNA:DNA base-pairing rules alone (Lin et al., 2019). Based on reliable recent data and methods, we also predicted HS TFs and their DBDs and DBSs. Cross-population and cross-species analyses based on HS TFs and their DBSs, HS lncRNAs and their DBSs, and substantial genomic and transcriptomic data suggest that HS lncRNAs and their DBSs have distinctly and continuously reshaped gene expression for adaptive human evolution.

**Figure 1.**
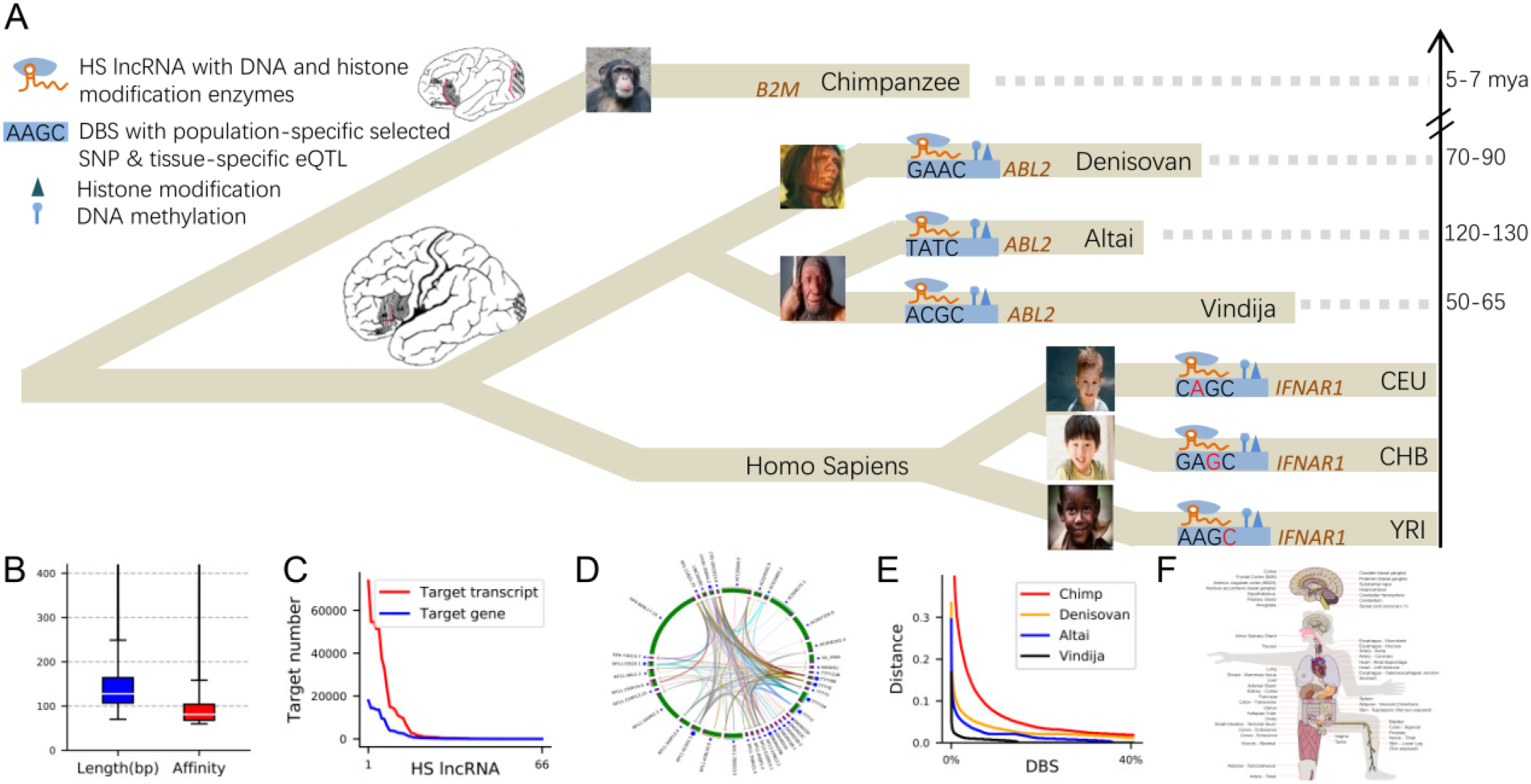
Study Overview. (A) The relationships between chimpanzees, the three archaic humans (Altai Neanderthals, Denisovans, and Vindija Neanderthals), and the three modern human populations, with dashed lines indicating the phylogenetic distances from modern humans based on related studies. Based on the left-top icons, the DBS in *B2M* lacks a counterpart in chimpanzees; the DBS in *ABL2* has great differences between archaic and modern humans; the DBS in *IRNAR1* is polymorphic in modern humans (red letters indicate tissue-specific eQTLs or population-specific mutations). (B) The mean length and affinity of strong DBSs. (C) Numbers of target genes and target transcripts of HS lncRNAs. (D) An illustrative figure showing the targeting relationships between HS lncRNAs. (E) The sequence distances of DBSs (the top 40%) from modern humans to chimpanzees and archaic humans. (F) An illustrative figure showing the impacts of HS lncRNA-target transcript on gene expression in GTEx tissues (see Supplementary Note 2 and Figure 3).

## 2. RESULTS

### 2.1. HS lncRNAs regulate diverse genes and transcripts

How many human lncRNAs are human-specific still lacks a precise estimate. Based on the orthologues of 13562 GENCODE-annotated human lncRNA genes (v18) in 16 mammalian genomes we searched using the *Infernal* program (which features in a combined sequence and structure alignment) (Lin et al., 2019; Nawrocki, 2014; Nawrocki et al., 2009), we identified 66 HS lncRNA genes that exist in humans but not in any other species (Supplementary Table 1). Using *the LongTarget* program, we predicted DBDs in exons of HS lncRNAs and their DBSs in the 5000 bp promoter regions of the 179128 Ensembl-annotated transcripts (release 79) (Lin et al., 2019). DBS prediction was validated using multiple methods and datasets (Supplementary Note 1). Predicted DBSs overlap well with experimentally identified DNA methylation and histone modification signals in multiple cell lines (https://genome.UCSC.edu), and overlap well with experimentally detected DNA binding sites of NEAT1, MALAT1, and MEG3 (Mondal et al., 2015; West et al., 2014). Many DBSs also co-localize with annotated cis-regulatory elements (cCREs) in promoter regions (https://genome.UCSC.edu). Specifically, we used CRISPR/Cas9 to delete multiple DBDs (100-200 bp) in several cell lines, performed RNA-seq before and after DBD knockout (KO), and analyzed the resulting differential gene expression. The seven DBD KO cases demonstrate that the |fold change| of target genes was significantly larger than the |fold change| of non-target genes (one-sided Mann-Whitney test, p=3.1e-72, 1.49e-114, 1.12e-206, 2.58e-09, 6.49e-41, 0.034, and 5.23e-07). Because DBD1’s DBSs vastly outnumber any other DBD’s DBSs, DBD1s not only regulate more targets than other DBDs but also are more reliable. Thus, this study analysed only DBD1s (simply called DBDs) and their DBSs.

To perform quantitative and comparative analysis, we identified HS lncRNAs and their DBSs in archaic humans and chimpanzees (Figure 1A), defined the *binding affinity* (simply called affinity) of a pair of DBD and DBS as the product of their length and identity (the percentage of paired RNA and DNA nucleotides), and computed DBSs’ sequence distances from humans to archaic humans and chimpanzees. LncRNA:DNA binding analysis at known imprinted genes suggests that affinity better characterizes DBSs than length (He et al., 2015). With affinity and distance, DBSs were classified into strong (affinity>=60) and weak (36<affinity<60) ones (as well as old and young ones based on when large sequence changes occurred). 105141 strong DBSs (mean length>147 bp) were identified in 96789 transcripts of 23396 genes, and 152836 weak DBSs were identified in 127898 transcripts of 33185 genes (Supplementary Table 2). Several HS lncRNAs (especially RP11-423H2.3) have abundant DBSs; only about 1.5% of target genes (0.6% of target transcripts) are human-specific; many targets have DBSs of multiple HS lncRNAs; many HS lncRNAs have multiple DBSs in a target gene; and HS lncRNAs themselves show complex targeting relationships (Figure 1BCD; Supplementary Note 2). These suggest that HS lncRNAs have significantly reshaped gene expression regulation.

### 2.2. Target genes with strong and weak DBS characterize adaptive evolution and human traits

Genes with significant sequence differences between humans and chimpanzees range from 1.24% to 5% (Britten, 2002; Ebersberger et al., 2002). However, substantial genes exhibit expression differences due to variations in regulatory sequences. It is therefore interesting to examine whether genes with strong and weak DBSs are enriched in different functions. We performed over-representation analysis (ORA) using the *g:Profiler* program, Gene Ontology (GO) database, and genes sorted by affinity (Supplementary Table 2). When choosing a cutoff to distinguish strong from weak DBSs, we found that the GO terms of the top 2000 protein-coding genes contain, but those of the top 1500 protein-coding genes lack, terms such as “hair follicle development” and “skin epidermis development” that critically differentiate humans from chimpanzees. We therefore performed ORA using the top 2000 and bottom 2000 protein-coding genes, respectively. In addition to shared GO terms (e.g., “behaviour”), the two gene sets have many specific ones. GO terms specific to the top 2000 genes include “renal system development”, and GO terms specific to the bottom 2000 genes include “cellular response to alcohol”, “response to temperature stimulus”, and “female pregnancy” (Supplementary Table 3; Supplementary Note 3).

Genes with strongest DBSs (affinity>=300) include *IFNAR1* and *NFATC1* (important for immune function), *KIF21B* and *NTSR1* (critical for neural development), *SLC2A11* and *SLC2A1* (involved in glucose usage), *BAIAP3* (a brain-specific angiogenesis inhibitor), *TAS1R3* (a receptor for the sweet taste response), and several primate-specific lncRNA genes (e.g., *CTD-3224I3.3* with high expression in the cerebellum, lung, and testis). DBSs in these genes underwent large sequence changes from chimpanzees and archaic humans to modern humans (Table 1; Supplementary Table 4).

**Table 1.** Genes with DBSs that have largest affinity values and mostly changed sequence distances (from modern humans to archaic humans and chimpanzees)

| Target gene | Annotation | Binding affinity | Mostly changed |
| --- | --- | --- | --- |
| IFNAR1 | i.e., Interferon Alpha And Beta Receptor Subunit 1. | 794 | C,D |
| NFATC1 | A TF that induces gene transcription during immune responses. | 736 | C |
| NFATC1 | A TF that induces gene transcription during immune responses. | 491 | C,A,D |
| ANKLE2 | Diseases associated with ANKLE2 include microcephaly. | 527 | C,D |
| SEMA4D | Regulating phosphatidylinositol 3-kinase signaling, neuron projection development, and phosphate metabolic process. | 495 | C,A,D |
| KIF21B | Essential for neuronal morphology, synapse function, and learning and memory. | 471 | C |
| ALDH3B2 | An aldehyde dehydrogenase for alcohol metabolism. | 444 | C |
| NTSR1 | A brain and gastrointestinal peptide that mediates functions of neurotensin (e.g., hypotension, hyperglycemia, hypothermia, and antinociception). | 402 | C,A,D |
| MC5R | A receptor for melanocyte-stimulating hormone and adrenocorticotrophic hormone. | 397 | C |
| THEG | Specifically expressed in the germ cells and involved in spermatogenesis. | 395 | C,D |
| HERC6 | In pathways including class I MHC-mediated antigen processing and presentation, and the innate immune system. | 369 | C,A,D |
| SLC2A11 | Facilitating glucose transporter. | 356 | C |
| NGEF | Playing a role in axon guidance regulating ephrin-induced growth cone collapse and dendritic spine morphogenesis. | 354 | C |
| SHC2 | Involved in the signal transduction pathways of neurotrophin-activated Trk receptors in cortical neurons. | 345 | C,D |
| BAIAP3 | Regulating behavior and food intake by controlling calcium-stimulated exocytosis of neurotransmitters, serotonin, and hormones like Insulin. | 321 | C |
| SLURP1 | A marker of late differentiation of the skin. | 319 | C |
| MLPH | Involved in melanosome transport. | 307 | C |
| TAS1R3 | Responding to the umami taste stimulus and recognizing diverse natural and synthetic sweeteners. | 304 | C, D |
| SLC2A1 | A major glucose transporter in the mammalian blood-brain barrier. | 356 | C |
| CTD-3224I3.3 | An lncRNA is highly expressed in the cerebellum, lung, and testis. | 312 | C,A,D |

'C', 'A', 'D', and 'V' indicate that the DBS has mostly changed sequence distances from modern humans to chimpanzees, Altai Neanderthals, Denisovans, and Vindija Neanderthals, respectively. *NFATC1* is displayed in two rows because the DBSs of SNORA59B and TTTY8/TTTY8B have affinity values of 736 and 491.

### 2.3 Genes intensively regulated by HS lncRNAs may have promoted human evolution

The human genome is approximately 99%, 99%, and 98% identical to the genomes of chimpanzees, bonobos, and gorillas (Chimpanzee Sequencing and Analysis Consortium, 2005; Prufer et al., 2012). However, the sequence variants that critically differentiate these species remain unclear. To assess whether regulation by HS lncRNAs is critical, we first examined whether DBSs of HS lncRNAs are human-specific or conserved across these species, and found that 97.81% of the 105141 strong DBSs have counterparts in chimpanzees. To further determine whether the remaining 2.2% are human-specific gains, we checked them using the UCSC Multiz Alignments of 100 Vertebrates, and found that they are present in the human genome but absent from the chimpanzee genome and all other aligned vertebrate genomes. Since most DBSs have chimpanzee counterparts, yet have evolved considerably from chimpanzees to archaic and modern humans, they share features of HARs. However, of the 312 HARs identified by the Zoonomia Project as important for 3D genome rewiring and neurodevelopment (Keough et al., 2023), only eight overlap 26 DBSs of 14 HS lncRNAs, suggesting that DBSs and HARs may contribute differently to human evolution.

HS lncRNAs’ DBSs may be generated before, together with, or after HS lncRNAs, and the first situation suggests that HS lncRNAs reshape gene expression via DBSs. We therefore identified counterparts of HS lncRNAs and their DBSs in Altai Neanderthals, Denisovans, and Vindija Neanderthals (Meyer et al., 2012; Prufer et al., 2017; Pru fer et al., 2013). While both classes of counterparts were identified in these archaic humans, sequence distances (defined as the distance per base throughout the manuscript) of HS lncRNAs from humans to archaic humans are smaller than those of DBSs, suggesting that many DBS sequences were generated before HS lncRNAs.

We then computed DBS sequence distances using two methods. The first is from the reconstructed human ancestor (downloaded from the EBI website) to chimpanzees, archaic humans, and modern humans. In this result, many DBS distances from the human ancestor to modern humans are shorter than those to archaic humans (Supplementary Note 3). The second is from the human genome to chimpanzees and archaic humans. This set of distances agrees better with the phylogenetic distances between chimpanzees, archaic humans, and modern humans (Figure 1).

We postulate that genes with large DBS distances may have contributed more to human evolution than genes with small DBS distances. To test this postulation, we sorted genes by DBS distances from humans to chimpanzees and to Alai Neanderthals (Supplementary Table 5), and applied ORA to genes with large and small DBS distances using the *g:Profiler* program and GO database (Benjamini-Hochberg FDR, threshold=0.05, 50<terms size<1000). First, we examined the top 25% and bottom 25% of genes to determine whether they are enriched for different GO terms. The result indicates that the top 25% of genes generate more enriched GO terms and also more human evolution-related GO terms (Figure 2A; Supplementary Table 6). Second, we examined the top and bottom genes that also critically differentiate humans from chimpanzees. Agoglia et al. fused human and chimpanzee induced pluripotent stem cells to generate tetraploid hybrid stem cells (hybrid iPS), differentiated these cells into neural organoids, and measured allele-specific expression (ASE) of genes in hybrid iPS (Agoglia et al., 2021). We selected the top 50% and bottom 50% of genes, intersected them with the 2891 genes with significant ASE (p-adj<0.01 and |LFC|>0.5) (Supplementary Table 5), and applied ORA to the four gene sets. The four results indicate that more GO terms, especially more human evolution-related GO terms, were generated by the top 50% of genes (Figure 2B; Supplementary Table 7). Using ASE genes with less significance (just p-adj<0.01) yielded similar results (Supplementary Note 3). These results support the postulation that genes with large DBS distances contribute more to human evolution than genes with small DBS distances. Different GO terms, and GO terms with different significance levels, generated based on DBS distances to chimpanzees and archaic humans (e.g., “sensory perception of sound”), provide information on the timing and content of reshaped transcriptional regulation’s influence on human evolution.

**Figure 2.**
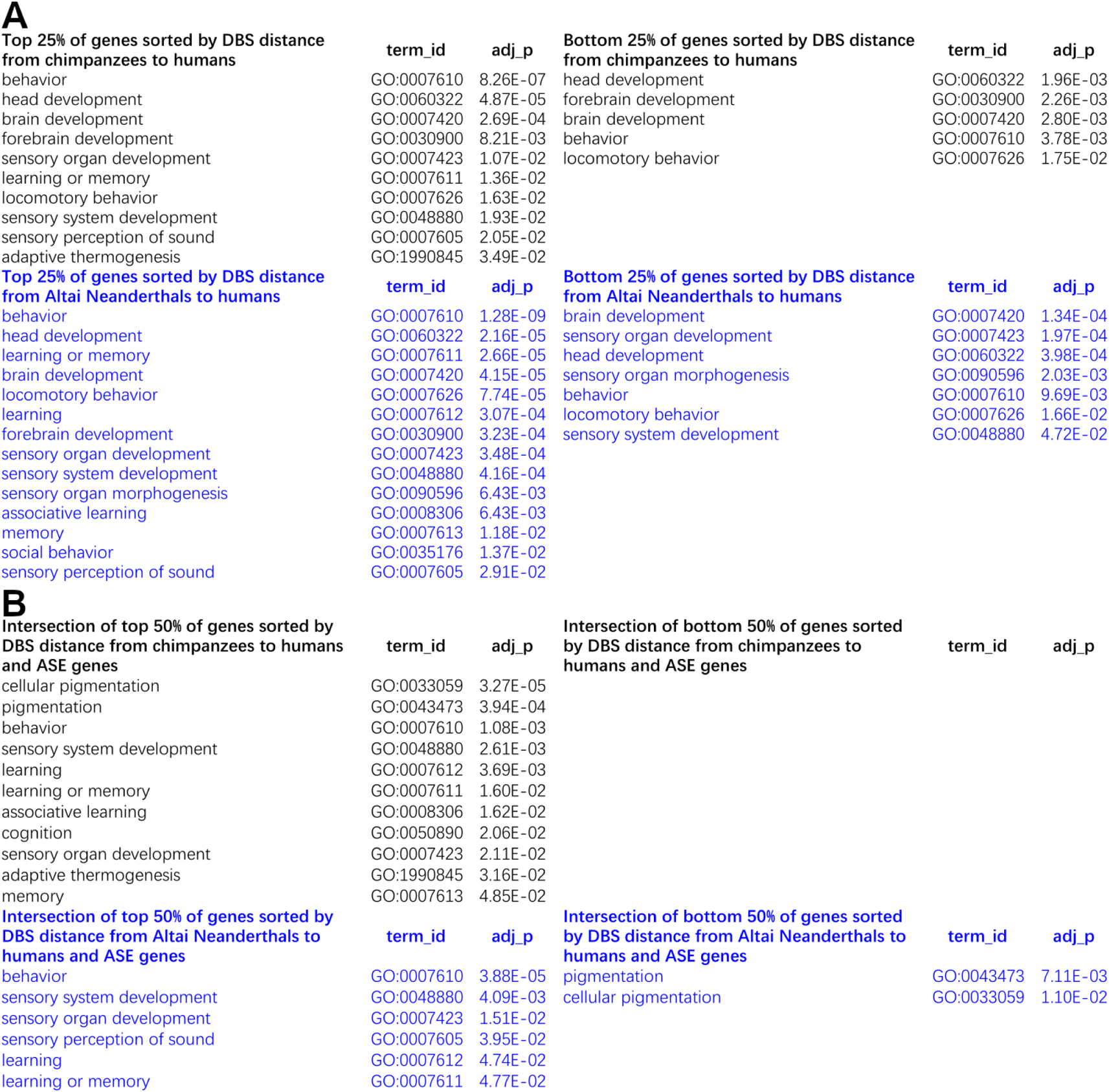
GO terms generated by different gene sets with large and small DBS distances from humans to chimpanzees and Altai Neanderthals. Shown are the presence and absence of GO terms highly related to human evolution. (A) Genes sorted by DBS distance from humans to chimpanzees and to Altai Neanderthals. Left: The top 25% of genes. Right: The bottom 25% of genes. (B) The intersections of the top 50% and bottom 50% of genes (with DBS distances from humans to chimpanzees and to Altai Neanderthals) and genes with significant ASE (p-adj<0.01 and |LFC|>0.5).

### 2.4. Regulation by HS lncRNAs shows cross-population differences

To reveal whether DBSs with large human-chimpanzee distances have further evolved in archaic and modern humans, we extracted those that (a) are in the top 20% of genes sorted by human-chimpanzee DBS distance (>0.037) and (b) also have a distance >0.037 to at least one archaic human. We label these DBSs using “A”, “D”, and “V” (e.g., “A,D” indicates the distances to both Altai Neanderthals and Denisovan >0.037), calculated SNP number therein (for SNPs with minimal allele frequency (MAF)≥0.1), and computed weighted Fst and Tajima’s D (1000 Genomes Project Consortium et al., 2012). SNP number per base, weighted Fst, and Tajima’s D reveal that many DBSs that undergo significant and continuous sequence change are also highly polymorphic and have been selected in specific modern humans (Supplementary Table 8). Many genes with these DBSs encode lncRNAs, and those encoding proteins include *SCTR*, *NCR*, *IFNAR1*, *NFATC1*, *TAS1R3*, *INS*, *ST3GAL4*, and *FN3KRP*, which are important for adaptive human evolution (Table 2). These suggest that the regulation of genes important for human evolution by HS lncRNAs has undergone continuous evolution.

**Table 2.**
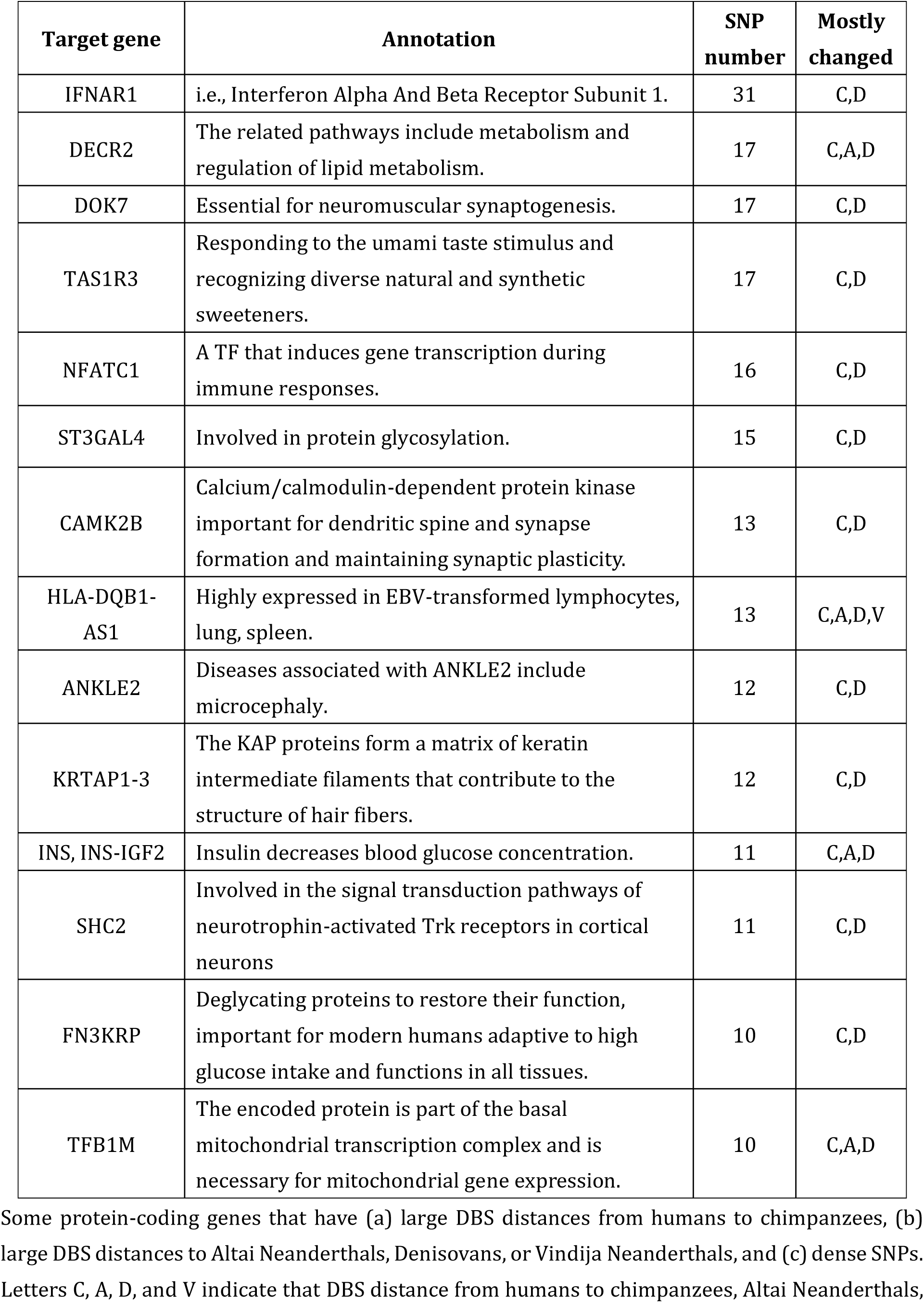
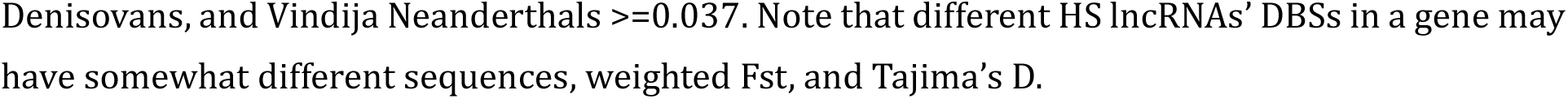
Genes with DBSs that are most polymorphic and have mostly changed sequence distances from humans to archaic humans and chimpanzees

| Target gene | Annotation | SNP number | Mostly changed |
| --- | --- | --- | --- |
| IFNAR1 | i.e., Interferon Alpha And Beta Receptor Subunit 1. | 31 | C,D |
| DECR2 | The related pathways include metabolism and regulation of lipid metabolism. | 17 | C,A,D |
| DOK7 | Essential for neuromuscular synaptogenesis. | 17 | C,D |
| TAS1R3 | Responding to the umami taste stimulus and recognizing diverse natural and synthetic sweeteners. | 17 | C,D |
| NFATC1 | A TF that induces gene transcription during immune responses. | 16 | C,D |
| ST3GAL4 | Involved in protein glycosylation. | 15 | C,D |
| CAMK2B | Calcium/calmodulin-dependent protein kinase important for dendritic spine and synapse formation and maintaining synaptic plasticity. | 13 | C,D |
| HLA-DQB1-AS1 | Highly expressed in EBV-transformed lymphocytes, lung, spleen. | 13 | C,A,D,V |
| ANKLE2 | Diseases associated with ANKLE2 include microcephaly. | 12 | C,D |
| KRTAP1-3 | The KAP proteins form a matrix of keratin intermediate filaments that contribute to the structure of hair fibers. | 12 | C,D |
| INS, INS-IGF2 | Insulin decreases blood glucose concentration. | 11 | C,A,D |
| SHC2 | Involved in the signal transduction pathways of neurotrophin-activated Trk receptors in cortical neurons | 11 | C,D |
| FN3KRP | Deglycating proteins to restore their function, important for modern humans adaptive to high glucose intake and functions in all tissues. | 10 | C,D |
| TFB1M | The encoded protein is part of the basal mitochondrial transcription complex and is necessary for mitochondrial gene expression. | 10 | C,A,D |

Some protein-coding genes that have (a) large DBS distances from humans to chimpanzees, (b) large DBS distances to Altai Neanderthals, Denisovans, or Vindija Neanderthals, and (c) dense SNPs. Letters C, A, D, and V indicate that DBS distance from humans to chimpanzees, Altai Neanderthals,
Denisovans, and Vindija Neanderthals $\geq 0.037$ . Note that different HS lncRNAs' DBSs in a gene may have somewhat different sequences, weighted $F_{st}$ , and Tajima's $D$ .

Genes involved in sugar metabolism are notable. A key feature of modern human diets is high sugar intake, which can cause non-enzymatic oxidation of proteins (i.e., glycation) and deactivate protein functions. Among proteins encoded by genes in this intersection, TAS1R3 recognizes diverse natural and synthetic sweeteners, Insulin decreases blood glucose concentration, ST3GAL4 regulates protein glycosylation, and FN3KRP deglycates proteins. These DBSs and target genes indicate the continuous evolution of transcriptional regulation with human- and population-specificity.

To examine whether SNPs in DBSs are neutral or indicate selection, we used multiple statistical tests with widely adopted parameters, including XP-CLR (Chen et al., 2010), iSAFE (Akbari et al., 2018), Tajima’s D (Tajima, 1989), Fay-Wu’s H (Fay and Wu, 2000), the fixation index (Fst) (Weir and Cockerham, 1984), and linkage disequilibrium (LD) (Slatkin, 2008), to detect selection signals in HS lncRNAs and strong DBSs in CEU, CHB, and YRI (Supplementary Note 4, 5; Supplementary Table 9, 10, 11). Selection signals were detected in several HS lncRNA genes in CEU and CHB but not in YRI, and selection signals were detected in more DBSs in CEU and CHB than in YRI. The same signals were often detected by multiple tests. These results agree with the reports that fewer selection signals are detected in YRI (Sabeti et al., 2007; Voight et al., 2006).

### 2.5. SNPs in DBSs exhibit cis-effects on gene expression

To verify that DBSs and SNPs influence gene expression, we analysed the GTEx data (GTEx Consortium, 2017). We identified expression quantitative trait loci (eQTLs) in DBSs using the widely adopted criteria of MAF≥0.1 and cis-effect size (ES)≥0.5. 1727 SNPs, with MAF≥0.1 in at least one population and |ES|≥0.5 in at least one tissue, were identified in DBSs in autosomal genes (Supplementary Table 12,13). These eQTLs include 372 “conserved” ones (i.e., also in DBSs in the three archaic humans) and 1020 “novel” ones (i.e., only in DBSs in modern humans). A notable eQTL with high derived allele frequencies (DAFs) across all three populations and a positive ES≥0.5 in 44 of 48 GTEx tissues is rs2246577 in the DBS in *FN3KRP,* whose encoded protein deglycates proteins to restore their function (Table 2). Many conserved eQTLs are expressed in brain tissues and exhibit high DAFs across all three modern populations. In contrast, many novel eQTLs are tissue-specific and exhibit population-specific DAFs (Supplementary Note 6).

Next, we performed two analyses to examine how eQTLs in DBSs influence gene expression. First, we computed the expression correlation between HS lncRNAs and target transcripts with DBSs having eQTLs in specific tissues. Most (94%) HS lncRNA-target transcript pairs showed significant expression correlation in tissues in which the eQTLs were identified (|Spearman’s rho|>0.3 and FDR<0.05). Second, we examined whether eQTLs are more enriched in DBSs than in Ensembl-annotated promoters by computing and comparing the eQTL density in the two classes of regions (promoters were used as the reference because they contain DBSs). The results indicate that eQTLs are more enriched in DBSs than in promoters (one-sided Mann-Whitney test, p=0.0) (Supplementary Note 6). Thus, population-specific SNPs and tissue-specific eQTLs, at the genome and transcriptome levels, respectively, suggest that DBSs enable HS lncRNAs to regulate transcription with population- and tissue-specific features.

### 2.6. HS lncRNAs promote brain evolution from archaic to modern humans

To further verify that SNPs in DBSs have cis-effects on gene expression, we examined the impact of HS lncRNA DBSs on gene expression in the GTEx tissues (GTEx Consortium, 2017). 40 autosomal HS lncRNAs are expressed in at least one tissue (median TPM>0.1), these HS lncRNAs and their target transcripts form 198876 pairs in all tissues, and 45% of pairs show a significant expression correlation in specific tissues (Spearman’s |rho|>0.3 and FDR<0.05). To assess the likelihood that these correlations could be generated by chance, we randomly sampled 10000 pairs of lncRNAs and protein-coding transcripts genome-wide and found that only 2.3% of pairs showed significant expression correlation (Spearman’s |rho|>0.3 and FDR<0.05). Moreover, a higher percentage (56%) of HS lncRNA-target transcript pairs with significant correlation was detected in at least one brain region, indicating more extensive gene expression regulation in the brain (Figure 3A).

**Figure 3.**
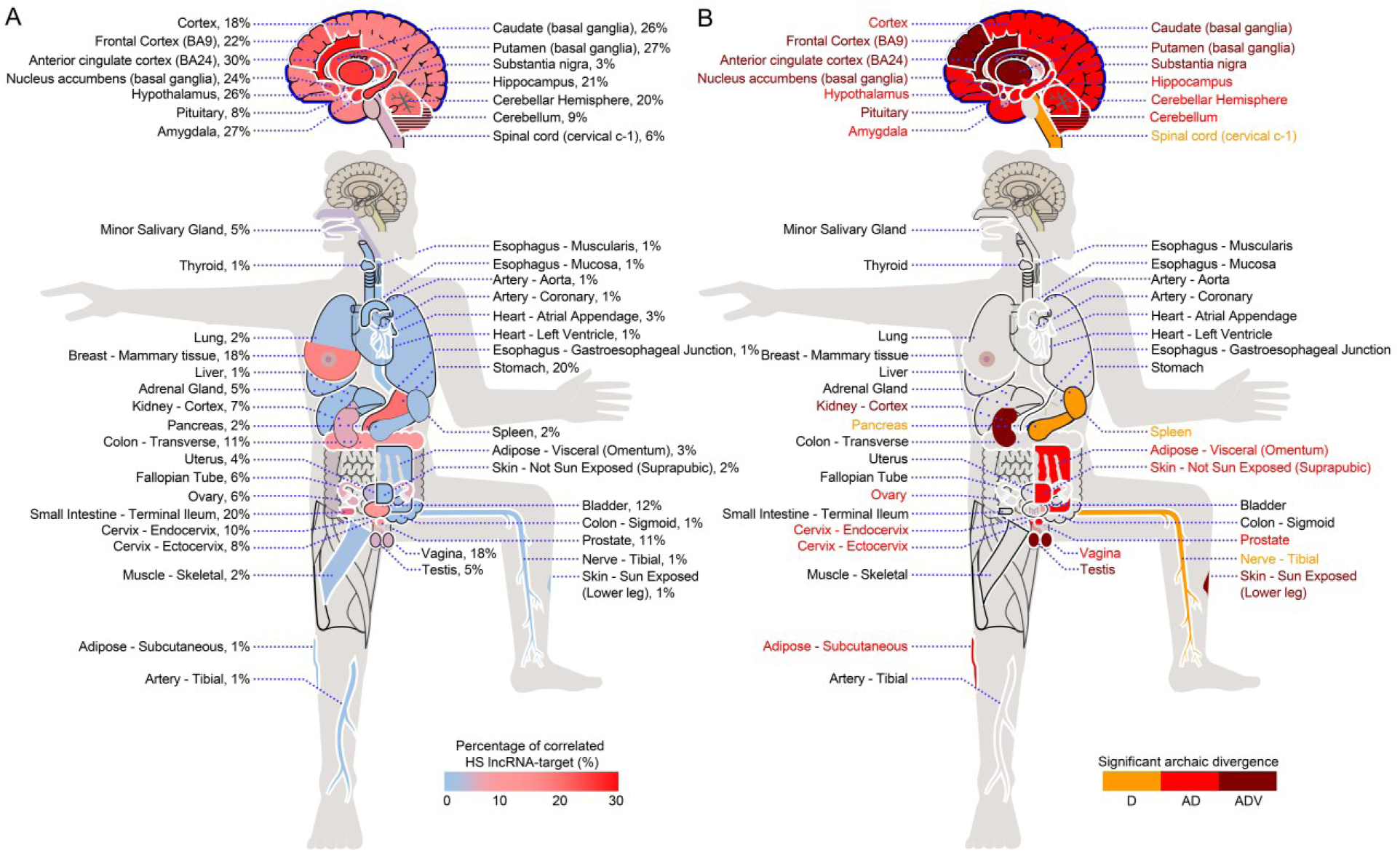
The impact of HS lncRNA-DBS interaction on gene expression in GTEx tissues and organs. (A) The distribution of the percentage of HS lncRNA-target transcript pairs with correlated expression across GTEx tissues and organs. Higher percentages of correlated pairs are in brain regions than in other tissues and organs. (B) The distribution of significantly changed DBSs (in terms of sequence distance) in HS lncRNA-target transcript pairs across GTEx tissues and organs between archaic and modern humans. Orange, red, and dark red indicate significant changes from Denisovans (D), Altai Neanderthals and Denisovans (AD), and all three archaic humans (ADV). DBSs in HS lncRNA-target transcript pairs with correlated expression in seven brain regions (in dark red) have changed significantly and consistently since the Altai Neanderthals, Denisovans, and Vindija Neanderthals (one-sided two-sample Kolmogorov-Smirnov test, significant changes determined by FDR <0.001).

To obtain more supporting evidence on the transcriptome level, we analyzed nine experimental datasets (Supplementary Note 7). First, we analyzed two datasets of epigenetic studies: one examining H3K27ac and H3K4me2 profiling in human, macaque, and mouse corticogenesis, and the other examining gene expression and H3K27ac modification in eight brain regions in humans and four other primates (Reilly et al., 2015; Xu et al., 2018). Compared with the genome-wide background, 84% and 73% of genes in the two datasets have DBSs for HS lncRNAs, indicating significantly higher enrichment in HS lncRNA targets (p=1.21e-21 and 1.2e-56, two-sided Fisher’s exact test). When chimpanzee gene expression was used as the control, 1851 genes showed human-specific transcriptomic differences in one or multiple brain regions, whereas only 240 genes showed chimpanzee-specific transcriptomic differences. Second, we analyzed two datasets of PsychENCODE studies: one examining spatiotemporally differentially expressed genes and spatiotemporally differentially methylated sites across 16 brain regions, and the other examining the spatiotemporal transcriptomic divergence between human and macaque brain development (Li et al., 2018; Zhu et al., 2018). In the first dataset, 65 HS lncRNAs are expressed in all 16 brain regions, 109 transcripts have spatiotemporally differentially methylated sites in their promoters, and 56 of the 109 transcripts have DBSs for HS lncRNAs. In the second dataset, 8951 genes show differential expression between the human and macaque brains, and 72% of differentially expressed genes in the human brain have DBSs for HS lncRNAs. Thus, both datasets show significant enrichment for HS lncRNA regulation. Third, three studies identified genes critically regulating cortical expansion (Florio et al., 2018; Johnson et al., 2018; Suzuki et al., 2018). Of the 40 reported protein-coding genes, 29 have DBSs of HS lncRNAs in promoter regions (Supplementary Table 14). Thus, these genes are enriched with DBSs of HS lncRNAs compared to the genome-wide background (p<0.01, two-sided Fisher’s exact test). Fourth, we analyzed two datasets of brain organoid studies. By establishing and comparing cerebral organoids between humans, chimpanzees, and macaques, Pollen et al. identified 261 human-specific gene expression changes (Pollen et al., 2019). By fusing human and chimpanzee iPS cells and differentiating the hybrid iPS cells into hybrid cortical spheroids, Agoglia et al. generated a panel of tetraploid human-chimpanzee hybrid iPS cells and identified thousands of genes with divergent expression between humans and chimpanzees (Agoglia et al., 2021). We found that 261 and 1102 genes in the two datasets are enriched for DBSs of HS lncRNAs compared to the genome-wide background (p=1.2e-16 and 3.4e-74).

Next, we examined the evolution of the impact of HS lncRNAs and their DBSs on gene expression using the GTEx data. For each DBS in an HS lncRNA-target transcript pair that shows correlated expression in a GTEx tissue, we computed the sequence distances of this DBS from the modern humans to the three archaic humans. Then, we compared the distribution of DBS sequence distances in each tissue with that in all tissues as the background (one-sided two-sample Kolmogorov-Smirnov test). DBSs in HS lncRNA-target transcript pairs with correlated expression in brain regions have significantly changed sequence distances since the Altai Neanderthals and Denisovans (Figure 3B), supporting that gene expression regulation by HS lncRNAs in the brain has undergone more significant evolution (Ponce de Leon et al., 2021).

To substantiate the above conclusion, we further examine whether HS lncRNAs have contributed more to human evolution than HS TFs. Based on the “hg38-panTro6” gene sets reported by Kirilenko et al. and the human TF lists reported by previous studies (Bahrami et al., 2015; Kirilenko et al., 2023; Lambert et al., 2018), five HS TFs were identified. We predicted their DBSs in the 5000 bp promoter regions of the same 179128 Ensembl-annotated transcripts (release 79) using the *FIMO* and *CellOracle* programs (Grant et al., 2011; Kamimoto et al., 2023), identified counterparts of these HS TF DBSs in archaic humans and chimpanzees, computed sequence distances of these DBSs from modern humans to archaic humans and chimpanzees (Supplementary Table 15), computed the Pearson correlation of HS TFs and their target transcripts across GTEx tissues, and computed sequence distances of DBSs in HS TF-target transcript pairs across GTEx tissues from modern humans to archaic humans. Highly correlated HS TF-target transcript pairs are distributed across many GTEx tissues, rather than being confined to the brain; also, significantly changed HS TF DBSs do not occur densely in the brain (Supplementary Note 8). These results further support that HS lncRNAs have promoted human brain evolution more than HS TFs.

### 2.7. HS lncRNAs mediate human-specific correlated gene expression in the brain

Finally, we further examined whether the gene expression pattern in the brain is human-specific. Based on the GTEx data from the human frontal cortex (BA9) and anterior cingulate cortex (BA24) (n=101 and n=83, respectively) (Consortium et al., 2017), and the gene expression data for the same brain regions in macaques (n=22 and n=25, respectively) (Zhu et al., 2018), we identified transcriptional regulation modules used the *eGRAM* program. In the two human brain regions, HS lncRNAs’ target genes form distinct modules, characterized by highly correlated gene expression (Pearson’s r>0.8) and enriched for KEGG pathways related to neurodevelopment (hypergeometric distribution test, FDR<0.05). In the same macaque brain regions, the orthologues of these human genes lack these features (Figure 4; Supplementary Note 9; Supplementary Table 16). The differences in gene modules suggest that the gene expression pattern in the human brain is highly human-specific and, to a great extent, can be attributed to the regulation by HS lncRNAs.

**Figure 4.**
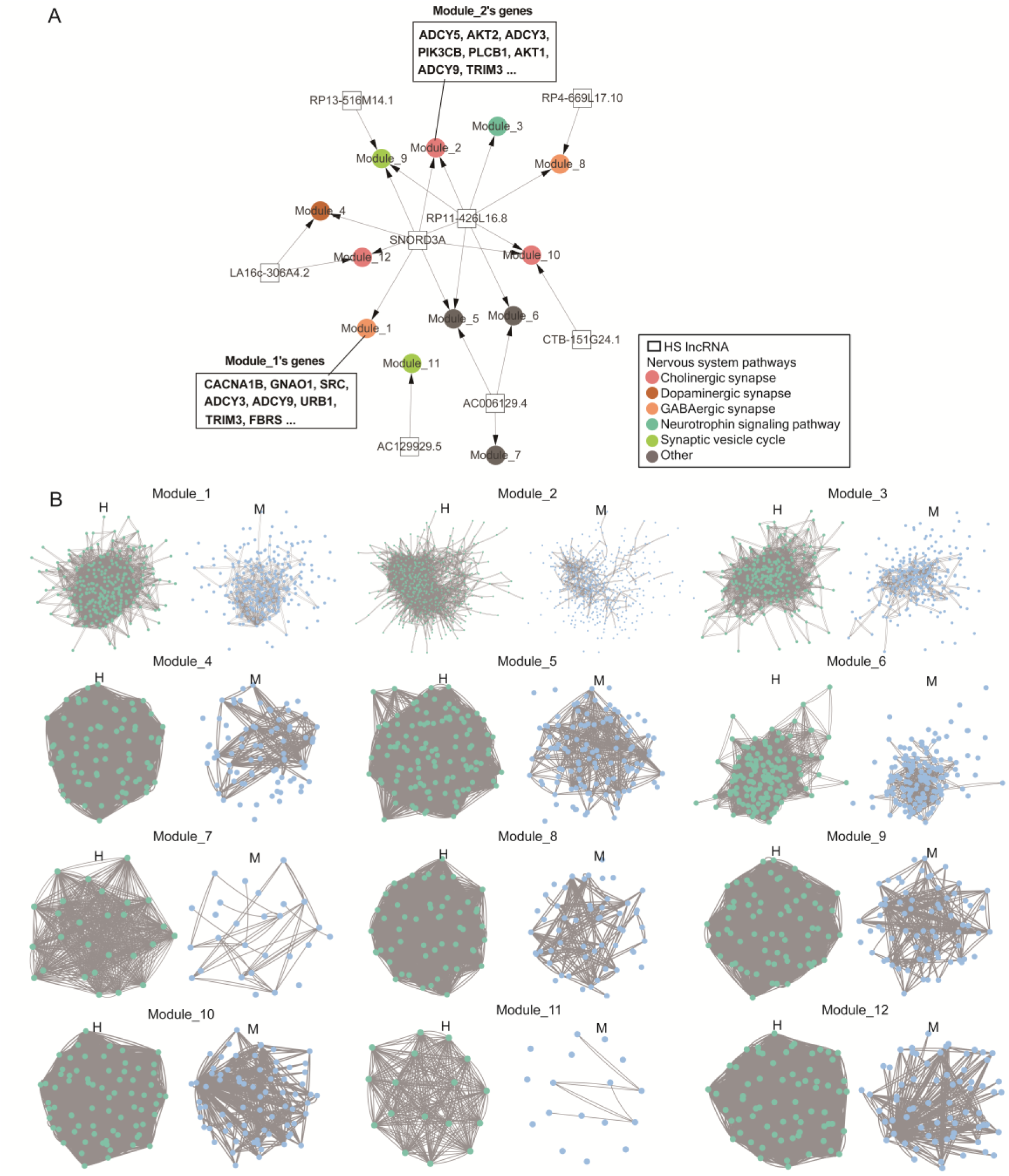
Human-specifically reshaped gene expression by HS lncRNAs in the frontal cortex (BA9). (A) Genes expressed in the human frontal cortex are enriched for HS lncRNAs’ target genes and neurodevelopment-related pathways. Squares, dots, and colors indicate HS lncRNAs, gene modules (Module_1 and Module_2 are illustrated), and enriched KEGG pathways, respectively. (B) Comparison of modules and genes in humans (indicated by H) and macaques (indicated by M). In each pair of modules, green and blue dots denote human genes and their orthologues, and lines between dots indicate correlated expression. Many orthologous genes in macaques (displayed at the corresponding positions) are not in the modules, and correlated expression is more prominent in humans than in macaques.

## 3. DISCUSSION

The limited genomic but substantial phenotypic and behavioral differences between modern humans, archaic humans, and apes make “what genomic differences critically determine modern humans” a profound question. This question has been addressed by many studies using various methods. Especially, studies examining protein-coding genes and HARs reported that HS protein-coding genes promote rapid neocortex expansion (Fiddes et al., 2018; Florio et al., 2018; Pinson et al., 2022) and that HARs are significantly enriched in genomic regions important for human-specific 3D genome organization (Keough et al., 2023). Many studies also examined lncRNAs in humans and mice. However, reports on HS lncRNAs and their DBSs have been rare. Similar to gene expression reshaping or rewiring by lineage-specific TFs and their DBSs (Johnson, 2017), gene expression can also be substantially rewired by lineage-specific lncRNAs and their DBSs. This study examined the postulation that HS lncRNAs and their DBSs have significantly reshaped gene expression during human evolution.

The nature of large-scale multi-omics data analysis means that the main results are unlikely to be generated by chance, although different parameters, thresholds, and significance levels can yield somewhat different results. Given the importance of DBS prediction, we used multiple methods, including DBD KO following differential gene expression analysis, to validate it. When predicting and analysing DBSs, we examined DBS in every transcript; thus, transcripts (including those in the brain) whose DBSs have had significant human-specific evolution can be identified. When running programs including *Infernal*, *LongTarget*, and g:*Profiler* and performing statistical tests that detect selection signals, we used the default parameters that suit most situations and the widely used significance levels. To compare the target genes of HS lncRNAs and their counterparts and orthologues in archaic humans and chimpanzees, we used multiple ORA-based methods and explored the reported ASE genes in tetraploid hybrid human-chimpanzee stem cells (Agoglia et al., 2021). Since weak DBSs reflect recent evolution but may be less reliable, we classified DBSs into strong/weak and old/young classes and found that different classes have distinct features. Understandably, not all detected signals reliably indicate positive selection; therefore, we also examined and described SNPs.

A few notes on several findings. First, relevant to and in line with the finding that Neanderthal-inherited sequences have measurable effects on gene expression in modern humans, but these effects are least detectable in the brain (McCoy et al., 2017), we find that HS lncRNAs and their DBSs have influenced gene expression in the brain more significantly than in other tissues. Second, the evolution of gene regulation by HS lncRNAs is rapid, and notable examples include those that enable humans to adapt to high-sugar intake. Third, regulated genes are enriched for different functions at different periods of human evolution, as evidenced by genes with young/old DBSs and with DBSs that contain “old” and “new” SNPs (e.g., DBSs containing “old” and “new” SNPs are in genes regulating neural development and glucose metabolism). Fourth, newly emerged regulations (possibly indicated by young weak DBSs) may have promoted recent human evolution.

Our findings also raise new questions. First, since many mouse lncRNAs are rodent-specific (Yue et al., 2014), two questions are how these lncRNAs specifically rewire gene expression in mice, and to what extent human- and mouse-specific lncRNAs cause differences in cross-species transcriptional regulation (Breschi et al., 2017; Hodge et al., 2019; Zhu et al., 2018). Second, whether mice and other mammals have such species-specific lncRNAs as RP11-423H2.3 that regulate the expression of substantial genes. Third, whether lineage- and species-specific lncRNAs would make many evolutionary novelties preordained.

## 4. MATERIAL AND METHODS

### 4.1. Data sources

The sequences of human (GRCh38/hg38, GRCh37/hg19) and chimpanzee genomes (panTro5) were obtained from the UCSC Genome Browser (http://genome.UCSC.edu). Three high-quality archaic human genomes were obtained from the Max Planck Institute for Evolutionary Anthropology (https://www.eva.mpg.de/genetics/index.html), which include an Altai Neanderthal that lived approximately 122 thousand years ago (kya) (Pru fer et al., 2013), a Denisovan (an Asian relative of Neanderthals) that lived approximately 72 kya (Meyer et al., 2012), and a Vindija Neanderthal that lived approximately 52 kya (Prufer et al., 2017). The ancestral sequences for the human genome (GRCh37) (which were generated upon the EPO alignments of six primate species human, chimpanzee, gorilla, orangutan, macaque, and marmoset) were downloaded from the EBI website (ftp://ftp.1000genomes.ebi.ac.uk/vol1/ftp/phase1/analysis_results/supporting/ancestral_alignments/). The SNP data of modern humans were obtained from the 1000 Genomes Project (phase I) (1000 Genomes Project Consortium et al., 2012). The three modern human populations are CEU (Utah residents with Northern and Western European ancestry), CHB (Han Chinese in Beijing, China), and YRI (Yoruba in Ibadan, Nigeria), which contain 100, 99, and 97 individuals, respectively. The genetic map files of modern humans were downloaded from the 1000 Genomes Project website (ftp://ftp-trace.ncbi.nih.gov/1000genomes/ftp/technical/working/20110106_recombination_hotspots/). The 1000 Genomes phased genotype data in VCF format were downloaded from the International Genome Sample Resource website (http://www.internationalgenome.org/). Annotated human genes and transcripts were obtained from the Ensembl website (http://www.Ensembl.org). FASTA-format nucleotide sequences were converted from VCFs using the *vcf-consensus* program in the *VCFtool*s package when necessary (Danecek et al., 2011). Data in the GeneCards Human Gene Database (www.genecards.org) were used to annotate genes and transcripts.

RNA-sequencing (RNA-seq) data (transcripts per million [TPM]) and eQTL data of human tissues were obtained from the Genotype-Tissue Expression Project (GTEx, v8) website (https://gtexportal.org/) (GTEx Consortium, 2017). Histone modification and DNA methylation signals in multiple cell lines, and ENCODE Candidate Cis-Regulatory Elements (cCRE), were obtained from the UCSC Genome Browser (http://genome.UCSC.edu). The genome-wide DNA-binding sites of lncRNA NEAT1, MALAT1, and MEG3 were obtained from experimental studies (Mondal et al., 2015; West et al., 2014). The predicted and experimentally detected DBSs in target genes of MALAT1, NEAT1, and MEG3 are given in the supplementary table (Supplementary Table 17).

### 4.2. Identifying HS lncRNA genes

We used the *Infernal* program (Nawrocki et al., 2009), which searches orthologous RNA sequences upon sequence and structure alignment, to identify orthologous exons in 16 mammalian genomes for each exon in each of the 13562 GENCODE (v18)-annotated human lncRNA genes (Lin et al., 2019). The 16 mammals were chimpanzee (CSAC 2.1.4/panTro4), macaque (BGI CR_1.0/rheMac3), marmoset (WUGSC 3.2/calJac3), tarsier (Broad/tarSyr1), mouse lemur (Broad/micMur1), tree shrew (Broad/tupBel1), mouse (GRCm38/mm10), rat (Baylor3.4/rn4, RGSC6.0/rn6), guinea pig (Broad/cavPor3), rabbit (Broad/oryCun2), dog (Broad CanFam3.1/canFam3), cow (Baylor Btau_4.6.1/bosTau7), elephant(Broad/loxAfr3), hedgehog (EriEur2.0/eriEur2), opossum (Broad/monDom5), and platypus (WUGSC 5.0.1/ornAna1) (http://genome.UCSC.edu). If the number of orthologous exons of a human lncRNA gene in a genome exceeded half the exon number of the human lncRNA gene, these orthologous exons were assumed to form an orthologous lncRNA gene. If a human lncRNA gene had no orthologous gene in all of the 16 mammals, it was assumed to be human-specific.

### 4.3. Identifying DBSs of HS lncRNAs

LncRNAs bind to DNA sequences by forming RNA:DNA triplexes. Each triplex comprises triplex-forming oligonucleotides (TFO) in the lncRNA and a triplex-targeting site (TTS) in the DNA sequence. We used the *LongTarget* program to predict HS lncRNAs’ DNA binding domains (DBD) and binding sites (DBS) with the default parameters (*Ruleset* = all, *T.T. penalty* = −1000, *CC penalty* = 0, *Offset* = 15, *Identity* ≥ 60, *Nt* ≥ 50) (Lin et al., 2019; Wen et al., 2022). The *LongTarget* program simultaneously predicts DBDs and DBSs, where a DBD comprises a set of densely overlapping TFOs and a DBS comprises a set of densely overlapping TTSs. For each HS lncRNA, we predicted its DBSs in the 5000 bp promoter regions (−3500 bp upstream and +1500 bp downstream the transcription start site) of the 179128 Ensembl-annotated transcripts (release 79). For each DBS, its binding affinity is the product of DBS length and the averaged *Identity* score of all TTSs (the *Identity* score is the percentage of paired nucleotides). Strong and weak DBSs were classified based on binding affinity ≥60 and <60. A transcript whose promoter region contains a strong DBS of an HS lncRNA was assumed to be a target transcript of the HS lncRNA, and the gene containing this transcript was assumed to be a target gene of the HS lncRNA. As the 1000 Genomes Project (phase I) data and the archaic human genomes are based on GRCh37/hg19, the DBS coordinates were converted from GRCh38/hg38 to GRCh37/hg19 using the *liftover* program from the UCSC Genome Browser (Kuhn et al., 2013).

### 4.4. Experimentally validating DBS prediction

A 157 bp sequence (chr17:80252565-80252721, hg19) containing the DBD of RP13-516M14.1, a 202 bp sequence (chr1:113392603-113392804, hg19) containing the DBD of RP11-426L16.8, and a 198 bp sequence (chr17:19460524-19460721, hg19) containing the DBD of SNORA59B, were knocked out in the HeLa cell line, RKO cell line, and SK-MES-1 cell line, respectively. Two sequences (chr1:156643524-156643684, chr10:52445649-52445740, hg38) containing the DBD of two wrongly transcribed noncoding sequences were knocked out in the HCT-116 and A549 cell lines, respectively. The seven knockouts were performed by UBIGENE, Guangzhou, China (http://www.ubigene.com) using CRISPR-U^TM^, a revised version of CRISPR/Cas9 technology. Before and after the seven DBD knockouts, RNA sequencing (RNA-seq) was performed by Novogene, Beijing, China (https://cn.novogene.com) and HaploX, Shenzhen, China ( https://www.haplox.cn/). The reads were aligned to the human GRCh38 genome using the *Hiasat2* program (Kim et al., 2019), and the resulting SAM files were converted to BAM files using *Samtools* (Li et al., 2009). The *Stringtie* program was used to quantify gene expression levels (Pertea et al., 2015). Fold change of gene expression was computed using the *edgeR* package (Robinson et al., 2010), and significant up- and down-regulation of target genes after DBD knockout was determined upon |log2(fold change)| > 1 with FDR < 0.1.

Genome-wide DBSs of NEAT1, MALAT1, and MEG3 were experimentally detected (Mondal et al., 2015; West et al., 2014). We also used these data to validate DBS prediction by predicting DBSs of the three lncRNAs and checking the overlap between predicted and experimentally detected DBSs (Supplementary Table 17).

### 4.5. Mapping DBSs of HS lncRNAs in the chimpanzee and archaic human genomes

We used the *liftover* program from the UCSC Genome Browser to map DBS loci from the human genome (hg38) to the chimpanzee genome (Pan_tro 3.0, panTro5). The mapping results were verified by inspecting the human-chimpanzee pairwise alignment in the UCSC Genome Browser. This initial screening identified 2248 DBSs (residing in 429 genes) that could not be mapped to the chimpanzee genome. To definitively determine whether these unmapped DBSs represent human-specific gains or chimpanzee-specific losses, we analyzed their sequences using the UCSC Multiz Alignments of 100 Vertebrates. This comparative genomics analysis confirmed that all 2248 DBSs are present in the human genome but are absent from the chimpanzee genome and all other aligned vertebrate genomes. Therefore, we classified these DBSs as human-specific gains.

We used *vcf-consensus* in the *VCFtool*s package to extract the DBSs of HS lncRNAs from the VCF files of Altai Neanderthals, Denisovans, and Vindija Neanderthals. The variant with the highest quality score was selected whenever multiple variant calls were observed at a given locus. The obtained DBS sequences in chimpanzees and three archaic humans are called counterparts of DBSs in these genomes.

### 4.6. Estimating sequence distances of DBSs between different genomes

We first aligned DBS sequences in the genomes of humans, chimpanzees, Altai Neanderthals, Denisovans, and Vindija Neanderthals using the *MAFFT*7 program to measure sequence distances from modern humans to chimpanzees and archaic humans (Katoh and Standley, 2013). We then computed sequence distances using the *dnadist* program with the Kimura 2-parameter model in the *PHYLIP* (3.6) package (http://evolution.genetics.washington.edu/phylip.html) and the *Tamura-Nei model* in the *MEGA7* package (Kumar et al., 2016). The two methods generated equivalent results. The largest distance between DBSs in humans and their chimpanzee counterparts is 5.383. Since 2248 DBSs in 429 human genes lack counterparts in chimpanzees, we assumed that these DBSs have a sequence distance of 10.0 between humans and chimpanzees.

We determined human ancestral sequences of DBSs using the human ancestor sequences from the EBI website, which were generated from the EPO alignments of six primate species. We used the above-mentioned methods to calculate the sequence distances DBSs from the human ancestor to chimpanzees, archaic humans, and modern humans. We found that when the human-chimpanzee ancestral sequence has the ancestral sequence (which means the inference of ancestral allele is of high confidence), DBS distances from the human ancestor to modern humans are larger than to archaic humans, but this situation accounts for only about 63.8%. For many DBSs, the distances from the human ancestor to modern humans are smaller than to archaic humans (especially Neanderthals and Denisovans) and even to chimpanzees. This defect may be caused by the absence of archaic humans in building the human ancestral sequence.

### 4.7. Detecting positive selection signals in HS lncRNA genes and DBSs

We used multiple tests to detect positive selection signals in HS lncRNA genes. First, we used the XP-CLR test (parameters = −w1 0.001 300 100 −p0 0.95, window size = 0.1 cM, grid size = 100 bp) to perform six pairwise genome-wide scans (i.e., CEU-CHB, CEU-YRI, CHB-CEU, CHB-YRI, YRI-CEU, and YRI-CHB) (Chen et al., 2010). The upper 1% of scores across the entire genome in each pairwise scan was 34.6 in the CEU-YRI scan, 16.8 in the CEU-CHB scan, 45.0 in the CHB-YRI scan, 26.9 in the CHB-CEU scan, 14.1 in the YRI-CEU scan, and 14.1 in the YRI-CHB scan. These scores were used as the thresholds of positive selection signals in these populations. Second, we used the *iSAFE* program to scan each genomic region containing an HS lncRNA gene and its 500-kb upstream and downstream sequences (Akbari et al., 2018). Strongly selected loci were detected only in CEU and CHB. Third, we used the *VCFtools* program to calculate Tajima’s D values for each HS lncRNA gene in CEU, CHB, and YRI (Danecek et al., 2011). The calculation was performed using a 1500-bp non-overlapping sliding window because the lengths of these genes exceed 1500 bp. To generate a background reference for assessing significant increases or decreases in Tajima’s D for HS lncRNA genes in a population, we calculated Tajima’s D across the whole genome using a sliding window of 1500 bp. As the values of Tajima’s D were compared with the background reference, significant D<0 and D>0 indicate positive (or directional) selection and balancing selection, respectively, rather than population demography dynamics (Tajima, 1989). Fourth, we used the integrated Fst to detect positive selection signals in HS lncRNA genes. The Fst of each HS lncRNA gene was computed for three comparisons, i.e., CEU-YRI, CHB-YRI, and CHB-CEU. Extreme Fst values of SNPs were detected in HS lncRNA genes in the comparisons of CEU-YRI and CHB-YRI. Since allele frequencies for different loci vary across a genome and genetic drift may have different effects at different loci, we used a sliding window of 1500 to compare Fst values of HS lncRNA genes with the genome-wide background. Extreme Fst values indicate positive selection. Finally, we applied linkage disequilibrium (LD) analysis to each HS lncRNA gene. We computed the pairwise LD (r²) in CEU, CHB, and YRI for common SNPs (with minimal allele frequency (MAF) ≥ 0.05 in at least one population) in HS lncRNA genes and DBSs using the *PLINK* program (Purcell et al., 2007). Significantly increased LD was detected in SNPs in HS lncRNA genes in CEU and CHB. The LD patterns were portrayed using the *Haploview* program (Barrett et al., 2005).

Next, we used the above tests to detect positive selection signals in DBSs. First, the 100 bp grid size of the XP-CLR test also allowed the detection of selection signals in DBSs. Second, we performed Tajima’s D and Fay-Wu’s H tests (Fay and Wu, 2000; Tajima, 1989). We calculated Tajima’s D values for each DBS in CEU, CHB, and YRI using the *VCFtools* program (Danecek et al., 2011), with a sliding window of 147 bp (the mean length of strong DBSs). To generate a background reference for judging the significant increase or decrease of Tajima’s D in a population, we calculated Tajima’s D values across the whole genome using the same sliding window. When Tajima’s D values were compared with the background reference, significant D<0 and D>0 indicate positive (or directional) selection and balancing selection, respectively. Fay-Wu’s H values were calculated similarly using the *VariScan* program (Vilella et al., 2005). Calculating Fay-Wu’s H demands the ancestral sequences as the outgroup. We extracted the ancestral sequences of DBSs from the human ancestral sequence, which was generated upon the EPO alignments of six primate species (human, chimpanzee, gorilla, orangutan, macaque, and marmoset) (ftp://ftp.1000genomes.ebi.ac.uk/vol1/ftp/phase1/analysis_results/supporting/ancestral_alignments/). Third, we computed the Fst to measure the frequency differences of alleles in DBSs between populations. For the CEU-YRI, CHB-YRI, and CHB-CEU pairwise comparisons, we used the revised *VCFtools* program to compute the weighted Fst for all SNPs in each DBS (Weir and Cockerham, 1984). Fourth, we integrated the weighted Fst values in the three populations into an “integrated Fst” which indicated whether the DBS locus was under selection in a certain population (Nielsen and Slatkin, 2013). We used sliding windows of 147 bp for comparing Fst values of DBSs with the genome-wide background. We empirically defined the upper 10% of integrated Fst scores across the entire genome as statistically significant. To determine positive selection more reliably, we used Tajima’s D and the integrated Fst to jointly determine if a DBS was under positive selection in a population. The thresholds that determined the upper 10% of Tajima’s D values across the entire genome in CEU, CHB, and YRI were −0.97, −0.96, and −0.97, respectively, and the threshold that determined the upper 10% of integrated Fst values across the entire genome in the three populations was 0.22. For example, a DBS was assumed to be under positive selection in CEU if (a) the DBS had a Tajima’s D<-0.97 in CEU and Tajima’s D>0.0 in the two other populations and (b) the DBS had an integrated Fst>0.22. Analogously, a DBS was assumed to be under positive selection in both CEU and CHB if the DBS had a Tajima’s D <-0.97 in CEU, <-0.96 in CHB, and >0.0 in YRI, and had an integrated Fst>0.22.

### 4.8. Functional enrichment analysis of genes

We used the *g:Profiler* program (with the parameter settings: Organism=Homo sapiens, Ordered query = No, Significance threshold = Benjamini-Hochberg FDR, User threshold = 0.05, 50<terms size<1000) and the Gene Ontology (GO) database to perform over-representation analysis (Raudvere et al., 2019). This analysis determines which pre-defined gene sets (GO terms) are more prevalent (over-represented) in a list of “interesting” genes than would be expected by chance. The lists of genes included genes with strong and weak DBSs, genes with large and small DBS distances from humans to chimpanzees, and genes with large and small DBS distances from humans to archaic humans. Strong DBSs have top affinity, and genes with weak DBSs not only have DBS affinity≤40 but also have DBSs of >=5 HS lncRNAs to help ensure that these genes are likely HS lncRNAs’ targets.

### 4.9. Analysing SNP frequencies in human populations

The frequencies of common SNPs (MAF ≥ 0.05) in DBSs across the three modern human populations were computed using the *VCFtools* package (Danecek et al., 2011). The ancestral/derived states of SNPs were inferred from the human ancestor sequences and were used to determine derived allele frequencies.

### 4.10. Analysing the cis-effect of SNPs in DBSs on target gene expression

SNPs with MAF >0.1 in DBSs in any of the three modern human populations and absolute values of cis-effect size >0.5 (FDR <0.05) in any of the GTEx tissues were examined for an influence on the expression of the target genes (GTEx Consortium, 2017). SNPs that are eQTLs in the GTEx tissues and have biased derived allele frequencies (DAF) in the three modern human populations were examined to estimate whether the eQTL is population-specific.

### 4.11. Examining the tissue-specific impact of HS lncRNA-regulated gene expression

First, we examined the expression of HS lncRNA genes across the GTEx tissues. HS lncRNA genes with a median TPM value >0.1 in a tissue were considered robustly expressed in that tissue. Upon this criterion, 40 HS lncRNA genes were expressed in at least one tissue and were used to examine the impact of HS lncRNA regulation on gene and transcript expression (other HS lncRNAs may function in the cytoplasm). Since an HS lncRNA gene may have multiple transcripts, we selected the transcript containing the predicted DBD and with the highest TPM as the representative transcript of the HS lncRNA. We calculated the pairwise Spearman’s correlation coefficient between the expression of an HS lncRNA (the representative transcript) and the expression of each of its target transcripts using the *scipy.stats.spearmanr* program in the *scipy* package. The expression of an HS lncRNA and a target transcript was considered to be significantly correlated if the |Spearman’s rho|>0.3, with Benjamini-Hochberg FDR<0.05. We examined the percentage distribution of correlated HS lncRNA-target transcript pairs across GTEx tissues and organs (Figure 3A).

We examined all GTEx tissues to determine which tissues may have exhibited significant changes in HS lncRNA-regulated gene expression from archaic to modern humans. GTEx data include both gene expression matrices and transcript expression matrices; we used the latter to examine changes in HS lncRNA-target transcript pairs from modern humans to archaic humans. If a pair of HS lncRNA and target transcript is robustly expressed in a tissue and their expression shows a significant correlation (|Spearman’s rho|>0.3, with Benjamini-Hochberg FDR<0.05) in the tissue, we computed the sequence distance of the HS lncRNA’s DBS in the transcript from the three archaic humans to modern humans. We compared the sequence distances of all DBSs in each tissue with the sequence distances of all DBSs in all GTEx tissues (as the background). A one-sided two-sample Kolmogorov-Smirnov test was used to examine whether the sequence distances of all DBSs in a specific tissue deviate from the background distribution (which reflects the “neutral evolution” of gene expression). For each tissue, if the Benjamini-Hochberg FDR was <0.001, the tissue was considered to have significantly altered gene expression regulated by the HS lncRNA. We used different colors to mark tissues with significantly changed gene expression regulation since Altai Neanderthals, Denisovans, and Vindija Neanderthals, and used “D”, “A.D.”, and “ADV” to indicate changes since Denisovans, since Altai Neanderthals and Denisovans, and since Altai Neanderthals, Denisovans, and Vindija Neanderthals, respectively (Figure 3B).

### 4.12. Examining enrichment of favored and hitchhiking mutations in DBSs

Using the deep learning network *DeepFavored*, which integrates multiple statistical tests for identifying favored mutations (Tang et al., 2022), we identified 13339 favored mutations and 244098 hitchhiking mutations in 17 human populations (Tang et al., 2023). In this study, we classified DBSs in two ways: into strong ones (affinity>60) and weak ones (36<affinity<60), and into old ones (Human-Chimp distance>0.034 AND Human-Altai Neanderthals distance=0), young ones (Human-Altai Neanderthals distance>0.034 OR Human-Denisovan distance>0.034), and others. We then examined the number of favored and hitchhiking mutations in each class of DBSs. The weak young DBSs have the largest proportion of favored and hitchhiking mutations.

### 4.13. The analysis of HS TFs and their DBSs

Kirilenko et al. recently identified orthologous genes in hundreds of placental mammals and birds and organized genes into pairwise datasets using humans and mice as the references (e.g., “hg38-panTro6”, “hg38-mm10”, and “mm10-hg38”) (Kirilenko et al., 2023). Based on the “many2zero” and “one2zero” gene lists (which contain 0 and 147 genes, respectively) in the hg38-panTro6 dataset, which indicate multiple human genes and a single human gene that have no orthologues in chimpanzees, we identified HS protein-coding genes. Further, based on three human TF lists reported by two studies and used in the *SCENIC* package (Bahrami et al., 2015; Lambert et al., 2018), we identified HS TFs. Using the JASPAR database and the CellOracle program (with default parameters), we predicted DBSs for HS TFs. Then, we repeated the steps of our HS lncRNA analyses (Supplementary Note 8; Supplementary Table 15), including computing sequence distances of DBSs and examining the impact of HS TF-target transcript pairs on gene expression in GTEx tissues and organs.

### 4.14. Identifying and analyzing transcriptional regulatory modules

Most clustering algorithms classify genes into disjoint modules based on expression correlation without considering regulatory relationships (Saelens et al., 2018). The *GRAM* program identifies gene regulatory modules based on correlation and TF-TFBS binding (Bar-Joseph et al., 2003). LncRNAs transcriptionally regulate genes based on lncRNA-DBS binding and correlate gene expression. We developed the *eGRAM* program to identify gene modules based on correlated expression, TF-TFBS interaction, and lncRNA-DBS interaction. In this study, we used *eGRAM* to identify gene modules in the same regions of the human and macaque brains and enriched KEGG pathways using reported RNA-seq datasets (n=101 and 83 for frontal cortex and n=22 and 25 for anterior cingulate cortex) (GTEx Consortium, 2017; Zhu et al., 2018). The default parameters, DBS binding affinity=60, Pearson correlation=0.5, module size=50, and FDR=0.01 (hypergeometric distribution test), were used. The key steps of the program are as follows. (a1) Identify each lncRNA’s correlated lncRNAs, which may form a set of co-regulators. (a2, optional) Identify each TF’s correlated TFs, which may form a set of co-regulators. (b1) Compute the correlation between each lncRNA and all genes. (b2, optional) Compute the correlation between each TF and all genes. (c1) Identify each lncRNA’s target genes. (c2, optional) Identify each TF’s target genes. (d1) Identify each lncRNA set’s target module upon correlation and targeting relationships. (d2, optional) Identify each TF set’s target module upon correlation and targeting relationships. (e) Check whether TFs’ modules contain lncRNAs’ targets and whether lncRNAs’ modules contain TFs’ targets, which reveal genes co-regulated by TFs and lncRNAs and genes independently regulated by TFs and ncRNAs. (f) Performs pathway enrichment analysis for all modules.

## AUTHOR CONTRIBUTIONS

H.Z. designed the study and drafted the manuscript. J.L. performed most of the data analyses. Y.W. and J.T. performed XP-CLR, iSAFE, and favored mutation analyses. J.L. and H.Z. developed the *eGRAM* program. All authors have read the manuscript and consent to its publication.

## ACKNOWLEDGEMENTS

This work was supported by the National Natural Science Foundation of China (31771456 to H.Z.) and the China Postdoctoral Science Foundation (2020M682788 to J.L.).

## DATA AND CODE AVAILABILITY

GENCODE human lncRNAs are available at the website (https://www.gencodegenes.org/human/). Human lncRNAs’ orthologues in 16 mammals, as well as the *LongTarget* program, are available at the *LongTarget*/*LongMan* website (http://www.gaemons.net/). The *eGRAM* code is available on the GitHub website (https://github.com/LinjieCodes/eGRAMv2R1). Other programs and tests are open-source and freely available. The RNA-seq data of the three HS lncRNAs before and after DBD KO have been deposited in the NCBI GEO database (https://www.ncbi.nlm.nih.gov/geo) (accession number GSE213231). The RNA-seq data of the other four cases of DBD KO have been deposited in the NCBI GEO database (https://www.ncbi.nlm.nih.gov/geo) (accession number is GSE229846). The data on favored and hitchhiking mutations are available from the PopTradeOff database or upon request. All other data are in Supplementary Tables.

## SUPPLEMENTARY DATA

Two supplementary files are available online with the manuscript. One is a PDF file containing 9 Supplementary Notes and 26 Supplementary Figures, and the other is an Excel file containing 17 Supplementary Tables.

