## Supplementary Notes for "Human-specific lncRNAs contributed critically to human evolution by distinctly regulating gene expression"

**Supplementary Note 1 – Supporting evidence for DBS prediction**

We used multiple methods and datasets to validate DBS prediction. First, since lncRNAs can recruit epigenetic regulatory enzymes to lncRNAs' DNA binding sites, we uploaded predicted DBSs of HS lncRNAs to the UCSC Genome Browser as a custom track, compared these DBSs with tracks such as "ENCODE DNA Methylation tracks" (Meissner et al., 2008) and "ENCODE Histone Modification tracks" (Ernst et al., 2011). We found that predicted DBSs overlap well with experimentally identified DNA methylation and histone modification signals in multiple cell lines (Supplementary Figure 1).

Second, since lncRNAs' DBSs can be detected genome-wide by experiments such as ChIRP-seq, we predicted the DBSs of the lncRNAs MALAT1, NEAT1, and MEG3 and compared predicted DBSs with the experimentally identified DNA-binding sites (Mondal et al., 2015; West et al., 2014). We found predicted DBSs agree well with experimentally identified DNA-binding sites (Supplementary Figure 2).

Third, we found that many DBSs also co-localize with ENCODE Candidate Cis-Regulatory Elements (cCREs) in promoter regions of genes (Supplementary Figure 3). cCREs are a subset of representative DNase hypersensitive sites across ENCODE and Roadmap Epigenomics samples supported by either histone modifications (H3K4me3 and H3K27ac) or CTCF-binding data.

Fourth, we used the CRISPR/Cas9 technique to knock out the sequences containing the DBD of seven lncRNAs (three HS lncRNAs and four wrongly transcribed long noncoding RNAs) in multiple cell lines and performed RNA-seq before and after the knockouts. The deleted sequences were just 100-200 bp, and gene expression analysis revealed that the |fold change| of target genes was significantly larger than the |fold change| of non-target genes (one-sided Mann-Whitney test, p = 3.1e-72, 1.49e-114, and 1.12e-206 for HS lncRNA RP13-516M14.1, RP11-426L16.8, and SNORA59B, and p = 2.58e-09, 6.49e-41, 0.034, and 5.23e-07 for the four wrongly transcribed lncRNAs in cancer cell lines) (Supplementary Figure 4). These results suggest that the deletion of DBD causes the changed expression of target genes (Supplementary Figure 5).

Fifth, the *LongTarget* program uses a variant of the Smith-Waterman algorithm to identify all local alignments in each lncRNA/DNA pair. Each alignment is an RNA:DNA triplex that consists of a TFO (triplex-forming oligonucleotides) and a TTS (triplex-targeting sites). A DBS is defined based on a set of densely overlapping TTSs, and a DBD is defined based on both TFOs and the DBS (He et al., 2015). *LongTarget* can more robustly and accurately identify DBS than some popular methods (Wen et al., 2022). To examine the likelihood that a local alignment of 147 bp (the average length of strong DBSs) can be generated by chance in a 5000 bp DNA sequence, we randomly simulated two sequences seqA and seqB (which were 5000 bp and 147 bp), and aligned seqB to seqA using the EMBOSS *Water* (https://www.ebi.ac.uk/Tools/psa/emboss_water/) local alignment program with alignment identity controlled to 60% (because the default *Identity* parameter of the *LongTarget* program was 60%). The equation $\mathrm{p} = 1 - \exp(-Kmne^{-\lambda S})$ was used to calculate a p-value to estimate the likelihood, in which $m$ and $n$ were the length of seqA and seqB, $K$ was a small adjusting constant (which was 0.1), λ was the normalization coefficient of the score (which was 0.3), and S was the alignment score. We repeated the simulation and calculation process 10000 times and determined that the maximal and minimal p-values were between 8.2e-19 and 1.5e-48. Thus, a DBS of 147 bp is extremely unlikely to be generated by chance. This result also supports that the changed expression of target genes is caused by the knockout of DBDs in the CRISPR/Cas9 experiments.

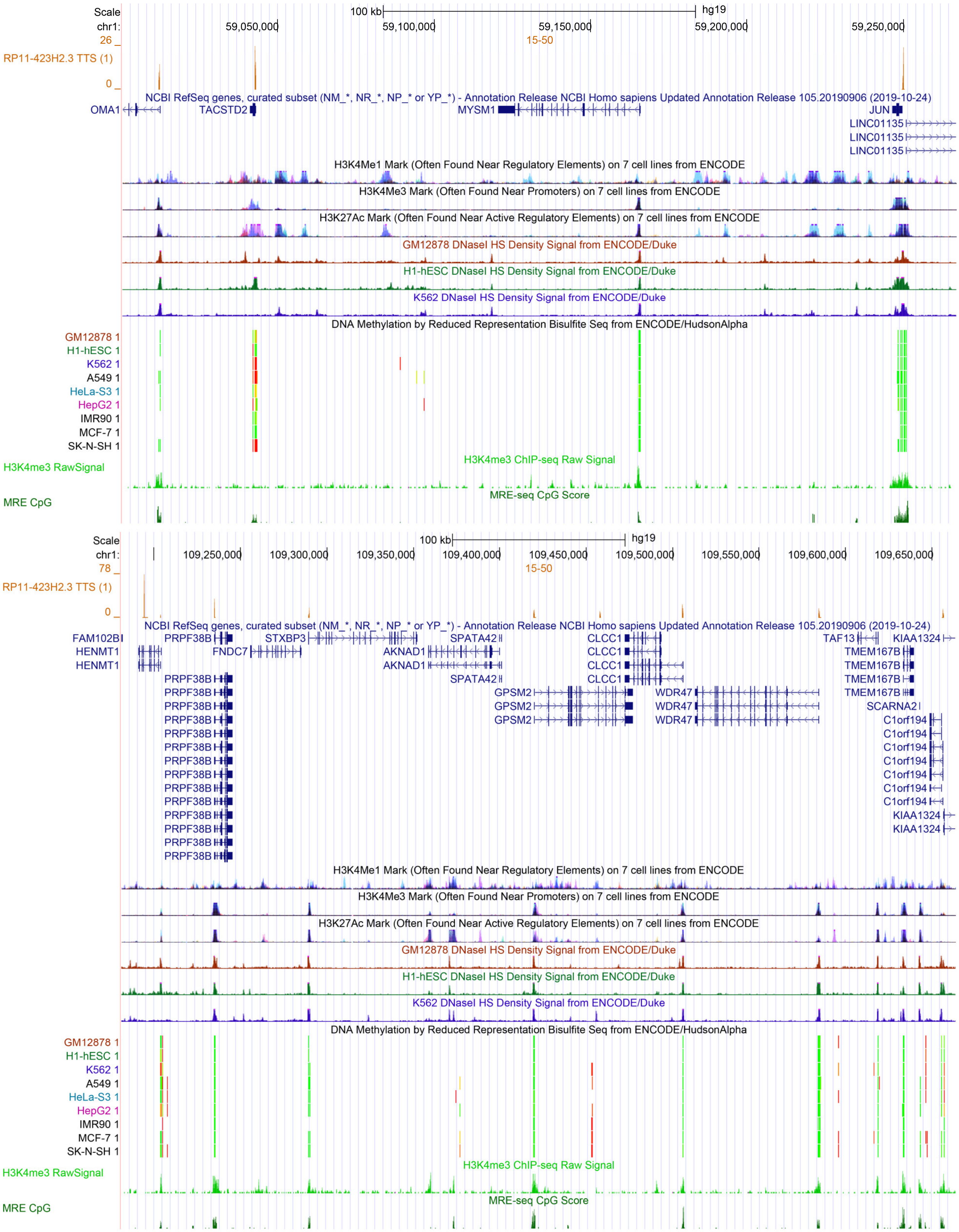


Supplementary Figure 1. DBSs of the HS lncRNA RP11-423H2.3 in two genomic regions. In the upper and lower panels that display two genomic regions, the tracks from top to bottom are DBSs (orange peaks), gene annotation, histone modification signals in cell lines, DNA methylation signals in cell lines, H3K4me3 RawSignal, and MRE CpG signals. DBSs overlap very well with DNA methylation and histone modification signals in multiple cell lines.

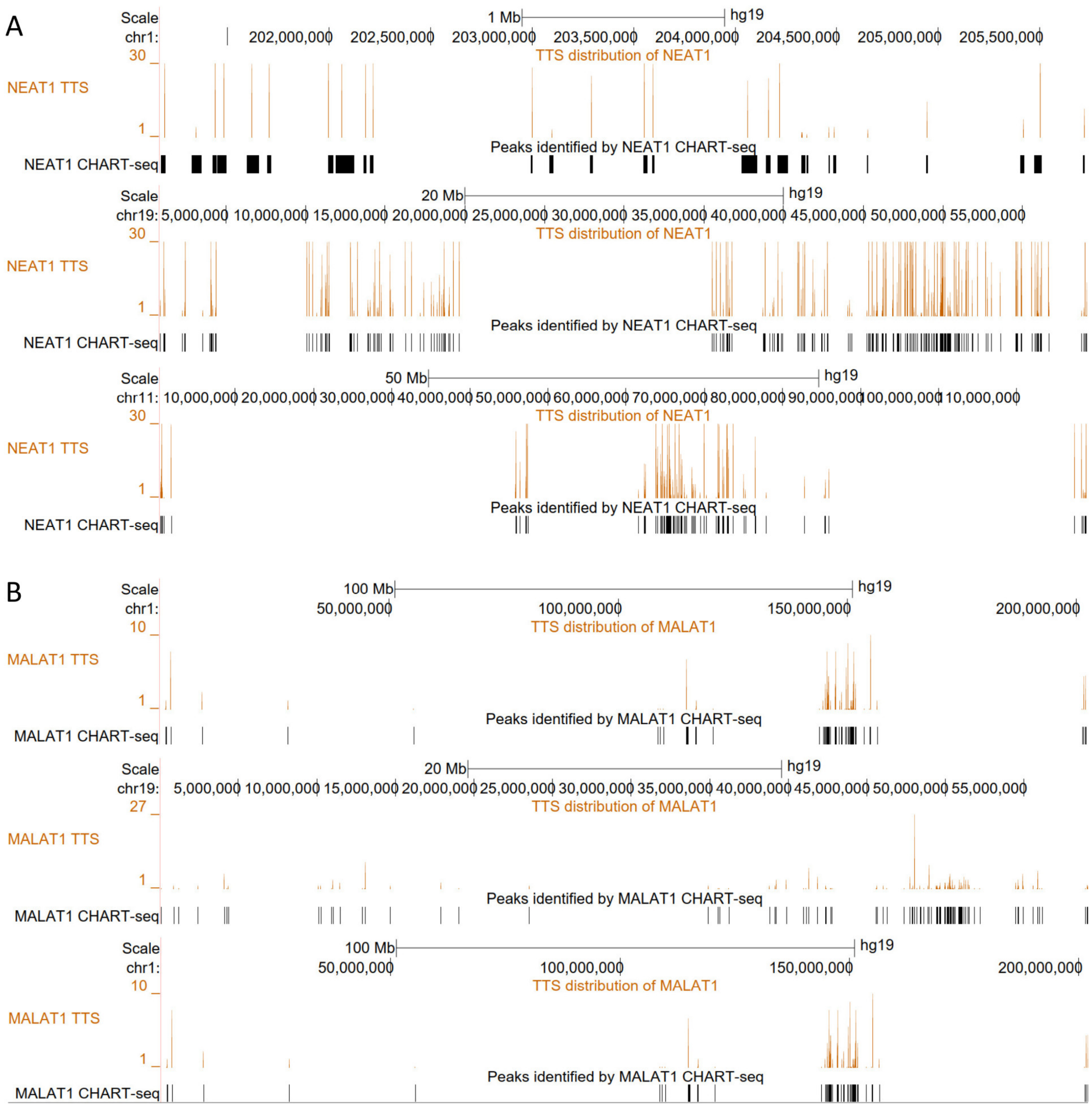


Supplementary Figure 2. Predicted DBSs and experimentally identified (by CHART-seq) DNA binding sites of NEAT1 and MALAT1 in two cell lines (West et al., 2014). DBSs were predicted using the DNA sequences of CHART-seq peaks. 99% and 87% of experimentally identified DNA binding sites of NEAT1 and MALAT1 overlap with predicted DBSs. (A) Predicted DBSs and experimentally identified DNA binding sites of NEAT1 in three genomic regions. (B) Predicted DBSs and experimentally identified DNA binding sites of MALAT1 in three genomic regions.

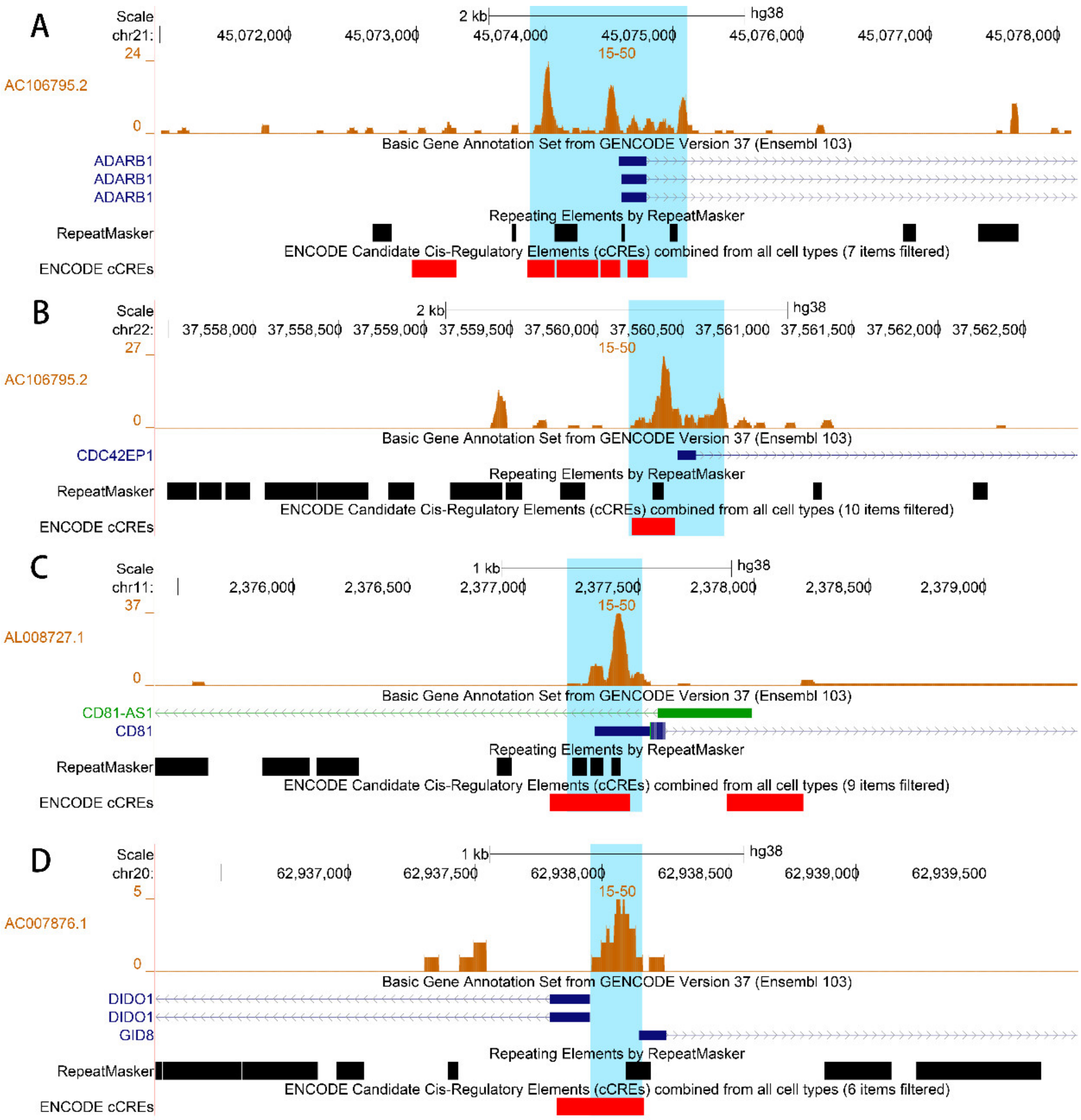


Supplementary Figure 3. Examples of co-localization of DBSs, TEs, and cCREs in the promoter regions of genes. (A) The DBSs of AC106795.2 in the promoter region of *ADARB1*. (B) The DBSs of AC106795.2 in the promoter region of *CDC42EP1*. (C) The DBS of AL008727.1 in the promoter region of *CD81*. (D) The DBS of AC007876.1 in the promoter region of *DIDO1* and *GID8*.

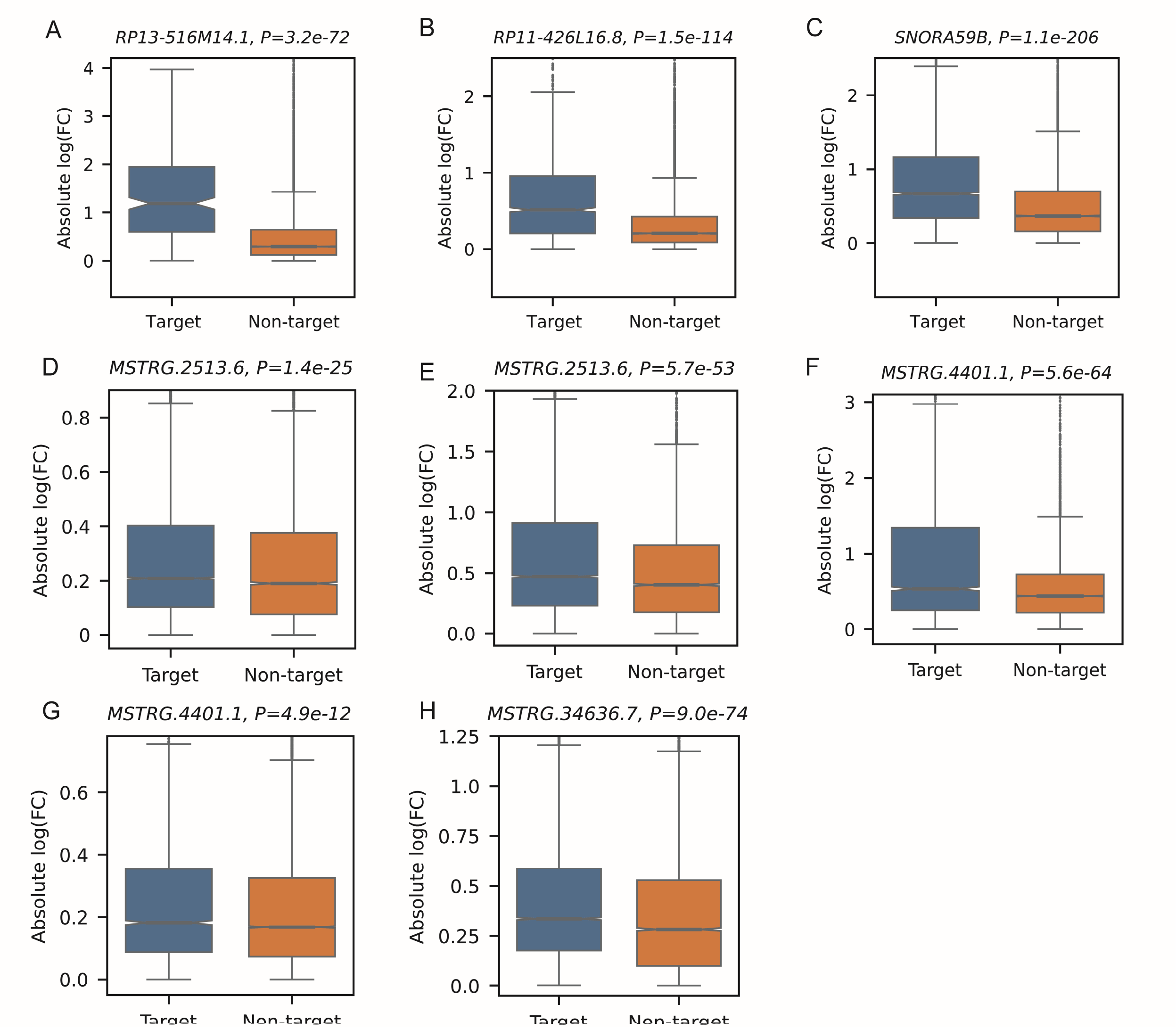


Supplementary Figure 4. The expression change of target genes was significantly larger than that of non-target genes after DBD knockout. The fold change of gene expression was computed using the edgeR package. The |fold change| distribution of target genes was compared with the |fold change| distribution of non-target genes (one-sided Mann-Whitney test). (A) The knockout of a 157 bp sequence (chr17:80252565-80252721 which contains the DBD of RP13-516M14.1, in the HeLa cell line. (B) The knockout of a 202 bp sequence (chr1:113392603-113392804 which contains the DBD of RP11-426L16.8, in the RKO cell line. (C) The knockout of a 198 bp sequence (chr17:19460524-19460721 which contains the DBD of SNORA59B, in the SK-MES-1 cell line. (D-E) The knockout of the DBD of a wrongly transcribed long noncoding RNA (chr1:156641670-156661464) in the A549 cell line and the HCT116 cell line. (F-G) The knockout of the DBD of a wrongly transcribed long noncoding RNA (chr10:52443915-52455313) in the A549 cell line and the HCT116 cell line. These wrongly transcribed long noncoding RNAs are labeled as “MSTRG” transcripts by the *Stringtie* package. (H) The knockout of the DBD of a third wrongly transcribed long noncoding RNA in the HCT116 cell line.

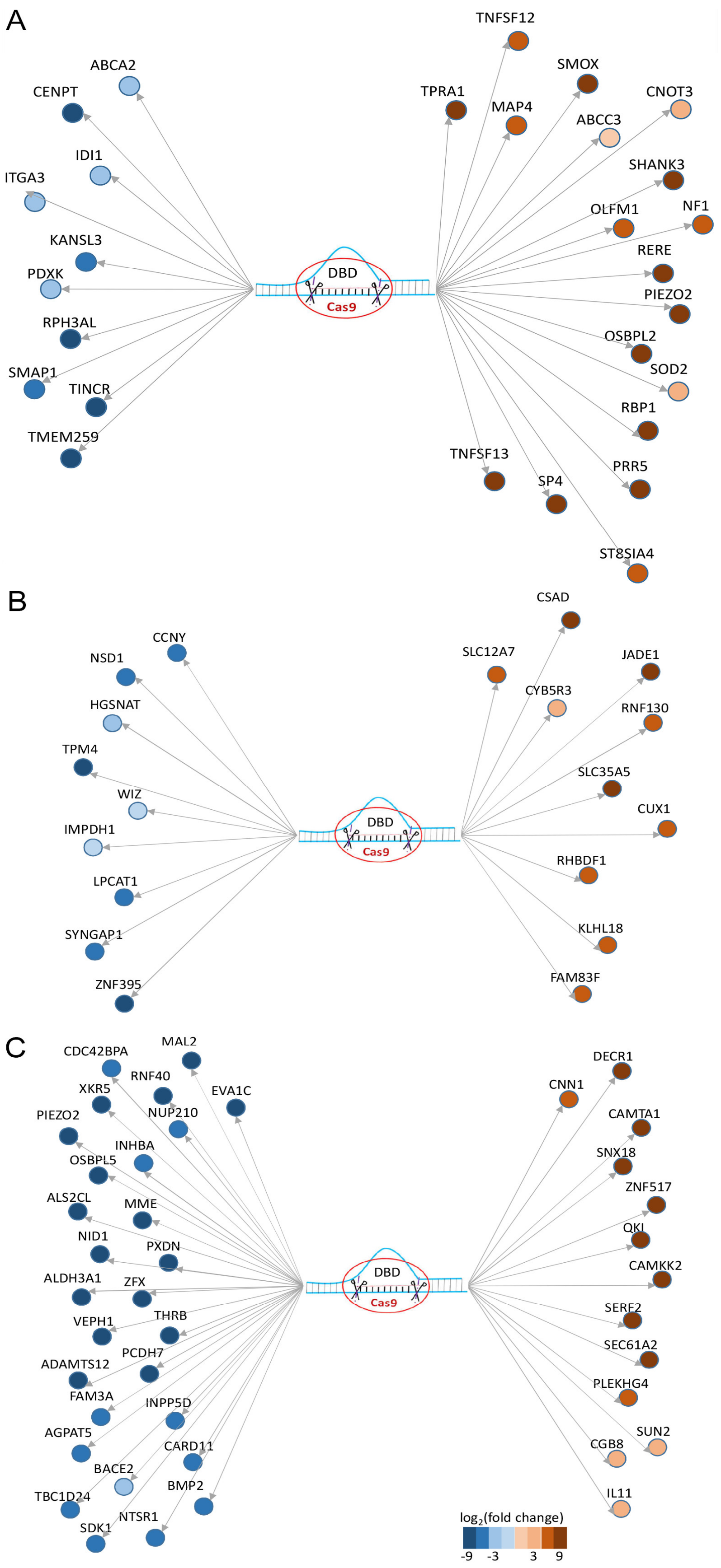


Supplementary Figure 5. Significant up- and down-regulation (|log2 (fold change)| > 1, FDR < 0.1) of target genes after DBD knockout. (A) RP13-516M14.1. (B) RP11-426L16.8. (C) SNORA59B.

**Supplementary Note 2 – Features of HS lncRNA-mediated gene expression regulation**

HS lncRNA-mediated gene expression regulation shows multiple features. First, HS lncRNAs themselves form complex targeting relationships (Supplementary Figure 6), suggesting networks and cascades of regulation. Second, some DBSs are human-specific sequences (Supplementary Figure 7), suggesting that human-specific sequences were explored by HS lncRNAs for regulating gene expression. Third, many genes and transcripts have DBSs for multiple HS lncRNAs (Supplementary Figure 8), suggesting co-regulation of functionally related genes. Fourth, certain genes and transcripts have multiple DBSs for one HS lncRNA, suggesting tissue-specific regulation. Fifth, selection signals were detected in some DBSs in specific populations (Supplementary Figure 9). Sixth, generally, HS lncRNAs on the Y chromosome have longer DBSs than HS lncRNAs on the autosomes (Supplementary Figure 10). Finally, SNPs in the DBSs of an HS lncRNA in multiple genes on a chromosome show LD, suggesting an association between these DBSs (Supplementary Figure 11).

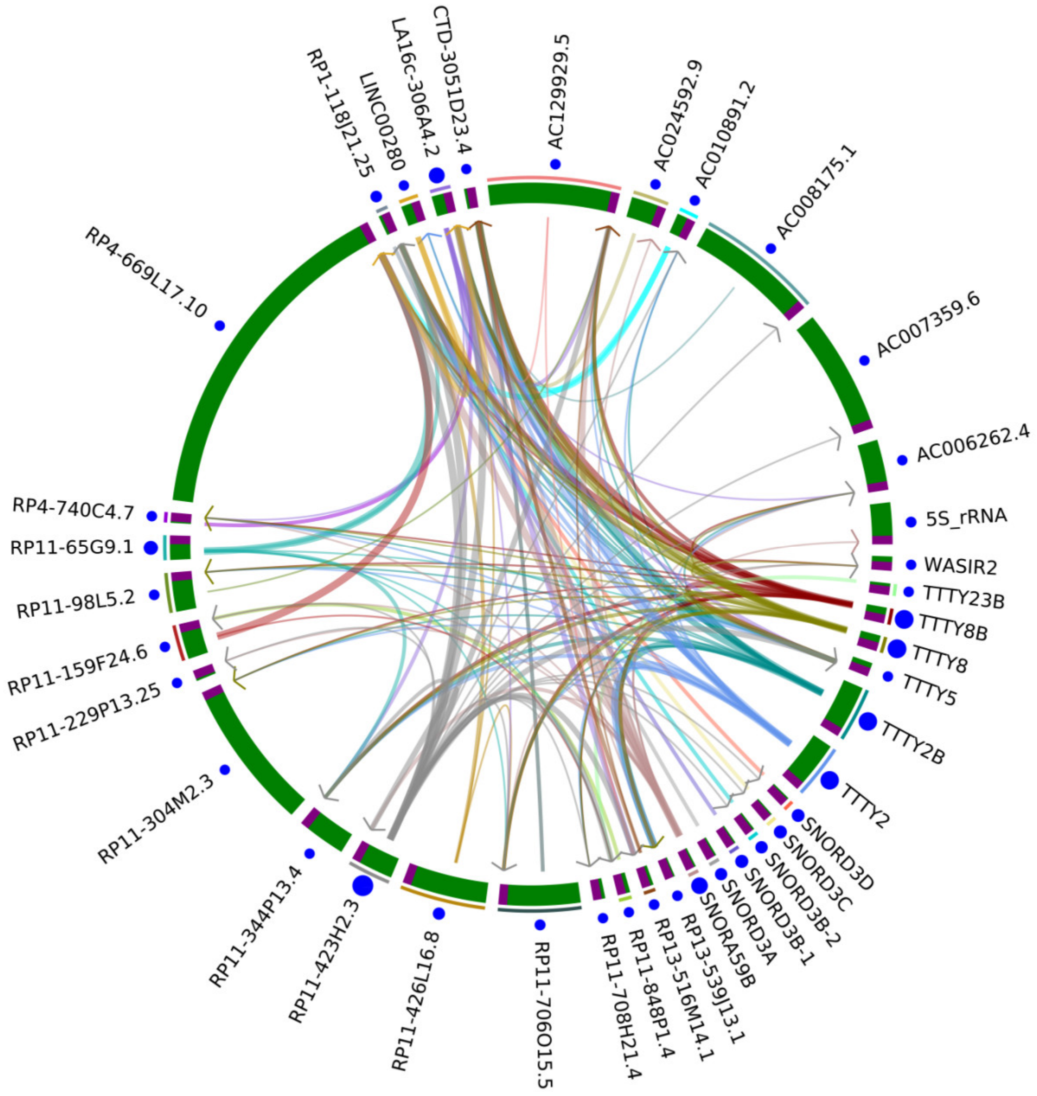


Supplementary Figure 6. Potential targeting regulation between HS lncRNAs. The circle's Brown and green regions indicate promoter and gene body regions. Arrows indicate the direction from the gene body to the promoter regions. The width of the arrows indicates the binding affinity of DBSs, and the sizes of blue dots indicate the number of DBSs of the lncRNA in the genome.

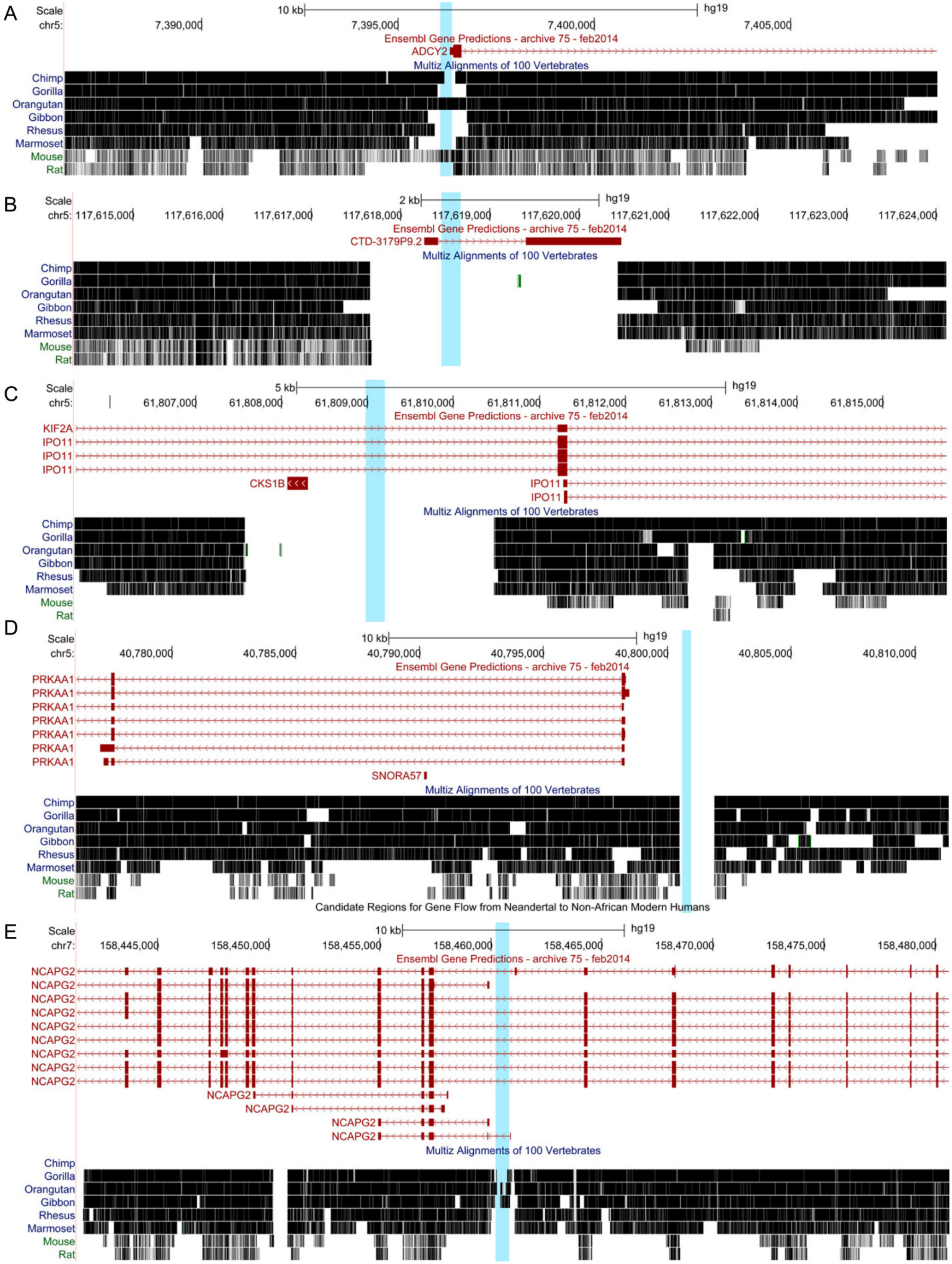


Supplementary Figure 7. Some DBSs (indicated by blue bars) are in human-specific genome sequences. (A-D) The DBSs of RP11-848P1.4 in the genes *ADCY2*, *CTD-3179P9.2*, *IPO11*, and *PRKAA1*. (E) The DBS of RP11-598D14.1 in the gene *NCAPG2*.

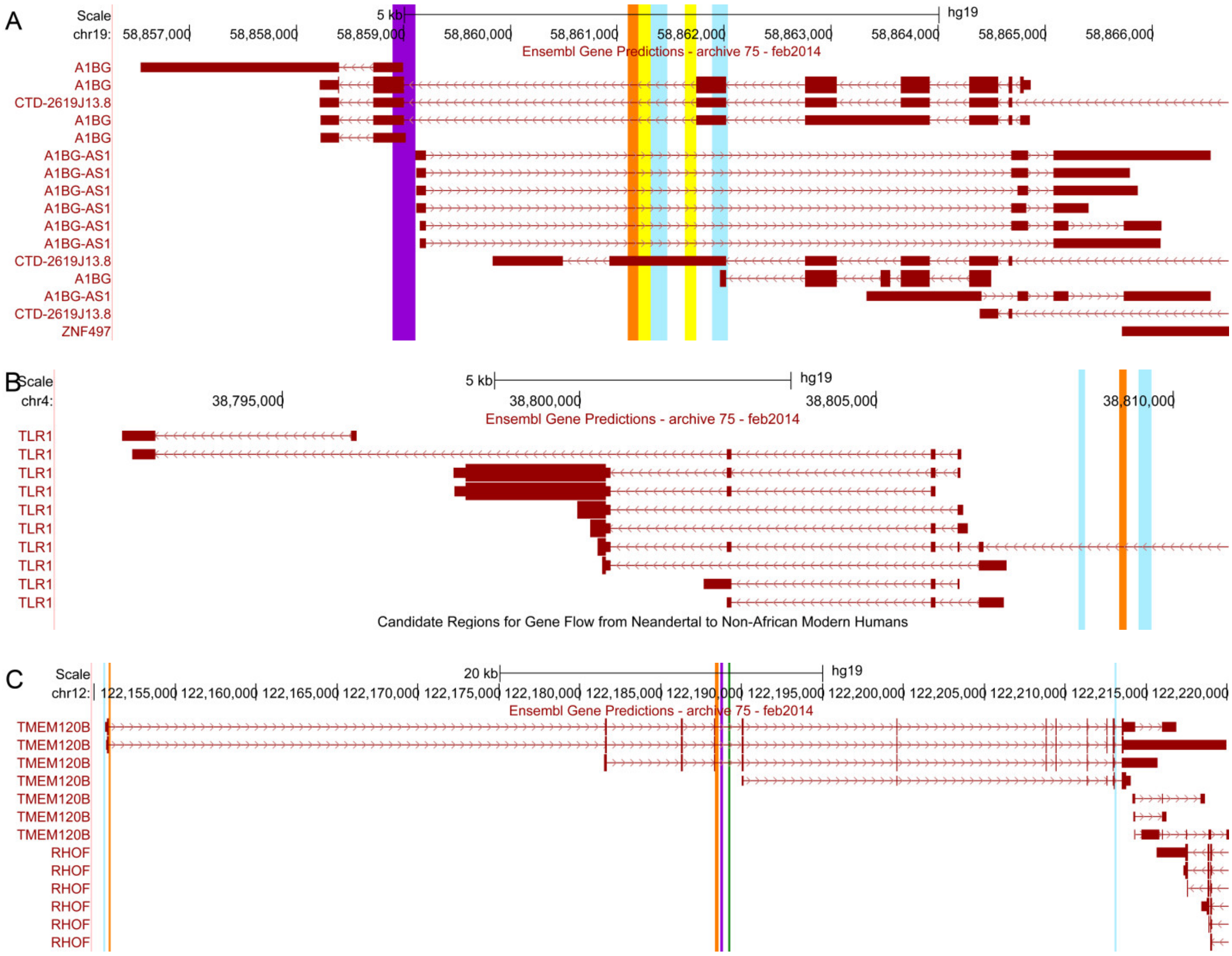


Supplementary Figure 8. Many genes and transcripts contain DBSs for multiple HS lncRNAs. (A) Left to right: the DBSs of RP11-65G9.1, LA16c-306A4.2, RP13-516M14.1, SNORA59B, RP11-423H2.3, and TTTY8/8B in the *A1BG*. (B) Left to right: the DBSs of TTTY8/8B, RP4-669L17.10, and RP11-423H2.3 in *TLR1*. (C) Left to right: the DBSs of LA16c-306A4.2, RP11-423H2.3, RP11-423H2.3, RP1-118J21.25, RP11-706O15.5, and SNORA59B in *TMEM210B*.

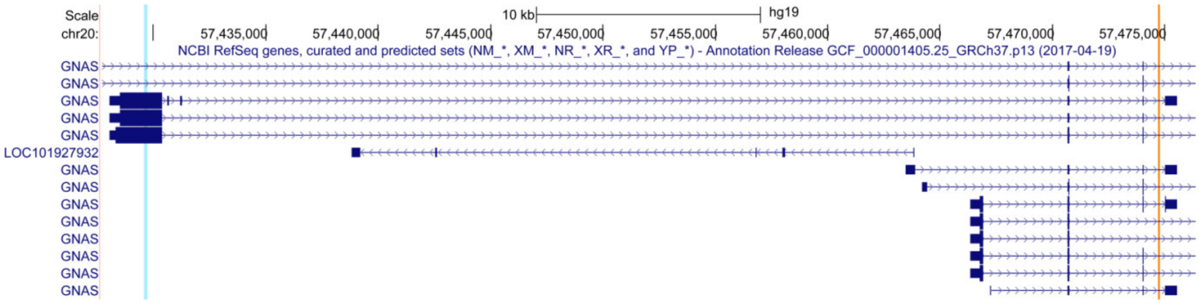


Supplementary Figure 9. In the *GNAS* region, RP11-423H2.3 has a DBS (indicated by the blue bar) wherein a selection signal was detected in CEU and CHB (Tajima's D=-0.99/-1.13/1.86 in CEU/CHB/YRI, integrated Fst=0.22), and has another DBS (indicated by the orange bar) wherein a selection signal was detected in YRI (Tajima's D=0.25/1.09/-1.17 in CEU/CHB/YRI, integrated Fst=0.33).

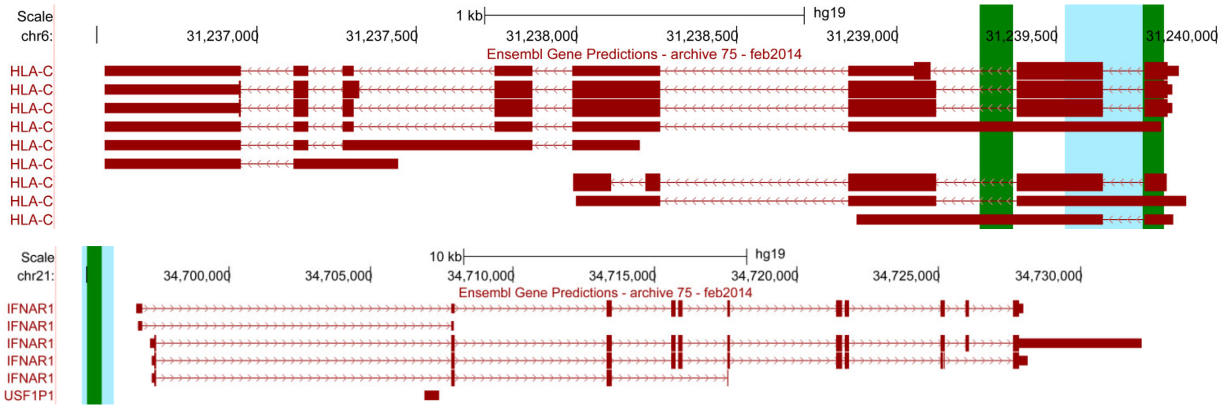


Supplementary Figure 10. HS lncRNAs on the Y chromosome often have longer DBSs than HS lncRNAs on the autosomes. The top panel shows that the DBS of TTTY2/2B in *HLA-C* (indicated by the blue bar) is longer than the two DBSs of RP11-423H2.3 (indicated by the green bars). The bottom panel shows that the DBS of TTTY8/8B in *IFNAR1* (indicated by the blue bar) is longer than the DBS of LINC00279 (indicated by the green bar).

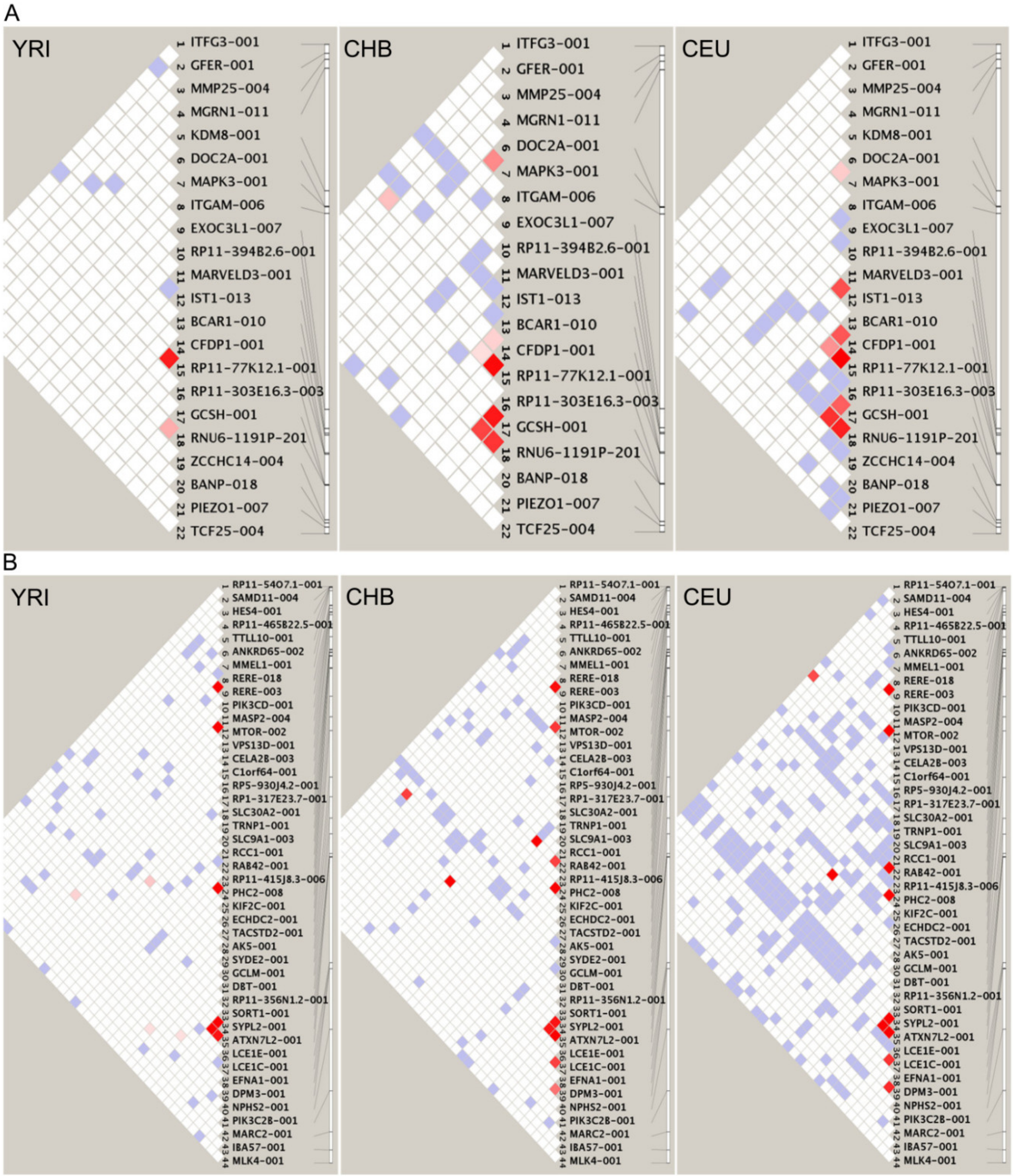
A
YRI
ITFG3-001
GFER-001
MMP25-004
MGRN1-011
KDM8-001
DOC2A-001
MAPK3-001
ITGAM-006
EXOC3L1-007
RP11-394B2.6-001
MARVELD3-001
IST1-013
BCAR1-010
CFDP1-001
RP11-77K12.1-001
RP11-303E16.3-003
GCSH-001
RNU6-1191P-201
ZCCHC14-004
BANP-018
PIEZO1-007
TCF25-004
CHB
ITFG3-001
GFER-001
MMP25-004
MGRN1-011
DOC2A-001
MAPK3-001
ITGAM-006
EXOC3L1-007
RP11-394B2.6-001
MARVELD3-001
IST1-013
BCAR1-010
CFDP1-001
RP11-77K12.1-001
RP11-303E16.3-003
GCSH-001
RNU6-1191P-201
BANP-018
PIEZO1-007
TCF25-004
CEU
ITFG3-001
GFER-001
MMP25-004
MGRN1-011
KDM8-001
DOC2A-001
MAPK3-001
ITGAM-006
EXOC3L1-007
RP11-394B2.6-001
MARVELD3-001
IST1-013
BCAR1-010
CFDP1-001
RP11-77K12.1-001
RP11-303E16.3-003
GCSH-001
RNU6-1191P-201
ZCCHC14-004
BANP-018
PIEZO1-007
TCF25-004
B
YRI
CHB
CEU
RP11-54O7.1-001
SAMD11-004
HES4-001
RP11-465B22.5-001
TTLL10-001
ANKRD65-002
MMEL1-001
RERE-018
RERE-003
PIK3CD-001
MASP2-004
MTOR-002
VPS13D-001
CELA2B-003
C1orf64-001
RP5-930J4.2-001
RP1-317E23.7-001
SLC30A2-001
TRNP1-001
SLC9A1-003
RCC1-001
RAB42-001
RP11-415J8.3-006
PHC2-008
KIF2C-001
ECHDC2-001
TACSTD2-001
AK5-001
SYDE2-001
GCLM-001
DBT-001
RP11-356N1.2-001
SORT1-001
SYPL2-001
ATXN7L2-001
LCE1E-001
LCE1C-001
EFNA1-001
DPM3-001
NPHS2-001
PIK3C2B-001
MARC2-001
IBA57-001
MLK4-001

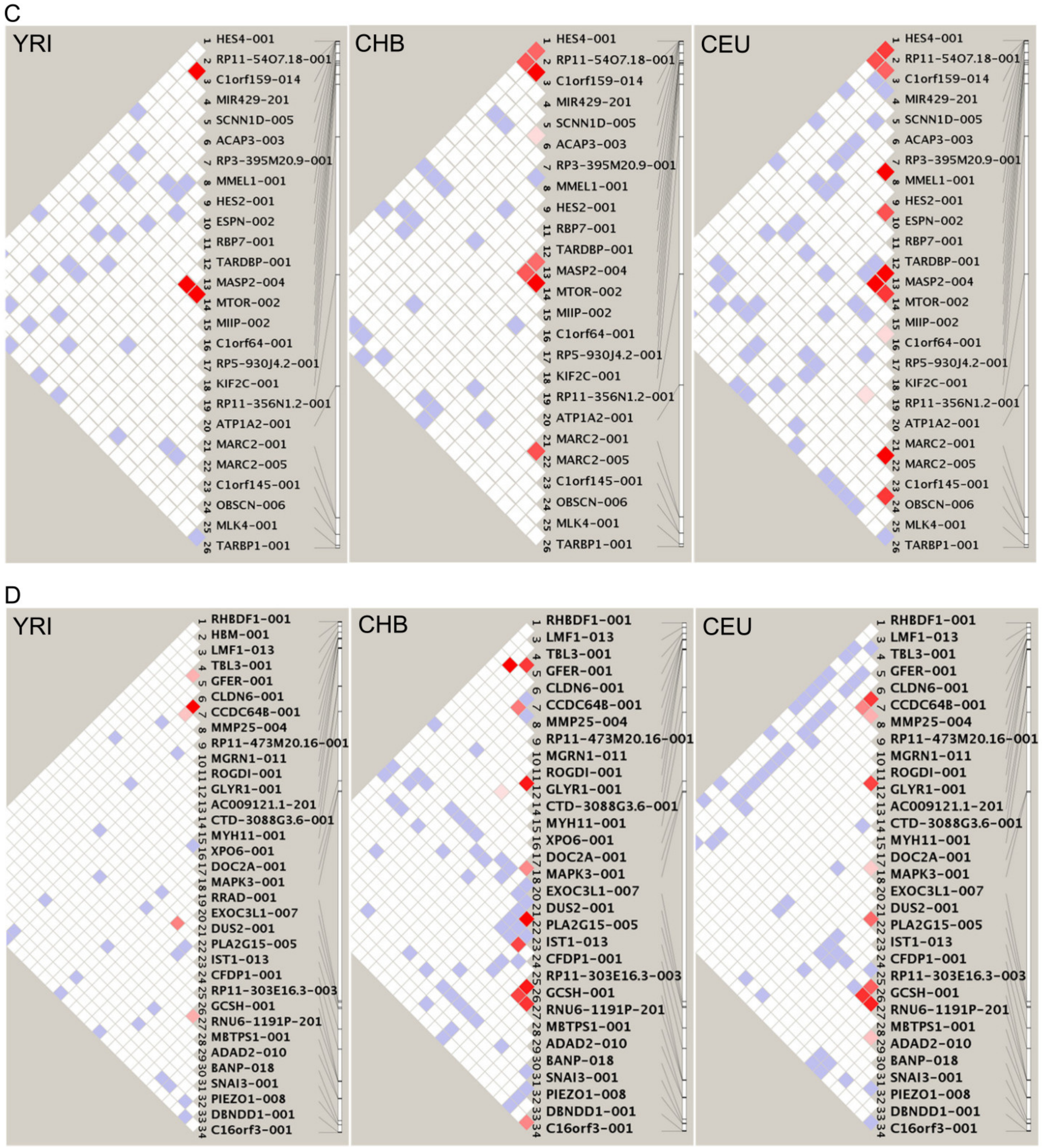


Supplementary Figure 11. The LD of the key SNP in DBSs of HS lncRNAs in genes on some chromosomes. (A) The LD of the key SNP in the DBSs of LA16c-306A4.2 in some genes on chromosome 16. (B) The LD of the key SNP in the DBSs of RP11-423H2.3 in some genes on chromosome 1. (C) The LD of the key SNP in the DBSs of SNORA59B in some genes on chromosome 1. (D) The LD of the key SNP in the DBSs of TTTY8B in some genes on chromosome 16.

**Supplementary Note 3 – Target genes with specific DBS features are enriched in specific functions**
Strong and weak DBSs may indicate well-established and recently occurred epigenetic regulation by HS lncRNAs. To examine if genes with DBSs of different features are enriched in different biological functions, we performed multiple over-representation analyses (ORA) using the *g:Profiler* program (Organism=Homo sapiens, Ordered query=No, Significance threshold=Benjamini-Hochberg FDR, User threshold=0.05, 50<terms size<1000) and the Gene Ontology (GO) database.

First, we examined the top 2000 genes and bottom 2000 genes, with the strongest and weakest DBSs, respectively (Supplementary Table 2). The two gene sets share many GO terms with different significance, including "neuron projection development" (GO:0031175) (2.36E-16, 3.79E-11), "neuron projection morphogenesis" (GO:0048812) (8.56E-16, 1.64E-09), and "female sex differentiation" (GO:0046660) (2.40E-2, 5.84E-04), yet genes in each set are also enriched in specific GO terms (Supplementary Table 3; Supplementary Figure 12). Of note, genes with weak DBSs are enriched in slightly more GO terms, including "response to alcohol" (GO:0097305), "negative regulation of locomotion" (GO:0040013), "female gonad development" (GO:0008585), and "response to temperature stimulus" (GO:0009266).

Next, we computed DBS sequence distances (per-base distances) in two ways. The first were the distances from the reconstructed human ancestor to chimpanzees, archaic humans, and modern humans; the second were the distances from modern humans to chimpanzees and archaic humans. We found that, when the human-chimpanzee ancestral sequence has the ancestral sequence (which means the inference of ancestral allele is of high confidence), more DBSs have a distance >0.015 from the human ancestor to archaic humans than to modern humans (Supplementary Figure 13). According to the file "homo_sapiens_ancestor_GRCh37_e71.README" and the paper "1000 Genomes Project Consortium. A global reference for human genetic variation. Nature 2015", a high-confidence call in the human ancestor sequence is made when all three sequences - the ancestral (the common ancestor of humans and chimpanzees), the sister (chimpanzees), and the ancestral of the ancestral sequences - agree. We found that only about 64% of calls in the human ancestor sequence are high-confidence calls, and that only DBSs in human ancestral loci with high-confidence have correct distances to the five leaf nodes. We therefore computed DBS distances from modern humans to archaic humans and chimpanzees.

To examine whether DBS distances computed using the second method reflect genetic changes in the human lineage or the chimpanzee lineage, we also computed DBS sequence distances with the most changed sequences between humans and gorillas. We found that DBS distances between humans and chimpanzees were significantly correlated with those between humans and gorillas (Spearman's rho=0.57, p=0.0) and that these DBSs had larger distances between humans and gorillas (Supplementary Figure 14A). These results suggest that the sequence differences of the most changed DBSs between humans and chimpanzees are determined mainly by genetic changes occurring in the human lineage, but not in the chimpanzee lineage. The same results were observed when the archaic humans were examined (Supplementary Figure 14B).

When distances were computed using the second method, for DBSs without chimpanzee counterparts, we assumed their human-chimpanzee distances were 10.0. Then, ORA was performed in two ways, based on genes sorted by human-chimpanzee DBS distance and human-Altai Neanderthal DBS distance, respectively(Supplementary Table 5). First, we used the top 25% and bottom 25% of genes to examine whether genes with large DBS distances are enriched for more human evolution-related GO terms than genes with small DBS distances. Second, we used the top 50% and bottom 50% of genes, intersected with ASE genes reported by Agoglia et al. 2021 (Agoglia et al., 2021). For genes intersected with significant ASE genes (p-adj<0.01), those with large DBS distances are enriched with more GO terms and also more human evolution-related GO terms than genes with small DBS distances (Supplementary Figure 15).

Finally, we classified strong and weak DBSs with "mostly changed" sequence distances into the following six classes. Since only about 20% of genes have human-chimpanzee DBS distances >=0.034, and much fewer genes have human-Altai Neanderthal DBS distances exceeding this value, 0.034 is a stringent (reliable) threshold for defining "mostly changed". DBSs with "Human-Chimp distance>0.034 AND Human-Altai Neanderthals distance=0" (little distance change occurred since Altai Neanderthals) were defined as old ones, and DBSs with "Human-Altai Neanderthals distance>0.034 OR Human-Denisovan distance>0.034" ("mostly changed" since Altai Neanderthals or Denisovans) were defined as young ones. These six classes - strong old, strong young, strong others, weak old, weak young, and weak others – therefore reflect early and late, and strong and weak, DBS sequence changes in human evolution. Based on favored and hitchhiking mutations in 17 human populations (Tang et al., 2022; Tang et al., 2023), we examined the number of favored and hitchhiking mutations in each class. Favored and hitchhiking mutations are most enriched in the weak young class.

| **Hitchhiking SNPs** | strong old | strong young | strong others | weak old | weak young | weak others |
|---|---|---|---|---|---|---|
| | 3/15685 | 11/163007 | 78/170389 | 10/180505 | 44/47251 | 57/168692 |
| **Favored SNPs** | strong old | strong young | strong others | weak old | weak young | weak others |
| | 0/10216 | 1/16040 | 4/92153 | 0/26532 | 5/31242 | 2/108014 |

| Term_name (highest binding affinity) | Term_id | Adjusted_p_value | Term_name (lowest binding affinity) | Term_id | Adjusted_p_value |
|---|---|---|---|---|---|
| small GTPase mediated signal transduction | GO:0007264 | 8.55E-17 | neuron projection development | GO:0031175 | 3.79E-11 |
| neuron projection development | GO:0031175 | 2.36E-16 | cell morphogenesis involved in differentiation | GO:0000904 | 5.36E-11 |
| cell projection morphogenesis | GO:0048858 | 5.53E-16 | cellular component morphogenesis | GO:0032989 | 6.81E-11 |
| neuron projection morphogenesis | GO:0048812 | 8.56E-16 | regulation of plasma membrane bounded cell projection | GO:0120035 | 2.58E-10 |
| plasma membrane bounded cell projection | GO:0120039 | 8.56E-16 | plasma membrane bounded cell projection | GO:0120039 | 3.44E-10 |
| cell junction organization | GO:0034330 | 1.96E-15 | regulation of anatomical structure morphogenesis | GO:0022603 | 4.34E-10 |
| cell part morphogenesis | GO:0032990 | 5.24E-15 | cell projection morphogenesis | GO:0048858 | 4.88E-10 |
| synaptic signaling | GO:0099536 | 1.38E-14 | cell part morphogenesis | GO:0032990 | 6.25E-10 |
| cellular component morphogenesis | GO:0032989 | 3.76E-14 | regulation of cell projection organization | GO:0031344 | 1.13E-09 |
| trans-synaptic signaling | GO:0099537 | 1.88E-13 | neuron projection morphogenesis | GO:0048812 | 1.64E-09 |
| cell morphogenesis involved in differentiation | GO:0000904 | 3.16E-13 | actin filament-based process | GO:0030029 | 1.65E-09 |
| regulation of small GTPase mediated signal transduction | GO:0051056 | 4.03E-13 | actin cytoskeleton organization | GO:0030036 | 1.99E-09 |
| chemical synaptic transmission | GO:0007268 | 4.18E-13 | organophosphate metabolic process | GO:0019637 | 2.05E-09 |
| anterograde trans-synaptic signaling | GO:0098916 | 4.18E-13 | cell morphogenesis involved in neuron differentiation | GO:0048667 | 2.47E-09 |
| regulation of plasma membrane bounded cell projection | GO:0120035 | 1.72E-12 | regulation of cellular component biogenesis | GO:0044087 | 8.54E-09 |
| regulation of cell projection organization | GO:0031344 | 1.74E-12 | regulation of neuron projection development | GO:0010975 | 1.22E-08 |
| cell morphogenesis involved in neuron differentiation | GO:0048667 | 3.83E-12 | cell junction organization | GO:0034330 | 3.32E-08 |
| dendrite development | GO:0016358 | 4.71E-11 | organophosphate biosynthetic process | GO:0090407 | 3.82E-08 |
| enzyme-linked receptor protein signaling pathway | GO:0007167 | 4.71E-11 | growth | GO:0040007 | 3.85E-08 |
| cell surface receptor signaling pathway involved in cell- | GO:1905114 | 4.86E-11 | developmental growth | GO:0048589 | 5.54E-08 |
| actin filament-based process | GO:0030029 | 7.42E-11 | positive regulation of protein modification process | GO:0031401 | 9.39E-08 |
| regulation of transmembrane transport | GO:0034762 | 8.88E-11 | regulation of cell morphogenesis | GO:0022604 | 1.08E-07 |
| synapse organization | GO:0050808 | 1.07E-10 | negative regulation of cellular component organization | GO:0051129 | 1.64E-07 |
| regulation of cellular component biogenesis | GO:0044087 | 1.17E-10 | lipid biosynthetic process | GO:0008610 | 2.44E-07 |
| metal ion transport | GO:0030001 | 4.76E-10 | positive regulation of transport | GO:0051050 | 3.52E-07 |
| Ras protein signal transduction | GO:0007265 | 5.80E-10 | regulation of locomotion | GO:0040012 | 4.21E-07 |
| regulation of ion transport | GO:0043269 | 7.96E-10 | organelle assembly | GO:0070925 | 4.42E-07 |
| modulation of chemical synaptic transmission | GO:0050804 | 8.08E-10 | regulation of cell migration | GO:0030334 | 4.90E-07 |
| regulation of trans-synaptic signaling | GO:0099177 | 8.80E-10 | mitotic cell cycle | GO:0000278 | 6.42E-07 |
| cation transmembrane transport | GO:0098655 | 1.65E-09 | synapse organization | GO:0050808 | 6.74E-07 |
| regulation of anatomical structure morphogenesis | GO:0022603 | 1.75E-09 | actomyosin structure organization | GO:0031032 | 1.15E-06 |
| dendrite morphogenesis | GO:0048813 | 2.21E-09 | regulation of cell motility | GO:2000145 | 1.19E-06 |
| behavior | GO:0007610 | 2.36E-09 | neuron migration | GO:0001764 | 1.61E-06 |
| transmembrane receptor protein tyrosine kinase | GO:0007169 | 2.84E-09 | positive regulation of cell projection organization | GO:0031346 | 1.87E-06 |
| circulatory system process | GO:0003013 | 5.41E-09 | smooth muscle cell migration | GO:0014909 | 2.31E-06 |
| regulation of GTPase activity | GO:0043087 | 6.17E-09 | axon guidance | GO:0007411 | 2.43E-06 |
| Wnt signaling pathway | GO:0016055 | 7.92E-09 | supramolecular fiber organization | GO:0097435 | 2.59E-06 |
| regulation of neuron projection development | GO:0010975 | 8.14E-09 | neuron projection guidance | GO:0097485 | 2.73E-06 |
| cell junction assembly | GO:0034329 | 8.85E-09 | positive regulation of phosphorus metabolic process | GO:0010562 | 2.83E-06 |
| cell-cell signaling by wnt | GO:0198738 | 9.63E-09 | positive regulation of phosphate metabolic process | GO:0045937 | 2.83E-06 |
| inorganic ion transmembrane transport | GO:0098660 | 1.02E-08 | mitotic cell cycle process | GO:1903047 | 3.17E-06 |
| regulation of vesicle-mediated transport | GO:0060627 | 1.26E-08 | muscle cell migration | GO:0014812 | 3.36E-06 |
| actin cytoskeleton organization | GO:0030036 | 1.35E-08 | cell-substrate adhesion | GO:0031589 | 3.41E-06 |
| axon development | GO:0061564 | 1.43E-08 | cell projection assembly | GO:0030031 | 3.49E-06 |
| supramolecular fiber organization | GO:0097435 | 1.70E-08 | non-membrane-bounded organelle assembly | GO:0140694 | 3.80E-06 |
| inorganic cation transmembrane transport | GO:0098662 | 2.55E-08 | transmembrane receptor protein tyrosine kinase | GO:0007169 | 4.50E-06 |
| regulation of ion transmembrane transport | GO:0034765 | 4.34E-08 | positive regulation of locomotion | GO:0040017 | 4.88E-06 |
| blood circulation | GO:0008015 | 8.44E-08 | positive regulation of phosphorylation | GO:0042327 | 5.13E-06 |
| protein localization to cell periphery | GO:1990778 | 8.54E-08 | head development | GO:0060322 | 5.78E-06 |
| axonogenesis | GO:0007409 | 8.84E-08 | plasma membrane bounded cell projection assembly | GO:0120031 | 8.27E-06 |
| actin filament organization | GO:0007015 | 1.08E-07 | nucleobase-containing small molecule metabolic process | GO:0055086 | 8.56E-06 |
| potassium ion transmembrane transport | GO:0071805 | 1.64E-07 | positive regulation of cell motility | GO:2000147 | 9.88E-06 |
| peptidyl-serine modification | GO:0018209 | 2.10E-07 | regulation of axonogenesis | GO:0050770 | 1.06E-05 |
| positive regulation of cell differentiation | GO:0045597 | 2.30E-07 | developmental cell growth | GO:0048588 | 1.07E-05 |
| regulation of system process | GO:0044057 | 2.59E-07 | positive regulation of cell migration | GO:0030335 | 1.08E-05 |
| vesicle-mediated transport in synapse | GO:0099003 | 2.97E-07 | organelle localization | GO:0051640 | 1.26E-05 |
| sodium ion transport | GO:0006814 | 2.97E-07 | microtubule-based process | GO:0007017 | 1.30E-05 |
| regulation of membrane potential | GO:0042391 | 3.55E-07 | positive regulation of cellular component movement | GO:0051272 | 1.42E-05 |
| positive regulation of GTPase activity | GO:0043547 | 5.52E-07 | brain development | GO:0007420 | 1.42E-05 |
| heart development | GO:0007507 | 5.60E-07 | microtubule cytoskeleton organization | GO:0000226 | 1.42E-05 |
| peptidyl-threonine modification | GO:0018210 | 5.72E-07 | axonogenesis | GO:0007409 | 1.84E-05 |
| regulation of locomotion | GO:0040012 | 8.80E-07 | ion homeostasis | GO:0050801 | 2.14E-05 |
| tissue morphogenesis | GO:0048729 | 8.95E-07 | cell growth | GO:0016049 | 2.14E-05 |
| regulation of cell motility | GO:2000145 | 9.11E-07 | protein deacetylation | GO:0006476 | 2.43E-05 |
| peptidyl-serine phosphorylation | GO:0018105 | 1.18E-06 | axon development | GO:0061564 | 2.53E-05 |
| regulation of cell migration | GO:0030334 | 1.21E-06 | chemotaxis | GO:0006935 | 2.56E-05 |
| sodium ion transmembrane transport | GO:0035725 | 1.31E-06 | regulation of cell size | GO:0008361 | 2.67E-05 |
| protein localization to plasma membrane | GO:0072659 | 1.42E-06 | nucleotide metabolic process | GO:0009117 | 2.67E-05 |
| potassium ion transport | GO:0006813 | 1.86E-06 | cell surface receptor signaling pathway involved in cell- | GO:1905114 | 2.77E-05 |
| regulation of cation transmembrane transport | GO:1904062 | 2.14E-06 | taxis | GO:0042330 | 2.88E-05 |
| growth | GO:0040007 | 2.35E-06 | microtubule cytoskeleton organization involved in mitosis | GO:1902850 | 3.22E-05 |
| regulation of metal ion transport | GO:0010959 | 2.53E-06 | regulation of vesicle-mediated transport | GO:0060627 | 3.41E-05 |
| actin filament bundle assembly | GO:0051017 | 2.69E-06 | enzyme-linked receptor protein signaling pathway | GO:0007167 | 3.51E-05 |
| regulation of growth | GO:0040008 | 2.85E-06 | cell-cell signaling by wnt | GO:0198738 | 3.54E-05 |
| regulation of Wnt signaling pathway | GO:0030111 | 3.13E-06 | behavior | GO:0007610 | 3.87E-05 |
| regulation of actin filament organization | GO:0110053 | 3.50E-06 | actin filament organization | GO:0007015 | 3.87E-05 |
| regulation of cell adhesion | GO:0030155 | 3.73E-06 | sensory organ development | GO:0007423 | 3.90E-05 |
| regulation of cellular localization | GO:0060341 | 3.98E-06 | neuron projection extension | GO:1990138 | 4.04E-05 |

Supplementary Figure 12. The enriched GO terms for the top 2000 and bottom 2000 genes with largest and smallest binding affinity. (A) The top 2000 genes with strong binding affinities (left) and the bottom 2000 genes with weak binding affinities (right) are enriched in many common GO terms (marked in pink).

| | | |
|---|---|---|
| positive regulation of epithelial cell migration | GO:0010634 | 4.03E-02 |
| establishment of vesicle localization | GO:0051650 | 4.07E-02 |
| protein localization to extracellular region | GO:0071692 | 4.19E-02 |
| positive regulation of small molecule metabolic process | GO:0062013 | 4.19E-02 |
| negative regulation of growth | GO:0045926 | 4.22E-02 |
| vascular associated smooth muscle cell proliferation | GO:1990874 | 4.24E-02 |
| nuclear envelope organization | GO:0006998 | 4.24E-02 |
| negative regulation of cell development | GO:0010721 | 4.35E-02 |
| excitatory postsynaptic potential | GO:0060079 | 4.35E-02 |
| phospholipid biosynthetic process | GO:0008654 | 4.38E-02 |
| negative regulation of smooth muscle cell proliferation | GO:0048662 | 4.41E-02 |
| regulation of cell-cell adhesion | GO:0022407 | 4.43E-02 |
| regulation of autophagy | GO:0010506 | 4.46E-02 |
| erythrocyte differentiation | GO:0030218 | 4.46E-02 |
| regulation of angiogenesis | GO:0045765 | 4.48E-02 |
| regulation of JNK cascade | GO:0046328 | 4.48E-02 |
| myelination | GO:0042552 | 4.48E-02 |
| cell chemotaxis | GO:0060326 | 4.48E-02 |
| receptor internalization | GO:0031623 | 4.56E-02 |
| apoptotic signaling pathway | GO:0097190 | 4.56E-02 |
| regulation of tissue remodeling | GO:0034103 | 4.64E-02 |
| regulation of cell size | GO:0008361 | 4.66E-02 |
| leukocyte differentiation | GO:0002521 | 4.68E-02 |
| regulation of neuron death | GO:1901214 | 4.70E-02 |
| focal adhesion assembly | GO:0048041 | 4.72E-02 |
| response to metal ion | GO:0010038 | 4.75E-02 |
| smooth muscle cell differentiation | GO:0051145 | 4.75E-02 |
| phosphatidylinositol-mediated signaling | GO:0048015 | 4.84E-02 |
| cellular carbohydrate metabolic process | GO:0044262 | 4.84E-02 |
| regulation of transmembrane receptor protein serine/threonine kinase signaling pathway | GO:0090092 | 4.84E-02 |
| positive regulation of cellular carbohydrate metabolic | GO:0010676 | 4.84E-02 |
| positive regulation of stress fiber assembly | GO:0051496 | 4.84E-02 |
| detection of mechanical stimulus | GO:0050982 | 4.84E-02 |
| cell maturation | GO:0048469 | 4.84E-02 |
| negative regulation of fat cell differentiation | GO:0045599 | 4.84E-02 |
| synaptic vesicle localization | GO:0097479 | 4.84E-02 |
| cellular metal ion homeostasis | GO:0006875 | 4.84E-02 |
| glucose metabolic process | GO:0006006 | 4.85E-02 |
| regulation of cell-substrate junction organization | GO:0150116 | 4.85E-02 |
| cellular response to chemical stress | GO:0062197 | 4.91E-02 |
| positive regulation of cell population proliferation | GO:0008284 | 4.91E-02 |
| negative regulation of cellular catabolic process | GO:0031330 | 4.91E-02 |
| smooth muscle contraction | GO:0006939 | 4.91E-02 |
| regulation of ERBB signaling pathway | GO:1901184 | 4.98E-02 |
| cardiac muscle hypertrophy | GO:0003300 | 4.98E-02 |

| | | |
|---|---|---|
| vesicle cytoskeletal trafficking | GO:0099518 | 3.15E-02 |
| protein complex oligomerization | GO:0051259 | 3.20E-02 |
| glycoprotein biosynthetic process | GO:0009101 | 3.20E-02 |
| negative regulation of catalytic activity | GO:0043086 | 3.21E-02 |
| cell-substrate junction assembly | GO:0007044 | 3.23E-02 |
| regulation of metaphase/anaphase transition of cell cycle | GO:1902099 | 3.26E-02 |
| chloride transport | GO:0006821 | 3.31E-02 |
| DNA modification | GO:0006304 | 3.31E-02 |
| regulation of protein targeting | GO:1903533 | 3.31E-02 |
| lymphocyte activation involved in immune response | GO:0002285 | 3.31E-02 |
| inner ear development | GO:0048839 | 3.31E-02 |
| ruffle organization | GO:0031529 | 3.31E-02 |
| secretion by cell | GO:0032940 | 3.33E-02 |
| positive regulation of chemotaxis | GO:0050921 | 3.39E-02 |
| lipid catabolic process | GO:0016042 | 3.40E-02 |
| response to ketone | GO:1901654 | 3.42E-02 |
| mitotic spindle assembly | GO:0090307 | 3.47E-02 |
| regulation of synaptic transmission, glutamatergic | GO:0051966 | 3.47E-02 |
| response to alcohol | GO:0097305 | 3.47E-02 |
| muscle structure development | GO:0061061 | 3.47E-02 |
| negative regulation of programmed cell death | GO:0043069 | 3.47E-02 |
| positive regulation of proteasomal ubiquitin-dependent protein catabolic process | GO:0032436 | 3.50E-02 |
| regulation of protein kinase B signaling | GO:0051896 | 3.58E-02 |
| purine ribonucleotide biosynthetic process | GO:0009152 | 3.58E-02 |
| cellular response to environmental stimulus | GO:0104004 | 3.59E-02 |
| cellular response to abiotic stimulus | GO:0071214 | 3.59E-02 |
| muscle system process | GO:0003012 | 3.59E-02 |
| regulation of mitotic sister chromatid separation | GO:0010965 | 3.61E-02 |
| detection of abiotic stimulus | GO:0009582 | 3.63E-02 |
| regulation of secretion | GO:0051046 | 3.63E-02 |
| biological process involved in interaction with host | GO:0051701 | 3.63E-02 |
| membrane organization | GO:0061024 | 3.63E-02 |
| negative regulation of apoptotic process | GO:0043066 | 3.66E-02 |
| positive regulation of Wnt signaling pathway | GO:0030177 | 3.69E-02 |
| multi-organism reproductive process | GO:0044703 | 3.69E-02 |
| ncRNA transcription | GO:0098781 | 3.71E-02 |
| proteoglycan biosynthetic process | GO:0030166 | 3.71E-02 |
| gamete generation | GO:0007276 | 3.74E-02 |
| transport across blood-brain barrier | GO:0150104 | 3.74E-02 |
| vascular transport | GO:0010232 | 3.74E-02 |
| regulation of cell division | GO:0051302 | 3.76E-02 |
| TOR signaling | GO:0031929 | 3.76E-02 |
| cellular lipid catabolic process | GO:0044242 | 3.77E-02 |
| positive regulation of lipid metabolic process | GO:0045834 | 3.77E-02 |
| regulation of mitochondrion organization | GO:0010821 | 3.77E-02 |
| cellular response to inorganic substance | GO:0071241 | 3.77E-02 |
| actin filament polymerization | GO:0030041 | 3.79E-02 |
| negative regulation of synaptic transmission | GO:0050805 | 3.80E-02 |
| glycolipid metabolic process | GO:0006664 | 3.80E-02 |
| carboxylic acid catabolic process | GO:0046395 | 3.80E-02 |
| regulation of epithelial cell proliferation | GO:0050678 | 3.83E-02 |
| response to radiation | GO:0009314 | 3.85E-02 |
| protein methylation | GO:0006479 | 3.86E-02 |
| protein alkylation | GO:0008213 | 3.86E-02 |
| Golgi organization | GO:0007030 | 3.88E-02 |
| membrane depolarization | GO:0051899 | 3.97E-02 |
| skeletal system morphogenesis | GO:0048705 | 3.98E-02 |
| positive chemotaxis | GO:0050918 | 3.98E-02 |
| development of primary sexual characteristics | GO:0045137 | 3.98E-02 |
| metaphase/anaphase transition of cell cycle | GO:0044784 | 3.98E-02 |
| non-motile cilium assembly | GO:1905515 | 3.98E-02 |
| bone development | GO:0060348 | 3.98E-02 |
| muscle contraction | GO:0006936 | 4.00E-02 |
| monovalent inorganic cation homeostasis | GO:0055067 | 4.00E-02 |
| chromatin organization | GO:0006325 | 4.01E-02 |
| positive regulation of response to external stimulus | GO:0032103 | 4.06E-02 |
| negative regulation of transmembrane transport | GO:0034763 | 4.06E-02 |
| anatomical structure homeostasis | GO:0060249 | 4.06E-02 |
| alpha-amino acid metabolic process | GO:1901605 | 4.07E-02 |
| cytokinesis | GO:0000910 | 4.09E-02 |
| regulation of regulated secretory pathway | GO:1903305 | 4.09E-02 |
| response to calcium ion | GO:0051592 | 4.09E-02 |
| regulation of chromosome segregation | GO:0051983 | 4.09E-02 |
| protein localization to cytoskeleton | GO:0044380 | 4.09E-02 |
| regulation of neurotransmitter transport | GO:0051588 | 4.09E-02 |
| liposaccharide metabolic process | GO:1903509 | 4.09E-02 |
| nucleotide catabolic process | GO:0009166 | 4.17E-02 |
| fatty acid biosynthetic process | GO:0006633 | 4.26E-02 |

Supplementary Figure 12. (B) The bottom 2000 genes with weak binding affinities (right) are also enriched in many specific GO terms (marked in yellow) with relatively low significance.

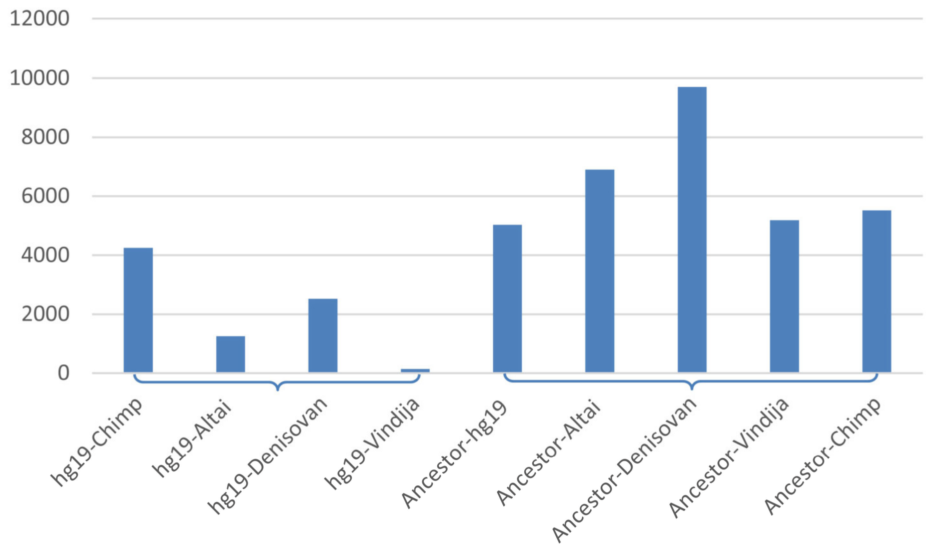


Supplementary Figure 13. Numbers of DBSs with large distances from modern humans to archaic humans and chimpanzees, and from the human ancestor to chimpanzees, archaic humans, and modern humans. Left: DBSs in 4248, 1256, 2513, 134 genes have distances >0.034 from modern humans to chimpanzees, Altai Neanderthals, Denisovans, and Vindija Neanderthals. Right: DBSs in 5033, 6908, 9707, 5189, and 5521 genes have distances >0.015 from the ancestor to modern humans, Altai Neanderthals, Denisovans, Vindija Neanderthals, and chimpanzees.

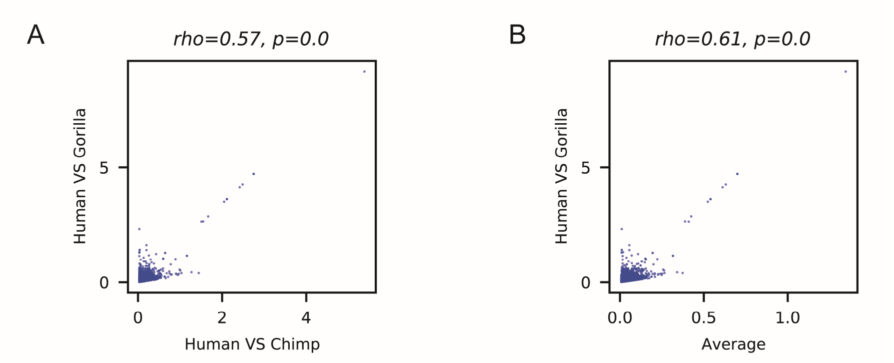


Supplementary Figure 14. The most changed DBSs also have large sequence distances between humans and gorillas. (A) Scatter plot showing the sequence distances between humans and chimpanzees and between humans and gorillas. (B) The scatter plot shows the average sequence distances between humans and chimpanzees, the three archaic humans, and between humans and gorillas. The rho and p values were estimated using the Spearman correlation test.

| Intersection of top 50% of genes sorted by DBS distance from chimpanzees to humans and ASE genes (adj-p<0.01) | term_id | adj_p | Intersection of bottom 50% of genes sorted by DBS distance from chimpanzees to humans and ASE genes (adj-p<0.01) | term_id | adj_p |
|---|---|---|---|---|---|
| cellular pigmentation | GO:0033059 | 2.41E-06 | brain development | GO:0007420 | 7.80E-03 |
| behavior | GO:0007610 | 5.78E-05 | forebrain development | GO:0030900 | 4.34E-02 |
| pigmentation | GO:0043473 | 7.68E-05 | | | |
| learning | GO:0007612 | 3.60E-04 | | | |
| associative learning | GO:0008306 | 2.08E-03 | | | |
| adaptive thermogenesis | GO:1990845 | 2.28E-03 | | | |
| sensory system development | GO:0048880 | 3.19E-03 | | | |
| cold-induced thermogenesis | GO:0106106 | 3.78E-03 | | | |
| digestive system development | GO:0055123 | 3.94E-03 | | | |
| glucose metabolic process | GO:0006006 | 3.97E-03 | | | |
| learning or memory | GO:0007611 | 4.69E-03 | | | |
| cognition | GO:0050890 | 6.32E-03 | | | |
| regulation of cold-induced thermogenesis | GO:0120161 | 6.33E-03 | | | |
| memory | GO:0007613 | 4.42E-02 | | | |
| alcohol metabolic process | GO:0006066 | 4.82E-02 | | | |

| Intersection of top 50% of genes sorted by DBS distance from Altai Neanderthals to humans and ASE genes (adj-p<0.01) | term_id | adj_p | Intersection of bottom 50% of genes sorted by DBS distance from Altai Neanderthals to humans and ASE genes (adj-p<0.01) | term_id | adj_p |
|---|---|---|---|---|---|
| behavior | GO:0007610 | 2.52E-07 | pigmentation | GO:0043473 | 4.39E-04 |
| glucose metabolic process | GO:0006006 | 9.29E-04 | cellular pigmentation | GO:0033059 | 2.74E-03 |
| sensory system development | GO:0048880 | 1.05E-03 | brain development | GO:0007420 | 4.96E-02 |
| learning | GO:0007612 | 1.77E-03 | | | |
| learning or memory | GO:0007611 | 3.62E-03 | | | |
| cognition | GO:0050890 | 4.42E-03 | | | |
| associative learning | GO:0008306 | 6.95E-03 | | | |
| digestive system development | GO:0055123 | 8.54E-03 | | | |
| cold-induced thermogenesis | GO:0106106 | 1.43E-02 | | | |
| adaptive thermogenesis | GO:1990845 | 1.50E-02 | | | |
| brain development | GO:0007420 | 1.85E-02 | | | |
| forebrain development | GO:0030900 | 2.04E-02 | | | |
| regulation of cold-induced thermogenesis | GO:0120161 | 2.28E-02 | | | |
| alcohol metabolic process | GO:0006066 | 2.38E-02 | | | |
| memory | GO:0007613 | 2.72E-02 | | | |
| visual behavior | GO:0007632 | 4.57E-02 | | | |

Supplementary Figure 15. Enriched GO terms of different sets of genes with large and small DBS distances from humans to chimpanzees and Altai Neanderthals. Shown are the presence and absence of GO terms highly related to human evolution. The intersections of genes sorted by DBS distance from humans to chimpanzees and to Altai Neanderthals, respectively, and genes showing significant ASE (adj-p<0.01 and |LFC|>0.5).

### Supplementary Note 4 – Positive selection signals in HS lncRNA genes

We used multiple tests, including XP-CLR (Chen et al., 2010), iSAFE (Akbari et al., 2018), Tajima's D (Tajima, 1989), the fixation index (Fst) (Weir and Cockerham, 1984), and linkage disequilibrium (LD) (Slatkin, 2008), to detect positive selection signals in HS lncRNA genes.

First, we used the XP-CLR program to scan the genome regions that contain HS lncRNAs and their 500-kb upstream and downstream sequences. Six pairwise comparisons were applied to the three human populations (CEU-CHB, CEU-YRI, CHB-CEU, CHB-YRI, YRI-CEU, and YRI-CHB). Selective sweeps were detected in the genome regions containing RP11-848P1.4 and RP11-598D14.1 in the CEU-YRI and CHB-YRI comparisons (Supplementary Figure 16). Abundant SNPs in these regions have low derived allele frequency (DAF) in YRI, but are nearly fixed in CEU and CHB. Examples (with DAF in YRI, CEU, and CHB) include rs7208589 (DAF = 0.162, 0.990, and 0.995), rs8073226 (DAF = 0.148, 0.990, and 0.995), and rs9915124 (DAF = 0.181, 0.990, and 0.995) in the region containing RP11-848P1.4, and rs4690648 (DAF = 0.185, 0.939, and 0.917), rs11722101 (DAF = 0.269, 0.970, and 0.917), and rs11730933 (DAF = 0.292,

0.970, and 0.917) in the region containing RP11-598D14.1.

Second, we used the iSAFE program to detect favored mutations in HS lncRNA genes in these populations. Because selective sweeps caused by favored mutations may have varied lengths, we ran iSAFE four times with genome regions of 80, 500, 1000, and 2500 kb, which centered at the HS lncRNA. Selective sweeps and favored mutations were detected robustly in genome regions containing RP11-598D14.1, RP11-848P1.4, AC006129.1, AC006129.4, CTD-2291D10.1, CTD-2291D10.2, and LA16c-306A4.2 in CEU and CHB, and in the genome region containing CTD-3051D23.4 in CHB (Supplementary Figure 17). Mutations with top iSAFE scores were located in the gene body regions of HS lncRNAs, and their DAF values were low in YRI but high in CEU and CHB. In addition, all of the detected favored mutations have high LD ($r^2$) scores.

Third, we used Tajima's D and integrated Fst to detect positive selection signals in gene body regions of HS lncRNAs. Positive selection signals were detected in RP11-598D14.1, RP11-848P1.4, RP11-344P13.4, and RP11-426L16.8 in CEU and CHB, in AC129929.5 and RP11-423H2.3 in CEU, and in CTB-151G24.1 in CHB (Supplementary Figure 18). Since Tajima's D values were referenced with the genome-wide background, D<0 and D>0 indicate positive (or directional selection) and balancing selection, respectively, instead of population demography dynamics. The Fst of each HS lncRNA gene, also referenced with the genome-wide background, was computed for the CEU-YRI, CHB-YRI, and CHB-CEU comparisons. Extreme Fst values of SNPs were detected in RP11-598D14.1 and AC129929.5 in the comparisons of CEU-YRI and CHB-YRI. Since Fst values were referenced with the genome-wide background, extreme Fst values indicate positive selection.

Finally, we applied LD analysis to each HS lncRNA gene. Significantly increased LD was detected in SNPs in AC024592.9, AC129929.5, RP11-157B13.7, RP11-277P12.10, CTD-2142D14.1, and CTD-2291D10.1 in CEU and CHB (Supplementary Figure 19). Taken together, the above results indicate that HS lncRNA genes may have undergone more significant adaptive evolution in CEU and CHB than in YRI.

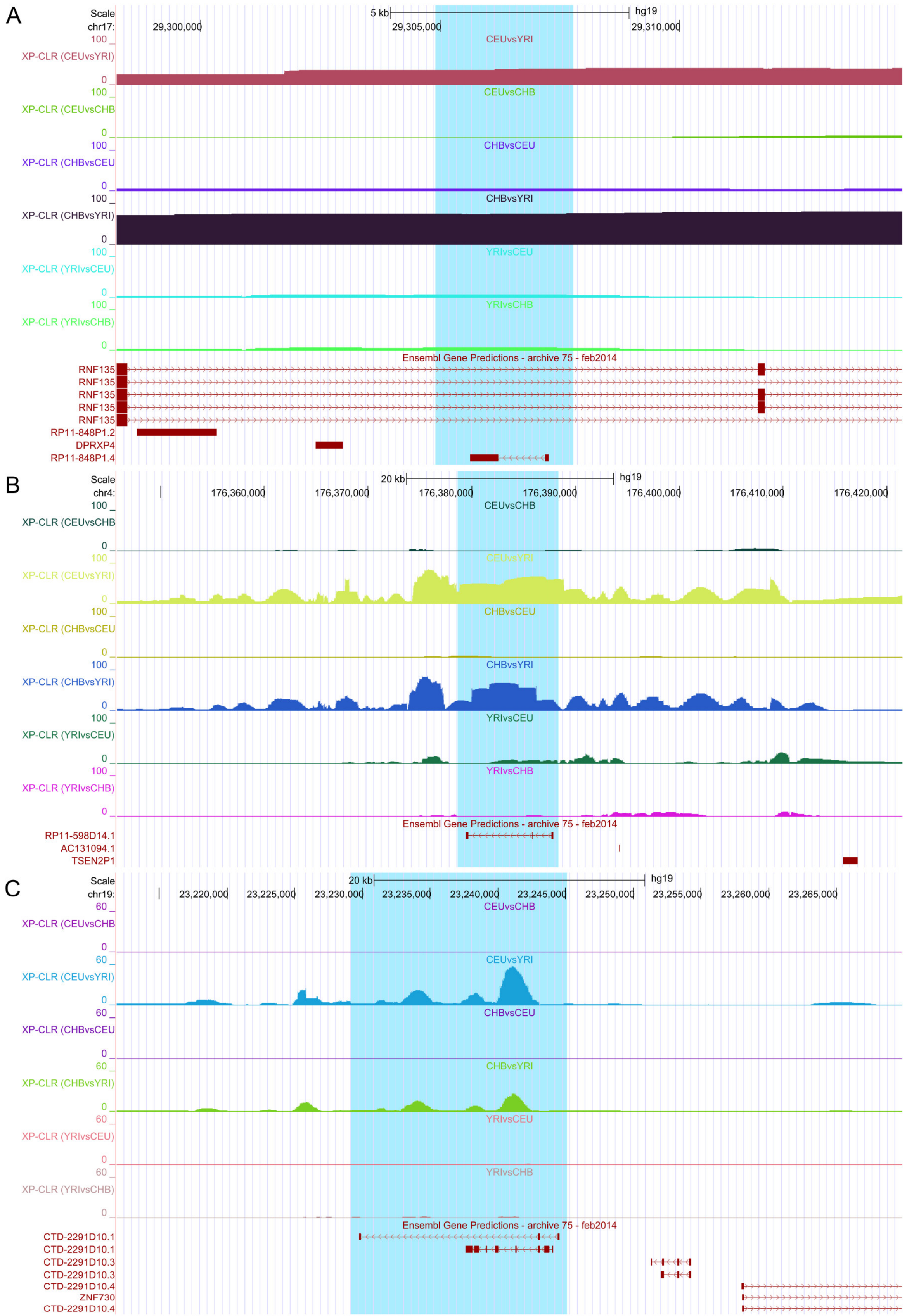


Supplementary Figure 16. Positive selection signals detected by the XP-CLR program in (A) RP11-848P1.4, (B) RP11-598D14.1, (C) CTD-2291D10.1 in CEU and CHB.

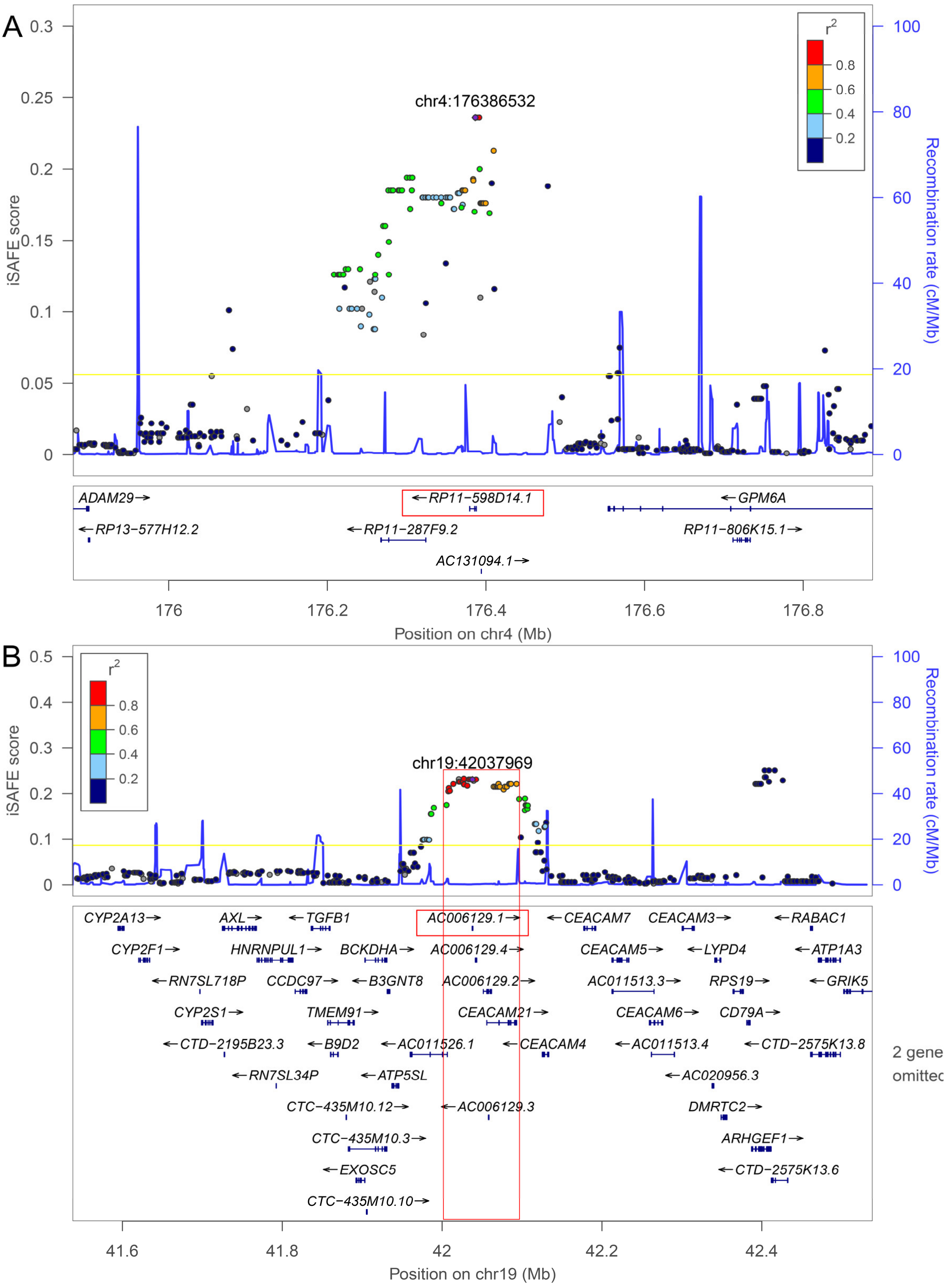


Supplementary Figure 17. Favored mutations detected by iSAFE. Left and right vertical axes indicate iSAFE scores and recombination rate. The purple diamond marks the top-scored mutation. Colors mark LD (r2) between the top-scoring mutation and others. The yellow line indicates that mutations above it have a probability of p=1e-6 to be neutral. The blue curve indicates the position-specific recombination rates. (A) SNPs in *RP11-598D14.1*. The top-scoring SNP has DAFs of 0.125/0.960/0.922 in YRI/CEU/CHB. (B) SNPs in *AC006129.1*. The top-scoring SNP has DAFs of 0.134/0.717/0.587 in YRI/CEU/CHB.

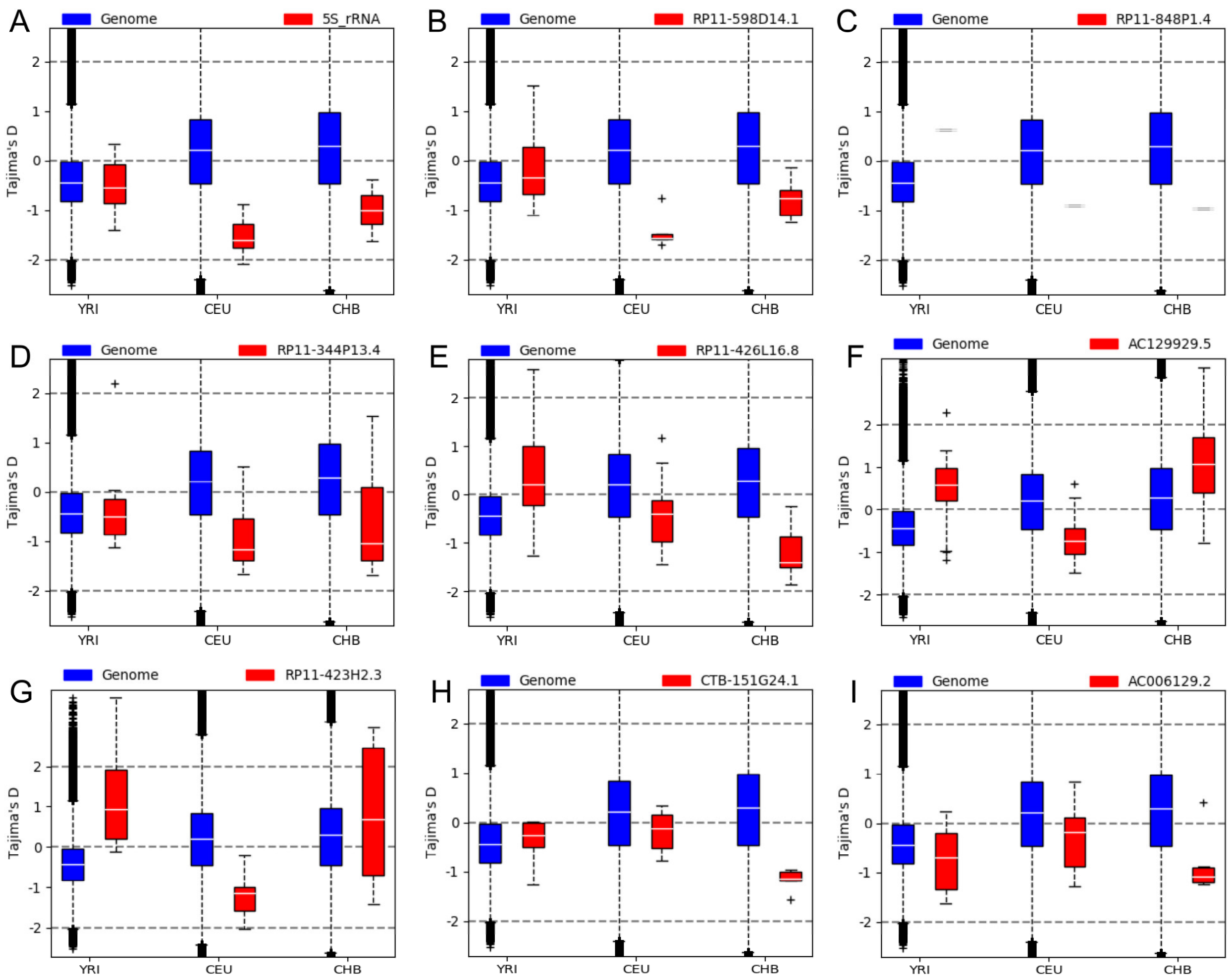


Supplementary Figure 18. HS lncRNA genes with significantly changed Tajima's D in CEU, CHB, and YRI. Negative and positive Tajima's D scores, which are significantly smaller or larger than the genome-wide background in a population, indicate the signature of positive selection or balancing selection, respectively, in the population.

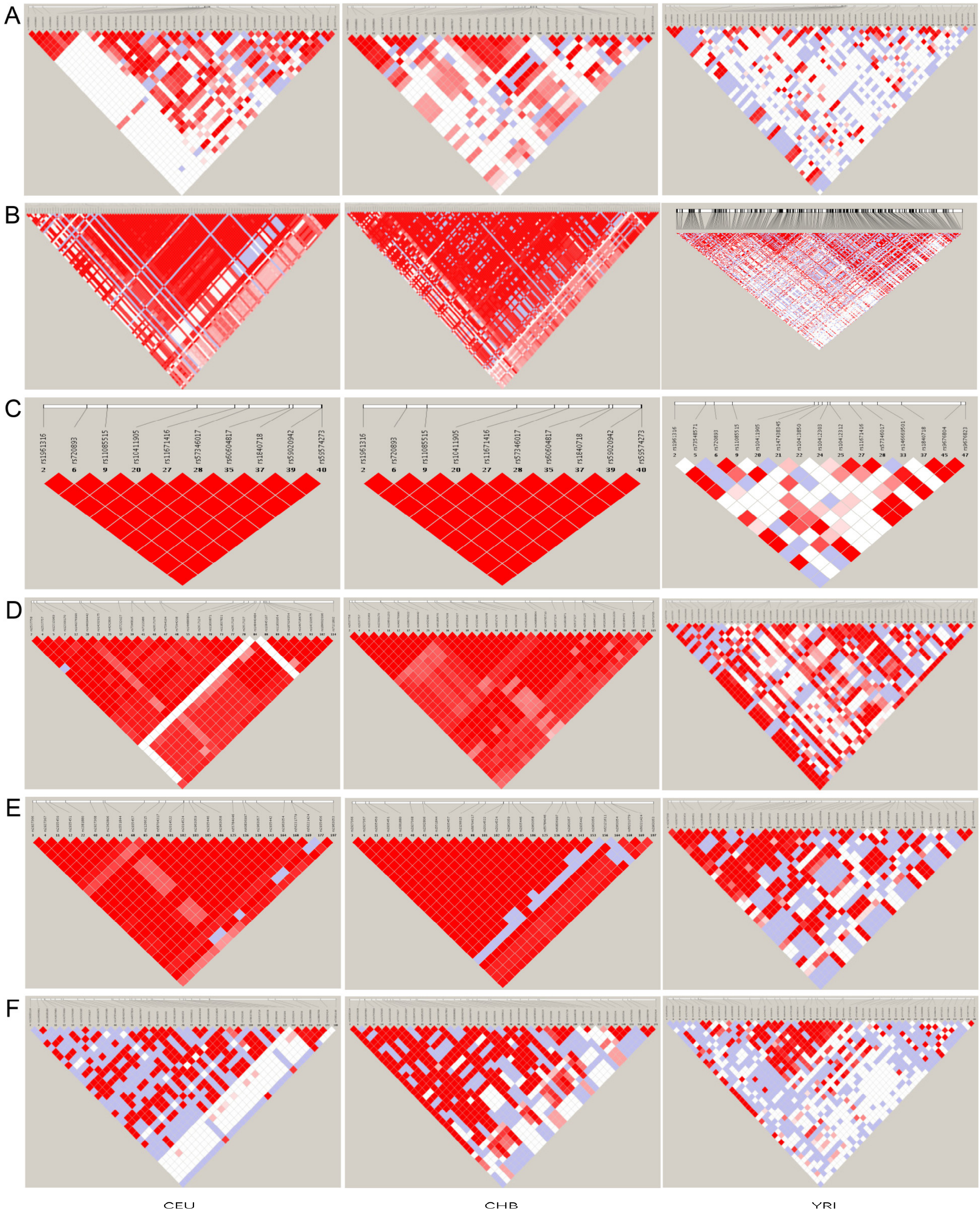


Supplementary Figure 19. The LD of SNPs in HS lncRNA genes in CEU, CHB, and YRI. The red color indicates high LD values. These panels show that LD between SNPs in CEU and CHB in these lncRNA genes is stronger than LD between SNPs in YRI. (A) AC024592.9. (B) AC129929.5. (C) RP11-157B13.7. (D) RP11-277P12.10. (E) CTD-2291D10.1. (F) CTD-2142D14.1.

**Supplementary Note 5 – Positive selection signals in DBSs**

We used the tests mentioned above, including Tajima's D (Tajima, 1989), Fay and Wu's H (Fay and Wu, 2000), integrated Fst (Weir and Cockerham, 1984), and LD (Slatkin, 2008), to detect positive selection signals in DBSs in each population. These tests reveal positive selection signals in DBSs in specific target genes in specific populations (Supplementary Table 9-11).

First, we calculated Tajima's D, Fst, and integrated Fst for polymorphic DBSs in CEU, CHB, and YRI. Positive selection signals were detected in DBSs in more genes in CEU and CHB than in YRI (Supplementary Table 10). These genes include the two pigmentation-related genes *MC1R* and *MFSD12*, the two odor reception-related genes *OR6C1* and *TAS1R3*, and the immune-related gene *TLR1.* These target genes may suggest adaptations in gene expression regulation in CEU and CHB in response to changes in diet and the environment. In YRI, positive selection signals were detected in DBSs in genes such as *SLCO4A1,* which encodes a protein mediating the Na-independent transport of organic anions (e.g., the thyroid hormones T3 and T4). DBSs in different transcripts of *GNAS* (a gene important for genomic imprinting) contain SNPs selected explicitly in different populations (Supplementary Figure 9).

Next, we calculated Fay-Wu's H and integrated Fst for each polymorphic DBS in CEU, CHB, and YRI. Strong negative values (H <- 2), together with significantly large integrated Fst (> 0.22), were obtained mainly in DBSs in CEU and CHB (Supplementary Table 11). These genes fall into two classes. The first class includes *PASK, CPT1A*, and *EXOC7,* which are important for glucose and lipid metabolism. *PASK* encodes a protein that plays a role in the regulation of insulin gene expression. *CPT1A* encodes a protein that exerts an important role in triglyceride metabolism. *EXOC7* encodes a protein that plays a crucial role in targeting SLC2A4 vesicles to the plasma membrane in response to insulin in adipocytes. The second class includes *COMT, TAS1R3*, and *ALMS1,* which are important for neural development. *COMT* is involved in the metabolism of adrenaline and noradrenaline. *TAS1R3* encodes a protein important for recognizing diverse natural and synthetic sweeteners. Mutations in *ALMS1* are associated with Alstrom syndrome.

Third, to examine whether gene expression regulation by HS lncRNAs is coordinated, we examined the LD between SNPs in DBSs of the same HS lncRNA chromosome-wide. When more than one SNP in a DBS has a minimal allele frequency (MAF) >= 0.05, we chose the SNP that had the strongest LD ($r^2$). Despite genetic recombination, LD between DBSs was detected on some chromosomes in CEU and CHB (Supplementary Figure 11; Supplementary Note 2).

Fourth, we computed the distances between DBSs in modern humans and their counterparts in archaic humans, and compared them with the distances between annotated promoters and their counterparts in archaic humans. A considerable portion of DBSs (note that DBSs are within promoter regions) have larger distances than promoters between modern and archaic humans (Supplementary Figure 20). We also computed the frequency distribution of SNPs (>0.05) in DBSs and found that SNPs are enriched at low and high frequencies (Supplementary Figure 21), indicating positive selection.

Finally, we analyzed two experimental datasets. It was reported that the expression of a set of genes in T cell activation in response to pathogens shows significant variations in 348 healthy European, African, and Asian individuals (Ye et al., 2014). To examine whether HS lncRNAs contribute to these variations, we examined whether DBSs in these genes exhibit different binding affinities across populations. We found that DBSs in *IFITM3*, *IL2RA*, *IL17F*, *MXRA7*, *CCL22*, and *FADS2* contain SNPs with biased frequencies in CEU, CHB, and YRI. A typical example is the DBSs in *FADS2,* a gene expressed differentially in Europeans and Africans. The two DBSs in *FADS2-003* and *FADS2-010*, respectively, contain four SNPs - rs71046746 has DAF 0.96/0.99/0.88 in CEU/CHB/YRI, but the other three unannotated SNPs have DAFs 0.24/0.22/0.01, 0.24/0.22/0.04, and 0.24/0.22/0.02 in CEU/CHB/YRI.

Another study examined genome-wide patterns of selection in 230 West Eurasians who lived between 6500 and 300 BC and identified selection signals in multiple genes that are associated with diet, pigmentation, and immunity (Mathieson et al., 2015). We detected signatures of positive selection in DBSs in *LCT*, *TLR1*, *TLR6*, *TLR10*, *SLC45A2*, *SLC22A4*, *MHC*, *ZKSCAN3*, *FADS1*, *FADS2*, *DHCR7*, *GRM5*, *ATXN2*, and *HERC2*. Reliable population-specific selection signals were identified in the DBS of RP11-423H2.3 in *HERC2* (the Tajima's D in CEU/CHB/YRI are -0.19/1.82/-1.12 and the integrated Fst is 0.27), in the DBSs of RP11-423H2.3 in *TLR1* and *TLR6* (the Tajima's D in CEU/CHB/YRI are -1.26/-1.2/1.38 and the integrated Fst is 0.24), and in the DBS of SNORA59B in *TLR1* and *TLR6* (the Tajima's D in CEU/CHB/YRI are -1.73/-0.9/1.76 and the integrated Fst is 0.24). Taken together, the above analyses suggest that population-specific selection signals in DBSs may help explain the phenotypic and physiological differences between human populations.

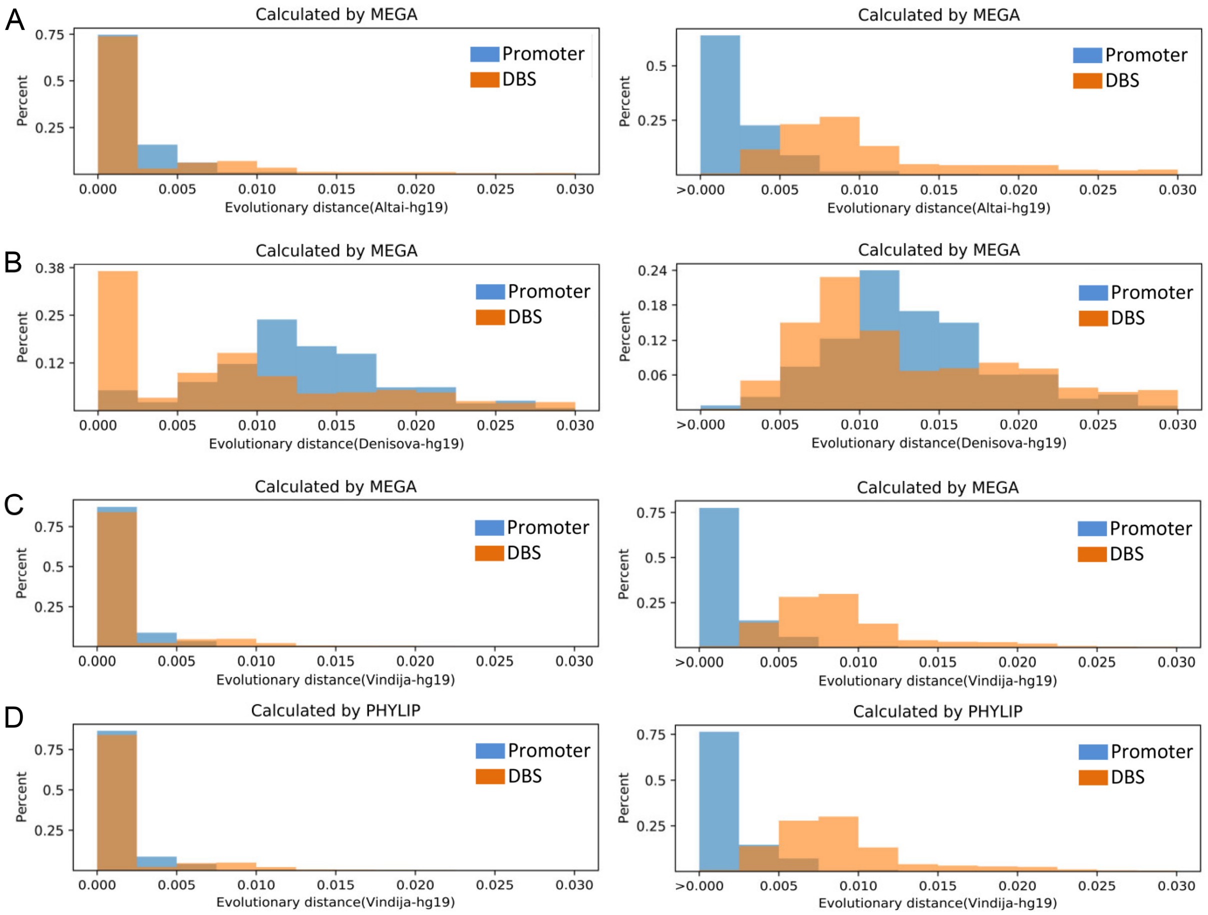


Supplementary Figure 20. The distributions of DBS sequence distances and promoter sequence distances from modern to archaic humans (right-hand panels illustrating distances >0.005). A fraction of DBSs has larger distances than promoters.

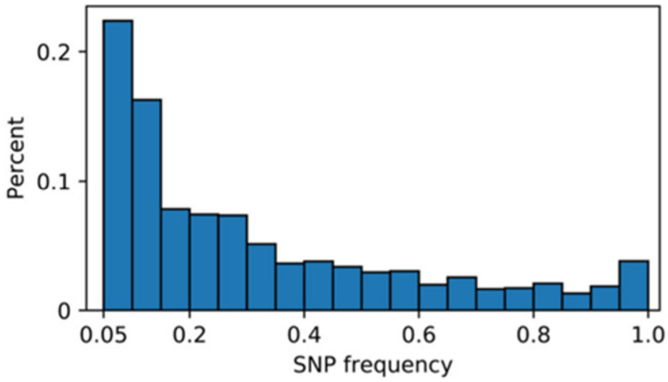


Supplementary Figure 21. The distribution of SNP frequencies (MAF>0.05) in DBSs.

**Supplementary Note 6 – Considerable SNPs in DBSs have a cis-effect on gene expression in specific tissues and populations**

To find supporting evidence for DBS influencing gene expression in tissues and organs, we first examined whether SNPs in DBSs have a cis-effect on gene expression using the data of the Genotype-Tissue Expression (GTEx) project (GTEx Consortium, 2017). The expression of some genes in specific tissues is of interest. For example, of the 21 SNPs that are eQTLs exclusively in the GTEx tissue Thyroid, 14 have high DAF in YRI (Supplementary Figure 22). Correspondingly, in Africans, positive selection signals were found in genes involved in energy metabolism (Fan et al., 2019). Second, we computed the eQTL density for each DBS and each Ensembl-annotated promoter and compared the eQTL density of the two kinds of sequences. We found that eQTLs were more enriched in DBSs than in promoters (one-sided Mann-Whitney test, p = 0.0) (Supplementary Figure 23). Third, for DBS harboring eQTLs in specific tissues, 94% of the corresponding HS lncRNA-target transcript pairs exhibit expression correlation (|Spearman's rho| > 0.3 and FDR < 0.05) in the eQTLs' tissues.

Finally, we examined how many SNPs DBSs are DNA methylation QTL (mQTL) and histone modification QTL (haQTL). One study analyzed DNA methylation data from three populations (10 Caucasians, 10 African Americans, and 10 Japanese) and identified *RGS6*, *CLEC2L*, *ABCF1*, *MIR3678*, and *EP400* as having differential methylation in African Americans compared with Japanese and Caucasians (Giri et al., 2017). Based on this study, we found that DBSs of several HS lncRNAs in some transcripts of *RGS6*, *CLEC2L*, and *EP400* (also *EP400NL*) had significantly different Tajima's D and integrated Fst in YRI. Another study applied QTL analysis to a multi-omics dataset and identified a set of SNPs significantly associated with levels of gene expression, DNA methylation, and histone modification in the human dorsolateral prefrontal cortex (Ng et al., 2017). Ng et al. called SNPs with cis-effect (including eQTL, mQTLs, and haQTLs) xQTL and found that xQTL SNPs were enriched close to transcription start sites. We re-examined these xQTL SNPs and identified 4735 in the DBSs of HS lncRNAs (Supplementary Table 13). Notably, 4319 out of the 4735 xQTLs were meQTL SNPs, many of which had biased frequencies in CEU, CHB, and YRI.

| SNP ID | CEU-frequency | CHB-frequency | YRI-frequency |
|---|---|---|---|
| rs75508216 | 0.01 | 0.05 | 0.1 |
| rs114086993 | 0.01 | 0.05 | 0.1 |
| rs201187971 | 0.01 | 0 | 0.1 |
| rs73677017 | 0 | 0 | 0.14 |
| rs11944829 | 0 | 0 | 0.14 |
| rs114884549 | 0 | 0 | 0.15 |
| rs77133472 | 0 | 0 | 0.15 |
| rs115688283 | 0 | 0 | 0.17 |
| rs113131895 | 0 | 0 | 0.17 |
| rs142522981 | 0 | 0.02 | 0.19 |
| rs112731299 | 0 | 0 | 0.2 |
| rs4565803 | 0.01 | 0 | 0.24 |
| rs4604779 | 0.01 | 0 | 0.24 |
| rs76612433 | 0.02 | 0.19 | 0.24 |

Supplementary Figure 22. The 14 SNPs have high DAF in YRI and are eQTLs exclusively in the GTEx tissue Thyroid.

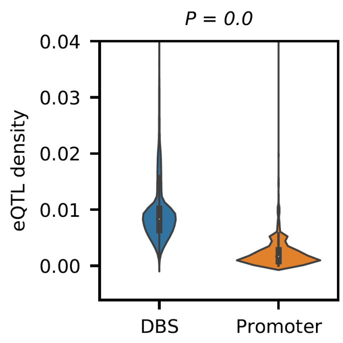


Supplementary Figure 23. DBSs have significantly higher eQTL density than promoters. DBSs and promoters harboring at least one eQTL were used to compute eQTL density and make the comparison. A one-sided Mann-Whitney test was used to compute the p-value.

**Supplementary Note 7 – HS lncRNAs critically regulate genes important for brain development**

The enlarged brain is one of the most prominent features of modern humans. We analyzed multiple experimental datasets to examine the impact of HS lncRNAs on brain development. First, we analyzed two datasets of epigenetic studies. A comparative H3K27ac and H3K4me2 profiling of human, macaque, and mouse corticogenesis revealed that strongly concordant epigenetic gains are enriched in promoters of 301 genes that are active during human corticogenesis (Reilly et al., 2015). Of these genes (257 are annotated in hg38), 84% have DBSs for at least one HS lncRNA, and 56% have DBSs for >= 5 HS lncRNAs. Thus, these 301 genes are enriched significantly for regulation by HS lncRNAs compared with genes without DBSs of HS lncRNAs (p = 1.32e-26 for the situation of 56% genes, and p = 1.21e-21 for the situation of 84% genes, two-sided Fisher's exact test). Another study examined gene expression and H3K27ac modification in eight brain regions in humans and four other primates and revealed 1851 genes (1687 genes annotated in hg38) with human-specific transcriptome differences and 240 genes with chimpanzee-specific transcriptome differences in at least one brain region (Xu et al., 2018). Of these genes, 73% have DBSs for at least one HS lncRNA, and 46% have DBSs for >=5 HS lncRNA. These data again indicate that genes related to brain development are significantly enriched in HS lncRNA targets (p=1.1e-75 for the situation of 46% genes and p=1.2e-56 for the situation of 73% genes, two-sided Fisher's exact test).

Second, we analyzed the data from two studies conducted by the PsychENCODE consortium. The data from one study cover 16 brain regions and 9 time windows and include spatiotemporally differentially expressed genes, spatiotemporally differentially methylated sites, spatiotemporal variations in H3K27ac enrichment, and cell-type marker genes (Li et al., 2018). Of the 66 HS lncRNAs, 65 were expressed in all of the 16 brain regions and in the 9 time windows, and 7 HS lncRNAs (RP11-423H2.3, RP11-706O15.5, RP13-539J13.1, SNORD3B-1, SNORD3B-2, RP4-740C4.7, and RP13-516M14.1) exhibited spatial or temporal expression differences. To assess the contribution of these 7 HS lncRNAs to the spatiotemporal changes of gene expression, we mapped the spatiotemporally differentially methylated sites to the promoters of annotated transcripts and identified 109 transcripts that had these sites in their promoters. We then predicted the DBSs of the 7 HS lncRNAs in the promoter regions of the 109 transcripts and found that 56 transcripts had DBSs for at least one of the 7 HS lncRNAs. In addition, 61% of 770 cell type-specific marker genes had at least one DBS of the s7 HS lncRNAs. These results suggest that HS lncRNAs regulate spatiotemporal gene expression and determine cell fate during brain development, probably by regulating DNA methylation in promoter regions. The other PsychENCODE study examined the spatiotemporal transcriptomic divergence across human and macaque brain development (Zhu et al., 2018). We examined the potential relationships between the HS lncRNAs and the 8951 genes showing differential expression between human and macaque brains during brain development. We identified DBSs of at least one HS lncRNAs in the promoter regions of 72% of differentially expressed genes in the human brain. Moreover, DBSs of at least 5 HS lncRNAs were identified in the promoter regions of 44% of differentially expressed genes. Compared with genes without DBS for HS lncRNAs, these differentially expressed genes are significantly enriched for regulation by HS lncRNAs (p = 0, Chi-square test). Of the 7 transcription factor genes differentially expressed between humans and macaques, four had DBSs of HS lncRNAs. These results support that gene expression in the human brain is highly regulated by HS lncRNAs.

Third, three recent studies identified several genes, including *NOTCH2NL* and *Aspm*, which are critical for regulating cortical expansion in the human brain (Florio et al., 2018; Johnson et al., 2018; Suzuki et al., 2018). *NOTCH2NL* is highly expressed in radial glia and activates Notch signaling to promote the clonal expansion of human cortical progenitors (Florio et al., 2018; Suzuki et al., 2018). *Aspm* is expressed in mice but shows no contribution to mouse corticogenesis. Nevertheless, an *Aspm* knockout greatly influenced corticogenesis in ferrets (Johnson et al., 2018). We examined whether HS lncRNAs potentially regulate these genes. Of the 40 protein-coding genes reported by the three studies, 14 have DBSs of >= 5 HS lncRNAs, and 29 have DBSs of >= 1 HS lncRNAs in their promoter regions (Supplementary Table 14). Compared with the background situation (22562 protein-coding genes' promoter regions contain, but 20944 protein-coding genes' promoter regions do not contain, DBSs of HS lncRNAs), the 40 protein-coding genes involved in cortical expansion are highly enriched for regulation by HS lncRNAs ($p < 0.01$, two-sided Fisher's exact test).

Recently, brain organoids have emerged as an important approach to studying primate neural development in vitro. By establishing and comparing cerebral organoids between humans, chimpanzees, and macaques, Pollen et al. identified 261 human-specific gene expression changes (Pollen et al., 2019). In another study, Agoglia et al. generated a panel of tetraploid human-chimpanzee hybrid iPS cells (i.e. hybrid induced pluripotent stem cells) by fusing human and chimpanzee iPS cells and differentiated the hybrid iPS cells into hybrid cortical spheroids (Agoglia et al., 2021). By allele-specific expression (ASE) analysis, Agoglia et al. identified thousands of genes with divergent expressions between humans and chimpanzees. We obtained the 261 genes from the first study and the 1102 genes with |logFC| > 1 and adjusted $p < 0.05$ from the second study. Compared with the background situation mentioned above, the two sets of genes were significantly enriched with DBSs of HS lncRNAs (p = 1.2e-16 and 3.4e-74, respectively).

**Supplementary Note 8 – The analysis of HS TFs and their DBSs**

We lastly examined the contribution of HS TFs to gene expression in GTEx tissues and organs. Kirilenko et al recently identified orthologous genes in hundreds of placental mammals and birds, organized genes into pairwise datasets using humans and mice as the references (e.g., "hg38-panTro6", "hg38-mm10", and "mm10-hg38") (Kirilenko et al., 2023). In the hg38-panTro6 dataset, the many2zero and one2zero lists (which contain 0 and 147 genes, respectively) indicate the multiple human genes and the one human gene that have no orthologues in chimpanzees. Two studies and the *SCENIC* package reported three human TF lists (Bahrami et al., 2015; Lambert et al., 2018). Based on these data, we identified FOXO4, ZNF41, ZNF843, ZNF717, and TIMM8A as HS TFs (but note that not all of these orthologues are compatible with the Ensembl-annotated ones).

Multiple programs have been developed to predict TF DBSs, which has been a challenging task because TFs have complex 3D structures, TF DBSs are very short (typically around 10 bp), and TF-DNA binding involves diverse co-factors (Bianchi et al., 2025). Two popular programs are *FIMO* and *CellOracle* (Grant et al., 2011; Kamimoto et al., 2023). For each DBS, *FIMO* reports the start and end coordinates, but *CellOracle* reports just a position. First, for the above 5 potential HS TFs, we used *FIMO* and *CellOracle* to predict their DBSs in the 5000 bp promoter regions of the 179128 Ensembl-annotated transcripts (release 79). Since TF DBSs predicted by *FIMO* have been incorporated into the JASPAR database (Rauluseviciute et al., 2024), we directly extracted the related DBSs from the "JASPAR Transcription Factor" track in the UCSC Genome Browser (GRCh38/hg38) with the default cutoff. For *CellOracle*, we used the author-recommended score cutoff of 10 and extended 5 bp on both sides from each reported position. The intersection of *FIMO* and *CellOracle* results includes 171208 DBSs of FOXO4 and ZNF41 (Supplementary Table 16). Second, we identified counterparts of HS TF DBSs in archaic humans and chimpanzees and computed sequence distances of these DBSs from modern humans to archaic humans and chimpanzees (as we did for HS lncRNA DBS analyses). A small portion of HS TF DBSs lack chimpanzee counterparts, and we assumed their distance=10 (as we did for HS lncRNA DBSs). Since detecting selection signals in sequences as short as 10 bp is challenging, we simply used the DeepFavored system to scan HS lncRNA DBSs and HS TF DBSs and compared the results (Tang et al., 2022). The number of favored mutations is about 4 times higher in HS lncRNA DBSs than in HS TF DBSs (the number / the total length). Third, we computed the Pearson correlation between HS TFs and their target transcripts in GTEx tissues to identify correlated expression (|Spearman rho|>0.3 and FDR<0.05). Of all HS TF-target transcript pairs, 95% show correlated expression in at least one tissue. However, unlike the high percentages of correlated HS lncRNA-target transcript pairs in the brain, correlated HS TF-target transcript pairs are distributed across many tissues and organs (Supplementary Figure 24). Finally, for each DBS in an HS TF-target transcript pair that shows correlated expression in a GTEx tissue, we computed the sequence distances of this DBS from modern humans to the three archaic humans, and then compared the distribution of DBS sequence distances in each tissue with the distribution of DBS sequence distances in all tissues (one-sided two-sample Kolmogorov-Smirnov test). Unlike DBSs in HS lncRNA-target transcript pairs (Figure 3), the significantly changed DBSs (in terms of sequence distance) in HS TF-target transcript pairs across GTEx tissues and organs do not show dense distribution in the brain (Supplementary Figure 25). Taken together, these results suggest that HS lncRNAs may have contributed more significantly to human evolution than HS TFs by regulating gene expression.

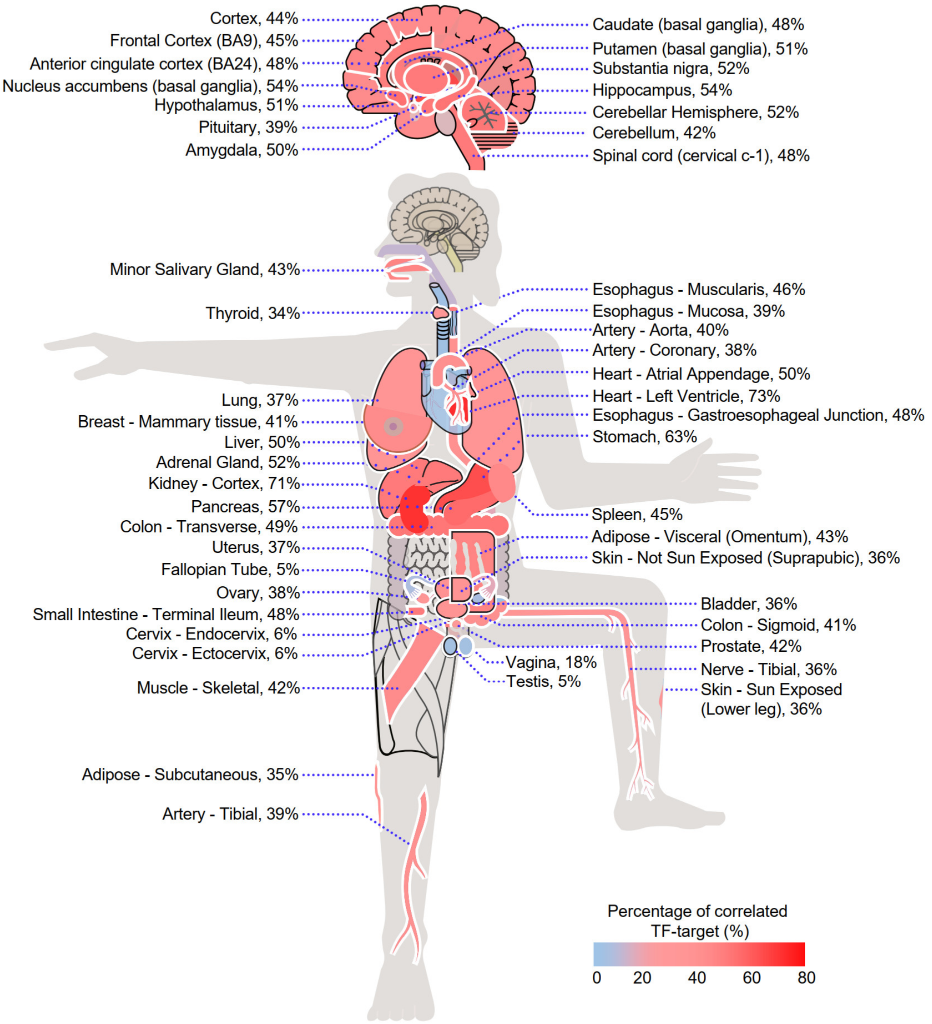


Supplementary Figure 24. The distribution of the percentage of HS TF-target transcript pairs with correlated expression across GTEx tissues and organs (see Figure 3A).

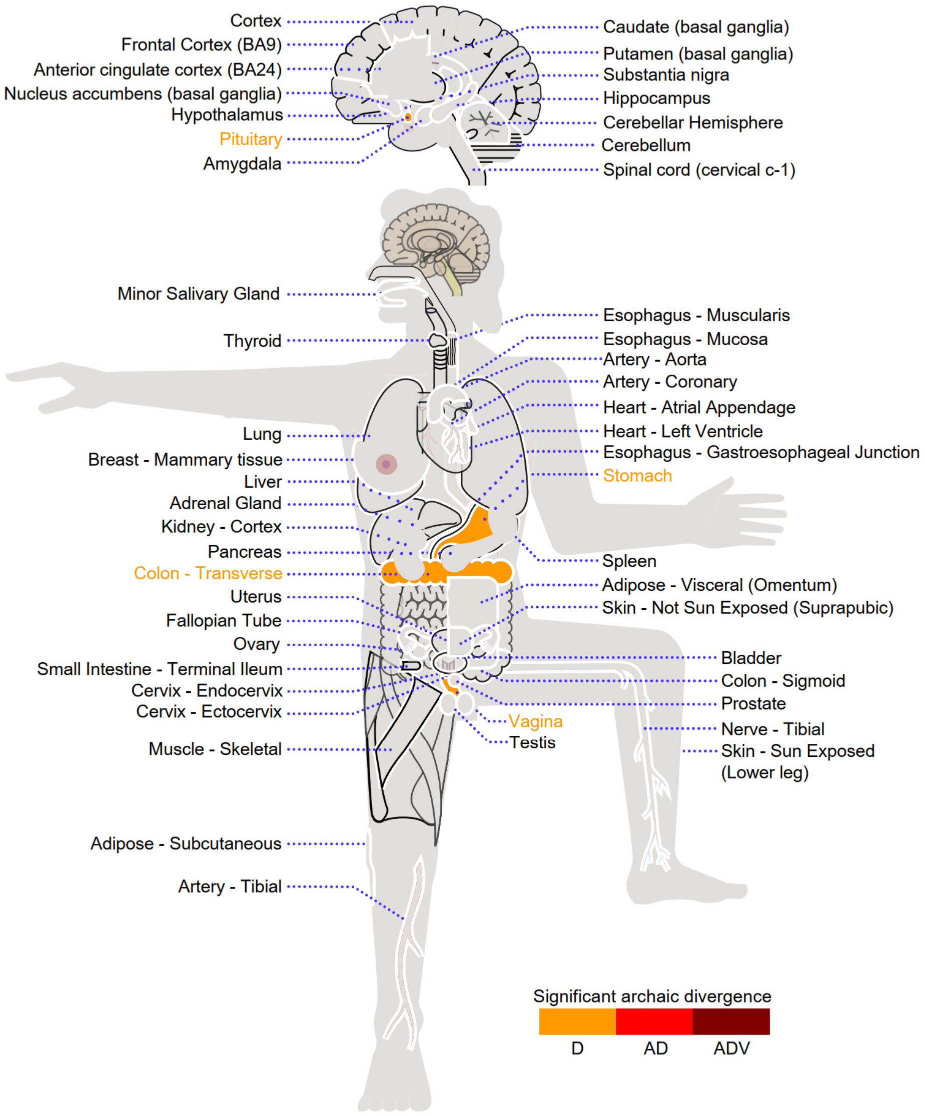


Supplementary Figure 25. The distribution of significantly changed DBSs (in terms of sequence distance) in HS TF-target transcript pairs across GTEx tissues and organs between archaic and modern humans. As in Figure 3B, orange, red, and dark red indicate significant changes from Denisovans (D), Altai Neanderthals and Denisovans (AD), and all three archaic humans (ADV).

### Supplementary Note 9 – Human-specifically rewiring of gene expression in the brain

Human-specific rewiring of gene expression should result in distinct correlations. To identify whether the pattern exists in the brain, we analyzed the transcriptomic data from two brain regions - frontal cortex (BA9) and anterior cingulate cortex (BA24) – in the human and macaque brain (Consortium et al., 2017; Zhu et al., 2018) (n=101 and 83, and n=22 and 25, respectively). We used the *eGRAM* to identify modules of genes with expression correlation and regulatory relationship (Supplementary Figure 26). Orthologous genes (displayed at the same positions in each panel) show much more correlations in humans than in macaques and are significantly enriched for genes in neurodevelopment-related KEGG pathways. This result supports that gene expression is substantially rewired in the human brain by HS lncRNAs compared with in the macaque brain, and the correlations and modules provide novel information for interpreting the mechanisms of human cognition and behavior.

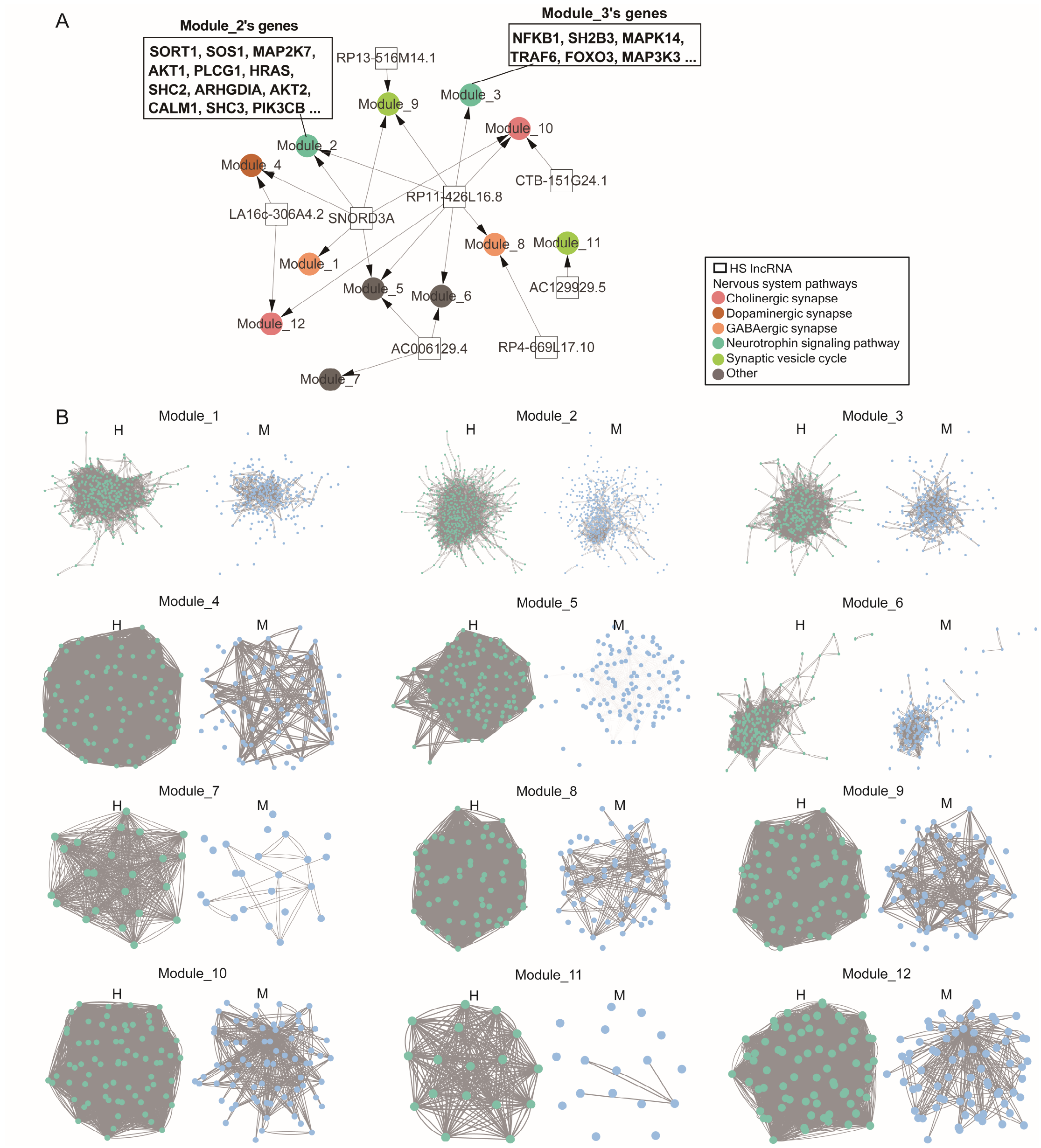


Supplementary Figure 26. Human-specifically rewired gene expression by HS lncRNAs in the anterior cingulate cortex (BA24). (A) Genes expressed in the anterior cingulate cortex are enriched for HS lncRNAs' target genes and neurodevelopment-related pathways. Squares, dots, and colors indicate HS lncRNAs, gene modules, and enriched KEGG pathways, respectively. (B) Comparison of modules and genes in humans (indicated by H) and macaques (indicated by M).
